# Pareto Optimality Reveals an Atlas of Cellular Archetypes

**DOI:** 10.1101/2024.12.04.626890

**Authors:** George Crowley, Uri Alon, Stephen R. Quake

## Abstract

We sought to discover universal organizing principles behind phenotypic variation within cell types. Pareto optimality describes how trade-offs between optimal solutions account for variation, predicting that the boundary points of a data distribution reflect specialized functions. We hypothesized that Pareto optimality dominates transcriptomic variation across all cell types. We used the Tabula Sapiens atlas of single-cell RNA sequencing across cell types and tissues in the human body to test this hypothesis and discovered that most cell types adhere to this theory. This enabled us to use this principled method to characterize the functions performed by each cell type. These phenotypes are derived from an unbiased approach and do not incorporate ideas from existing biological models or theories, and yet in many cases they recapitulate our understanding of the functions of major cell types. Ultimately, we conclude that multi-objective optimization broadly shapes the observed phenotypic variation within cell types. This finding enables us to write explicit representations of the low-dimensional manifolds on which transcriptomes of single cells reside. This can inform the design of the next generation of virtual cell language models, which aim to statistically learn low-dimensional transcriptomic manifolds.

## Main

Advances in single-cell RNA sequencing have made it possible to perform unbiased, high-throughput analysis of single-cell transcriptomes from entire organisms. Recently, the Tabula Sapiens Atlas has sequenced over one million cells across 28 organs of the human body, with tissue replicates across donors.(*1*) Similar projects have produced parallel databases for both human and a variety of model organisms(*2-4*), and substantial efforts have been made to compile these datasets.(*4-6*) These atlases have allowed us to understand cellular heterogeneity across the entire human body, and their analysis has yielded vast stores of rich insight across myriad cell types—both rare and common, including the immune(*7-9*) and stromal(*10, 11*) compartments. Major sources of variation within cell types have been discovered, including cell cycle phase, responses to extrinsic stimuli, and epigenetic modulation to name a few. Despite these advances, it is unclear if there are universal organizing principles that give rise to these numerous sources of phenotypic variation within a given cell type. One natural place to look for such an organizing principle is multi-objective optimization. Pareto optimality is the baseline formulation of multi-objective optimization, and describes the situation where no explicit preference or weighting among the objectives is assumed. This concept is treated mathematically in the study of multi-objective optimization(*12, 13*), and separately applied to biology(*14, 15*).

We hypothesized that multi-objective optimization dominates the phenotypic variation within cell types and used Tabula Sapiens to test this hypothesis. The Tabula Sapiens Atlas v1 is a single-cell RNA sequencing dataset of 456,101 high-quality single cell transcriptomes, covering 58,870 genes across 174 cell types, 25 tissues, and 15 donors.(*16*) We applied quality control filters to remove outlier cells on several metrics, yielding 309,193 cells across 173 cell types, 24 tissues, and 14 donors, **Table S1** and **Figure S1**. Cell type abundance filters left 110 cell types across the same number of tissues and donors, yielding 440 distinct cell type-tissue-donor strata for analysis.(*15, 17*)

The only assumption we make in this analysis is that fitness is an increasing function of performance.(*14*) Then, if there is a trade-off in performing multiple tasks, optimal phenotypes (i.e. those that maximize fitness) must lie in a region described by convex combinations of points that each maximize a single task’s performance.(*14*) This region is called the Pareto front. *Any* pruning mechanism that removes non-optimal phenotypes would restrict observed phenotypes to the Pareto front; pruning is a pervasive strategy across biology, and there could be a host of pruning mechanisms in multicellular organisms.

This approach does not require any assumptions about underlying regulatory dynamics or interactions among units. The Pareto front simply describes the region of optimal phenotypes, and its vertices are phenotypes each optimal at some task. Etiology and underlying regulatory dynamics can shape the Pareto front, but do not contradict that optimal phenotypes must lie on it.(*18*) The elegance and power of Pareto optimality are that no specific selection mechanism or regulatory dynamics are required to arrive at its conclusions.

Phenotypes could fill the Pareto front or occupy subregions of it, depending on the form of the fitness function and externalities including but not limited to physical or biochemical constraints, or spatiotemporal variation in the environment.(*19, 20*) Hence, only the two following requirements must be met to conclude that Pareto optimality dominates: 1) phenotypes are bounded by a polytope in trait-space; and 2) the vertices of the polytope have enriched features.(*17*) This conclusion can be further strengthened by identifying specific tasks associated with the enriched features, and showing the tasks’ recurrence across independent contexts.(*17*)

Therefore, our analysis aimed to determine whether the Tabula Sapiens data are well described by polytopes, and whether the vertices of those polytopes have significantly enriched features. In finding that both conditions are indeed satisfied, we then found further support by showing that the vertices are functionally relevant based on our prior understanding of cell biology.

These findings lead to the conclusion that multi-objective optimization broadly shapes phenotypic variation within cell types.(*17*) This framework then allows us to infer which tasks of a given cell type may share a common biochemical environment.

### Most cell types are described by polytopes

We performed dimensional reduction and model fitting independently for each distinct cell type-tissue-donor dataset, **Figure 1A**. We used this stratification to ensure that variation due to donor, tissue or cell type did not define the space modelled, and instead only variation within a donor-tissue-cell type was captured. After removing mitochondrial genes and genes known to be affected by tissue processing, we normalized the remaining genes to 10,000 counts (so that cell size effects would not dominate our results), and used principal component analysis (PCA) of protein-coding genes to represent cells in PCA space.(*16*)

**Figure 1.**
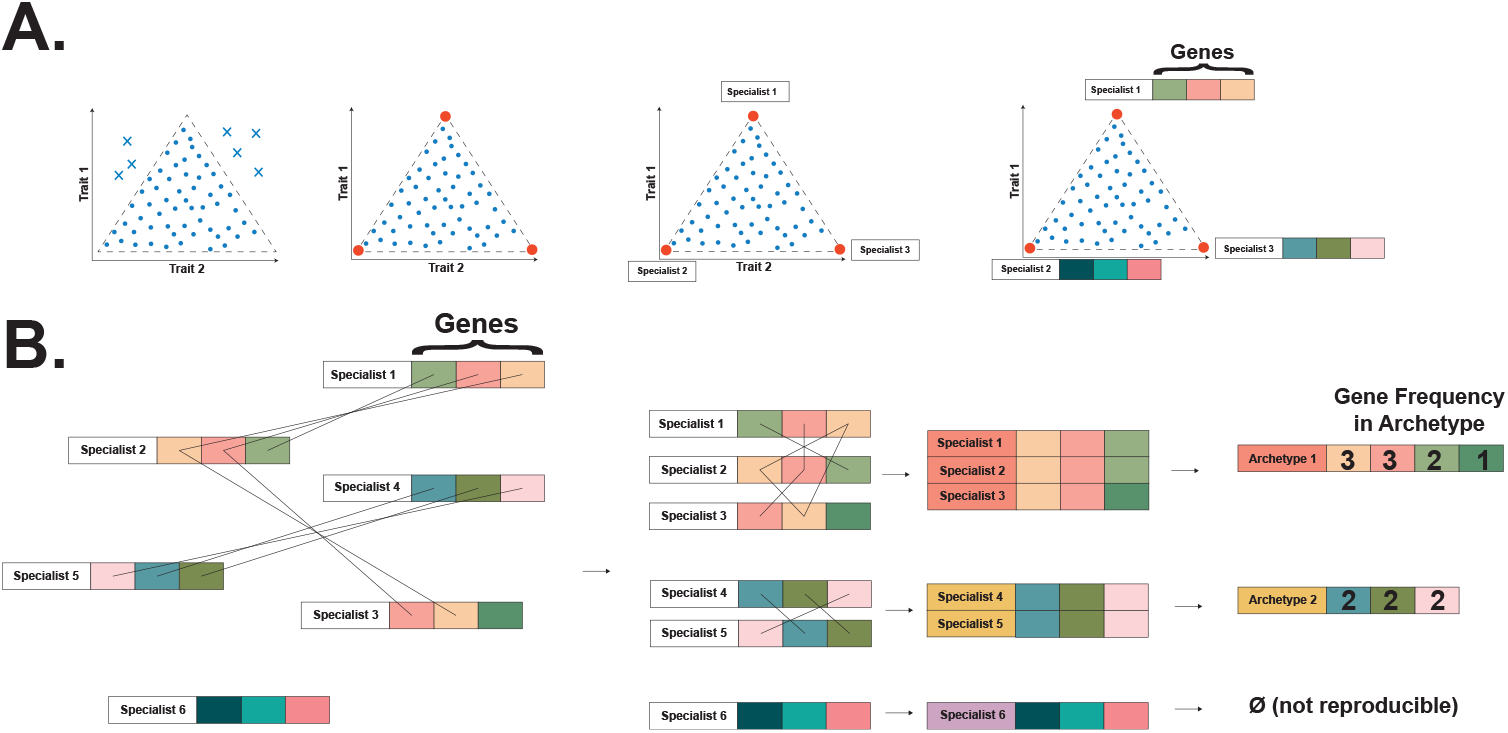
Process Overview. A. Pareto Optimality Theory. Pareto optimality places limits on allowable phenotypes. In trait space, this constrains phenotypes to be bound by a polytope. Specialist phenotypes lie at the vertices of the polytope and are enriched in traits relevant to their specialist functions. **B. Archetype Alignment**. Vertices are clustered on a per-cell-type basis, yielding clusters that are filtered for reproducibility across donors and tissues.

Several common steps in single-cell transcriptomics analysis are potentially incompatible with the analysis required to assess the dominance of Pareto optimality. Instead of subselecting only highly variable genes, we retained all protein-coding genes for analysis because we did not want to introduce unknown bias into analyzed phenotypes, and it is also possible that non-highly variable genes could still be enriched in cell populations near the archetypes.

Separately, data transformations are important to the space modelled in single cell transcriptomics. We only normalized the raw, protein-coding gene counts. Because the geometry of single-cell-type expression may be linear and the polytope vertices may be related to physical distances, we avoided common single-cell transformations that do not preserve pairwise distances, such as logging, scaling, and neighborhood-graph analysis.(*19*) Finally, a density-based filter removed outliers in PCA space, stabilizing subsequent PCA transformations performed on linear-scale expression data.(*21*) The use of this filter in archetypal analysis dates back over 30 years to the original archetypal analysis paper by Cutler and Breiman (1994).(*22*)

This experimental design precludes us from identifying rare cell populations by aggregating cells across donors or tissues. Thus, the density-based outlier filter is a conservative way to limit our analysis to the major cell populations, and try to achieve a more complete understanding of the variation of function just in these major populations. This is in-line with the goal of our investigation, which is to test if multi-objective optimization broadly shapes phenotypic variation within each cell type.

Due to a previous suggestion in the literature that systematic technical artifacts could produce polytopal structures(*23*), we excluded artifactual genes that have previously been described to be affected by the tissue dissociation process (*16*) (linked here for convenience) and mitochondrial genes (the 37 genes of the mitochondrial genome) before PCA to ensure they did not define the space. Following PCA, we dropped components that were correlated with artifactual or mitochondrial expression above a stringent threshold of |r|>0.3. To further dispel concerns that polytopes were related to technical artifact, these genes were still included in the enrichment analyses, and, as later elaborated, we observed no pattern of their enrichment at the vertices or influence on the fits. This gives us reason to believe that our observations in this space are driven by biological variation.

All data preprocessing steps are described in detail in the Methods.

We performed polytope fitting on the single-cell transcriptomes of each donor-tissue-cell type, **Figure 2**, with confidence bounds determined by bootstrapping, and p-values determined by the t-ratio test. The t-ratio measures the data’s similarity to a polytope.(*15, 17*) The t-ratio test was performed by comparing the goodness of fits of shuffled and unshuffled data. The p-value was the proportion of times that the goodness of fit of shuffled data was better than that of the unshuffled data over 1,000 runs.(*17*) To get properly calibrated p-values, the PCs are shuffled and not the genes. Hart et al. 2015 (*17*) provide an in-depth discussion of this test, and (*23*) shows it to be the more stringent of two procedures tested.

**Figure 2.**
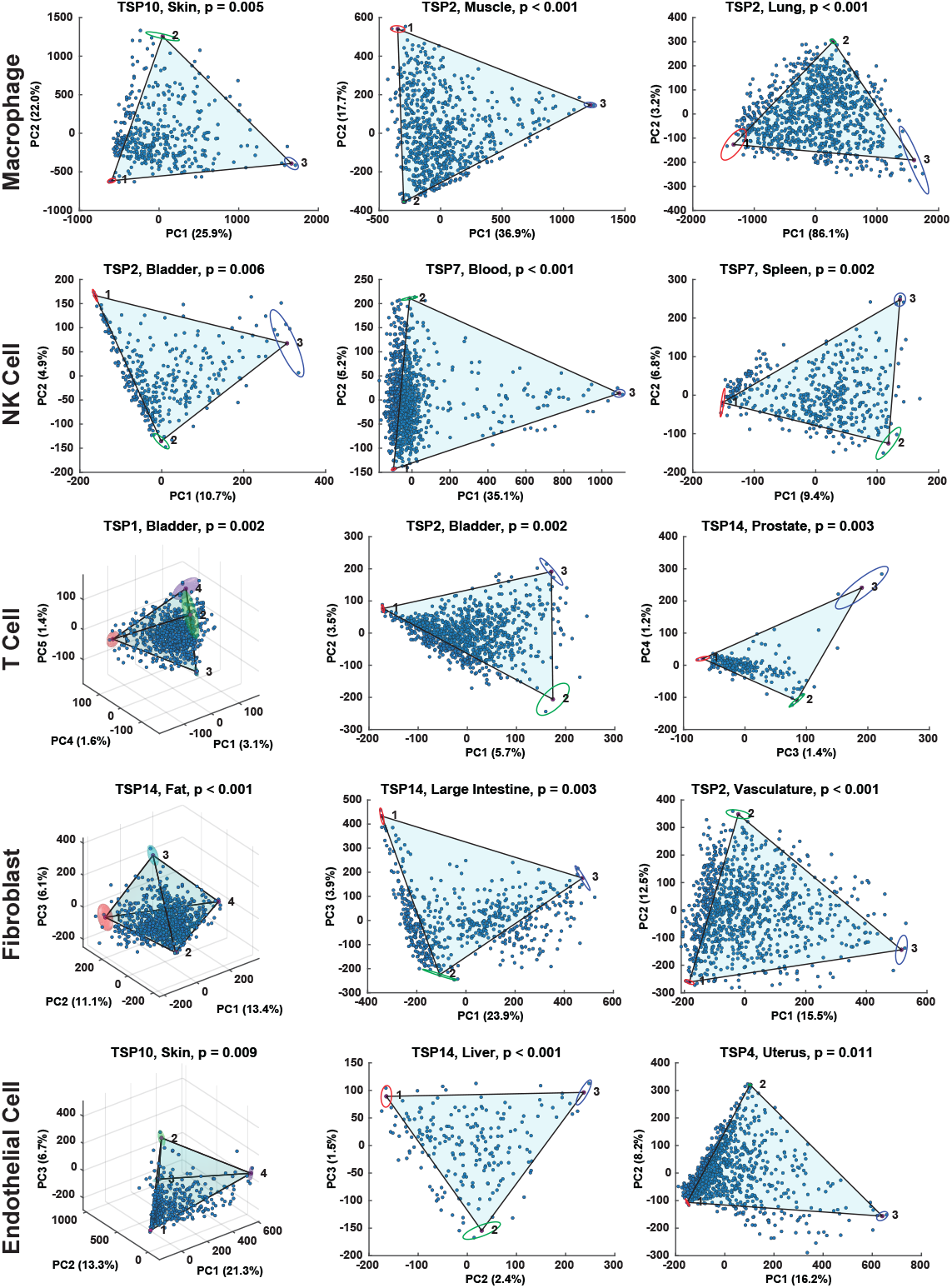
Reproducible Geometry of Donor-Tissue-Cell Types and Fits. Principal components analysis and Pareto Task Inference analysis performed independently on macrophages, NK cells, T cells, fibroblasts, and endothelial cells across 3 unique donor-tissues each demonstrates this geometry. Confidence bounds on vertex position are shown by ellipses and ellipsoids.

We set the significance threshold α = 0.05 for assessing significance of fit of a polytope to a given donor-tissue-cell type, and accepted a maximum false discovery rate (FDR) = 0.10 for determining which cell types were significantly fit overall. We labeled a cell type as significantly fit by polytopes overall if ≥ 50% of donors with that cell type available had significant tissues.

Three-quarters of cell types (82/110, 75%) were significantly well fitted by polytopes (defined by ≥ 50% of donors having significant tissues), **Figure 3A, B, C**. The associated false discovery rate is 0.0849, **Figure 3D**, which corresponds to roughly 7 false positives out of the 82 significantly fit cell types. Independently, a majority (247/440) of donor-tissue-cell type strata were each significantly fitted by polytopes. Furthermore, when removing singlet and doublet cell types (i.e. cell types with only one or two donor-tissues), 90% of cell types (35/39) were significantly fitted. We observed no substantial relationships between quality control metrics—including the percent of counts coming from artifactual genes—and significance of fits, **Figure S2**.

**Figure 3.**
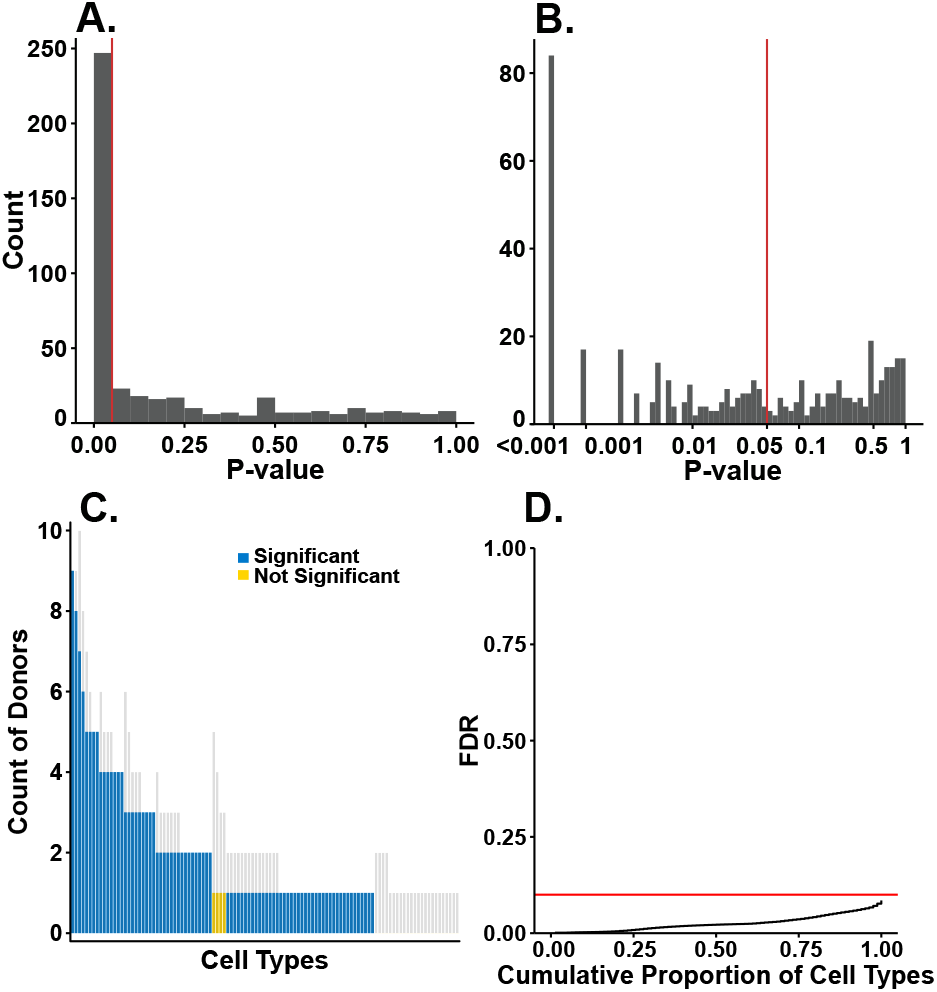
Most cell types are significantly well fitted by polytopes. Significance of fits of each donor-tissue-cell type in Tabula Sapiens (both **A**. standard and **B**. log-scaled axes shown). **C**. Counts of donors with significant tissue(s) present in each cell type, plotted against a background of all donors available for that cell type (grey background bars). Cell types with significance in at least 50% of available donors are considered significant. **D**. Cumulative distribution function of false discovery rate (FDR) as a function of proportion of significant cell types retained. Three quarters of available cell types are significantly fit at α=0.05 and maximum FDR=0.10. Vertical red lines are drawn at p=0.05 (A. and B.) and FDR=0.10 (C.).

A detailed analysis of the frequency of significance of cell types shows that each donor and tissue has significantly fitted cell types, except for TSP3 (only a single donor-tissue-cell type available) and kidney (only 2 donor-tissue-cell types available), **Figure S3**. We have thus substantiated the first of the two requirements to show that Pareto optimality dominates the phenotypic variation—the phenotypes are bounded by polytopes in trait-space.

What remains is then to show that the vertices of these polytopes have enriched features. If the polytopes arose due to the structure of statistical noise present in the data, we would not expect to observe features that were significantly enriched in the cells nearest to the vertices.(*17*) Across the 247 significant polytope fits of donor-tissue cell types, there were a total of 864 vertices, all of which had genes that were significantly enriched in the nearest cells as measured by Benjamini-Hochberg-corrected p-values with a maximum false discovery rate of 0.10, **Figure 4**. This dismisses the concern that the polytope data structure arose from statistical noise, and substantiates the second of the two requirements to show that Pareto optimality dominates: the vertices of the phenotype-bounding polytopes have enriched features. At this point, both the necessary and sufficient conditions have been satisfied to conclude that Pareto optimality drives phenotypic variation within most cell types in the human body.

**Figure 4.**
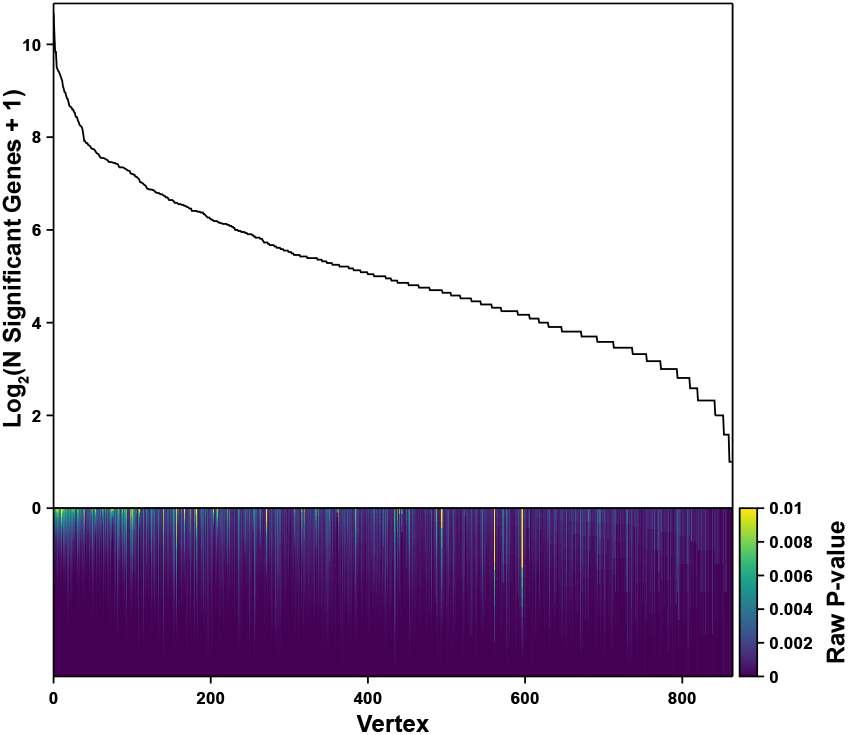
All vertices of significantly well fitted polytopes have significantly enriched genes. The number of significantly enriched genes in each of the 864 vertices from the 237 significant polytope fits, plotted on log scale. Below, a stacked bar chart, with each segment representing a significantly enriched gene and colored by that gene’s p-value. Note that the range of the color bar goes from p=0 to p=0.01.

### Polytope vertices reflect physiological functions

We next sought to identify specific tasks associated with the vertices and test whether those tasks recurred across independent contexts. Importantly, in Tabula Sapiens, there are 52 independent datasets (each donor-tissue that has significant cell types) across which to evaluate consistency.

The tissues with the highest number of donor-cell type pairs available were blood, spleen, and lung, **Figure S4**. The donors with the highest number of tissue-cell type pairs available were TSP14, TSP2, and TSP7, **Figure S5**. Finally, the cell types with the highest number of donor-tissue pairs available were macrophages, natural killer (NK) cells, T cells, fibroblasts, and endothelial cells, **Figures S4, S5**. We therefore focused on these cell types across tissues to understand the correspondence of polytope vertices to biological functions.

Each of the 864 vertices potentially represents a biological function. In order to make headway, we focused on the most commonly observed vertices within the most abundant cell types.

Thus, the remainder of our results focuses on broad strokes of the most confidently identified functions. There are many hundreds more vertices that could correspond to specialized cell functions that are tissue- or even donor-specific that we do not discuss here. We calculated gene enrichment at each vertex, and used the set of all protein-coding genes, including mitochondrial and artifact genes to allow the detection of artifactual vertices.

We restricted our functional inference to 864 vertices from the 35 cell types that were significantly fit overall and had more than two donor-tissues. Within each cell type, we then clustered vertices based on up to their ten most highly differentially expressed genes, and further narrowed our focus to clusters that contained vertices from more than two-thirds of available donors for that cell type, with a correction factor to control false negatives. This set of filters is highly conservative and restricts us to identifying only the most common, broad strokes of cell function. Therefore, while there is rich subtype literature on cell types that we discuss, we focus on the dominating expression patterns per cell-type. It is worth noting that for each cell type there are many vertices that we do not plot or discuss that could correspond to niche functions. However, even given these restrictions, we will see below that this principled analytical approach still identifies universal cell functions that remain undercharacterized by current cell subtype schema.

For functional inference, we considered the genes that were shared among the vertices within a cluster. We used a large language model to provide a broad annotation for each gene list.

Similar LLM approaches have been benchmarked recently and showed satisfactory agreement with gene ontology.(*24, 25*) We used Gene Ontology-Biological Process, and several commercially available LLMs (see methods); Claude 4 Sonnet generally gave the most interpretable annotations. Note that these LLM-generated labels were not designed to provide detailed biological insight and we subsequently perform detailed manual comparison to established cell subtype repertoires.

### Archetypes have distinct expressional programs

We calculated the average gene expression of the defining genes of each archetype. We then compared the gene expression of archetype-defining genes within their own archetype to the expression of these genes in other archetypes. In all cell types discussed—macrophages, NK cells, T cells, fibroblasts, and endothelial cells—archetypes were defined by unique expressional programs; we conclude that the defining genes of each archetype are highly expressed within their own archetype, and lowly expressed in other archetypes, **Figure 5A-E** (second row). Thus, the archetypes are phenotypically distinct from each other.

**Figure 5.**
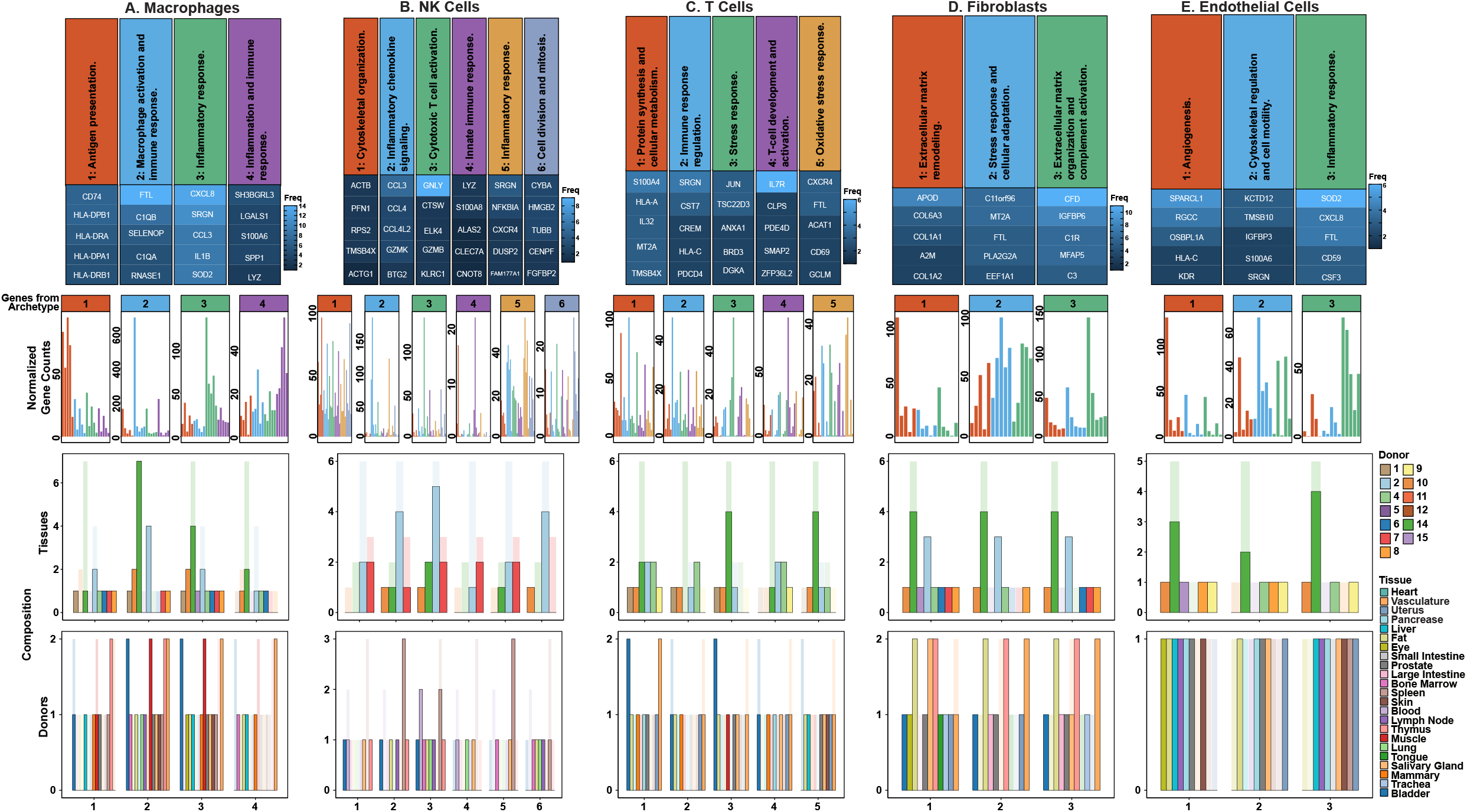
Archetypes of Ubiquitous Cell Types. For 5 ubiquitous cell types (**A. Macrophages**; **B. NK Cells**; **C. T Cells**; **D. Fibroblasts**; and **E. Endothelial Cells**), we show: 1^st^ row: Heatmap of the top 5 most frequent archetype-defining genes in archetypes containing approximately two-thirds or more of available donors, annotated via a large language model; 2^nd^ row: Gene expression of top 5 genes of each archetype, visualized across all archetypes, using the 5% of cells closest to each vertex on a donor-tissue-cell type basis. Within each archetype, genes are plotted in the descending order of appearance in the heatmap in the 1^st^ row; 3^rd^ row: Counts of tissues present in each donor for each archetype (opaque bars), plotted against a background of all tissues available for that donor-cell type (semi-transparent bars); 4^th^ row: Counts of donors present in each tissue for each archetype (opaque bars), plotted against a background of all donors available for that tissue-cell type (semi-transparent bars)

### Macrophage archetypes reflect immune, metabolic, tissue homeostatic roles

Macrophages are multitasking cells found in all tissues. Historic approaches subdivide macrophages into discrete categories such as M1 and M2 states. It has been argued that this taxonomy should be abandoned in favor of a function-based nomenclature.(*26*) More recently macrophages have been divided into discrete eight substates, although data shows that macrophage phenotype is continuous in gene expression space.(*8*) Here, we applied a principled method to extract these functions without discretization and define them based on molecular expression. We find that macrophages are significantly fit by polytopes across tissues. Many of the archetype defining genes for macrophages have had specific tasks associated with them in prior literature. Here, we show how sets of these tasks may be consolidated under shared biochemical phenotypes.

The first macrophage archetype describes macrophages’ role in antigen presentation via the Major Histocompatibility Complex (MHC) class II pathway (with enrichment in *CD74, HLA-DRA, HLA-DPB1, HLA-DPA1, HLA-DRB1, HLA-DQA1, HLA-DRB5*), **Figure 5A**.(*27*) This archetype also shows macrophage roles in activation and amplification of the complement system (with enrichment in *C1QA, C1QB, C1QC, C3*) and homeostatic regulation of these processes (*CST3*).(*28*) Antigen presentation and complement activation are known to be expressed in macrophages across tissues and are key to core macrophage function and immunity.(*29*) Our analysis indicates that these two functions may be performed by transcriptionally similar macrophage cells. This sheds light on an open question, which is to elucidate the transcriptional programs active in these pan-tissue functions, and assess the degree to which they are shared or unique.(*29*)

The second macrophage archetype reflects macrophages specialized for long-term tissue residence and metabolic coordination.(*30*) This population is characterized by iron homeostasis and storage (with enrichment in *FTL, FTH1*),(*31*) complement-mediated clearance functions (*C1QA, C1QB, C1QC*),(*32*) and antioxidant defense and tissue maintenance through Selenoprotein P (*SELENOP*)(*33*). Additional features include general RNA turnover and antimicrobial activity (*RNASE1*), cysteine protease inhibitors (*CSTB, CST3*), lipid metabolism regulation (*PLTP, APOE, APOC1*)(*34*), scavenger receptor-mediated clearance (*MARCO, MRC1*), and anti-inflammatory regulation (*VSIG4*)(*35*). This archetype represents tissue-resident macrophages that maintain tissue homeostasis through metal and lipid metabolic coordination and controlled immune surveillance.

The third macrophage archetype reflects macrophages’ role in the inflammatory response and was enriched in cytokine and chemokine production (*CXCL8, CCL3, IL1B, CCL4, CCL20, CCL3L1, CCL4L2*) and regulation (*SRGN*), as well as cell migration in response to inflammation (*GPR183*, also known as *EBI2*)(*36*). This population also shows metabolic adaptations for sustained inflammatory activity, including antioxidant defense during oxidative burst (*SOD2*)(*37*), energy metabolism to support high cytokine production (*NAMPT*), and metabolic reprogramming (*G0S2*)(*38*). Additionally, these macrophages exhibit early tissue repair signaling (*EREG*), indicating their role as sophisticated inflammatory coordinators that bridge innate immunity with tissue repair processes.

The fourth macrophage archetype represents highly motile macrophages specialized for acute inflammatory responses and tissue repair initiation. This population is distinguished by damage-associated molecular pattern (DAMP) signaling through S100 protein family members (*S100A4, S100A6, S100A8, S100A9, S100A10*)(*39*), matrix remodeling capacity (*SPP1, VCAN, VIM*), and enhanced cellular motility (*SH3BGRL3, PFN1, VIM*)(*40*). Additional characteristics include inflammation resolution programming (*LGALS1*), proteolytic activity (*CTSD*), and antimicrobial function (*LYZ, VIM*).(*41*) This archetype likely represents recruited macrophages derived from circulating monocytes that rapidly infiltrate inflamed tissues and coordinate both inflammatory responses and early tissue repair processes.

The first and second macrophage archetypes appear to share some functions, namely utilization of the complement system, its regulation, and antimicrobial activity. Indeed, Pareto optimality permits this manner of function sharing. Therefore, the defining function of the first archetype is antigen presentation through the MHC class II pathway, while the second archetype is distinguished by metabolic, tissue homeostatic functions. Complement (C1q) and cystatin (Cystatins B and C) production then seem to be performed without substantial preference by macrophages with either of these two specializations. In other words, antigen-presenting macrophages and those responsible for metabolic homeostasis seem to be similarly well-suited for the production of complement and cystatins.

Meanwhile, the fourth macrophage archetype expresses an additional form of antimicrobial activity via vimentin—a multipurpose protein also involved in cell motility and matrix remodeling. We note that while the fourth macrophage archetype shows an activated phenotype that would be consistent with extracellular secretion of vimentin, we cannot conclude vimentin’s specific fate from transcriptomics alone. However, it is perhaps efficient for this macrophage archetype to build several functions (antimicrobial activity, motility, and matrix remodeling) around the expression of one or more genes like *VIM* that are shared amongst these tasks. Finally, this archetype expresses SH3BGRL3, a protein that has been implicated in cell migration and TNF-α inhibition; This protein remains underexplored, and the present work highlights its importance to physiological macrophage function.

### Natural killer cell archetypes represented motifs of cytotoxicity and immune cell recruitment

NK cells are also found in all tissues and carry out cytotoxic functions. We found that NK cells are well described by a continuum of gene expression, **Figure 5B**. The first NK cell archetype represents specialization in cytoskeletal organization and is characterized by expression of core cytoskeletal machinery including β-actin (*ACTB*), profilin-1 (*PFN1*), and thymosin β4 (*TMSB4X*). Together, these reflect cells specialized for dynamic actin polymerization and cytoskeletal remodeling essential for cell motility and target engagement. Profilin-1 regulates actin nucleation and elongation, while thymosin β4 sequesters actin monomers, thereby halting actin filament assembly.(*42*) The presence of ribosomal protein S2 (*RPS2*) suggests active protein synthesis to support the high metabolic demands of cytoskeletal remodeling. Coordinated cytoskeletal dynamics are fundamental to NK cell function, enabling critical processes including tissue infiltration, granule secretion, immune synapse formation, target cell conjugation.(*43*) This archetype represents NK cells that have been optimized for coordinated cytoskeletal dynamics, thus recapitulating a crucial aspect of NK cell biology.

The second archetype demonstrates specialized inflammatory chemokine production through robust expression of C-C motif chemokine ligands (*CCL3, CCL4, CCL4L2, CCL3L1*). These cells also express granzyme K (*GZMK*), a unique granzyme that activates the all components of the complement cascade.(*44*) This combination suggests NK cells functioning as inflammatory coordinators that eliminate targets while simultaneously recruiting other immune effectors through potent chemokine production. The potential positive feedback between complement and macrophage inflammatory protein signaling lends further biological plausibility to this archetype. This archetype also expresses *CHMP1B*, is a subunit of the ESCRT-III complex, which is thought to contribute to granzyme and perforin resistance.(*45*) Along with expression of BTG anti-proliferation factor 2 (*BTG2*) and heat shock protein A6 (*HSPA6*), this collection of genes could indicate a protective phenotype that allows NK cells to survive in challenging environments when initiating inflammatory signaling and cytotoxicity.

The third archetype represents highly cytotoxic NK cells expressing multiple killing mechanisms, including a potent mediator of cell death, granzyme B (*GZMB*), an antimicrobial peptide, granulysin (*GNLY*), and a cysteine protease, cathepsin W (*CTSW*). These cells express the inhibitory receptor NKG2A (*KLRC1*) and L-selectin (*SELL*), indicating mature NK cells with regulated cytotoxic capacity and tissue homing potential. The ELK4 transcription factor (*ELK4*) suggests active transcriptional programs supporting effector functions. This archetype embodies the classical NK cell cytotoxic program with perforin-granzyme-mediated killing complemented by granulysin’s antimicrobial and cytolytic activities, representing cells specialized for direct target elimination.

The fourth archetype is distinguished by expression of lysozyme (*LYZ*) and S100 calcium-binding protein A8 (*S100A8*), indicating NK cells with enhanced antimicrobial functions. Lysozyme provides direct antimicrobial activity against bacterial cell walls, while S100A8 functions as a DAMP with additional antimicrobial properties. This minimal but functionally coherent gene signature suggests a specialized subset of NK cells that bridges traditional cytotoxic functions with antimicrobial immunity, potentially representing cells adapted for responses against intracellular pathogens or in inflammatory microenvironments where antimicrobial activity is prioritized.

The fifth archetype exhibits elevated *SRGN* for granule formation and degranulation, as well as *CXCR4* for tissue homing. Furthermore, this archetype expresses inflammatory response machinery, centered on transcriptional regulation through immediate early genes (*FOS, NR4A2*) and NF-κB signaling control (*NFKBIA, RPS3A, FAM177A1*), and MAPK downregulation (*DUSP2*). Ribosomal protein 3 (*RPS3A*) acts extraribosomally to regulate NF-κB and is phosphorylated by ERK1, which itself is regulated by *DUSP2*.(*46*) Meanwhile, *FAM177A1* is thought to attenuate the NF-κB cascade; however, its physiological roles—both broadly and specifically in NK cells—remain to be elucidated, and its pathological roles have only recently been studied.(*47*) This archetype also expresses *SLA* (*SLAP*), which is thought to be involved in immune receptor signal transduction(*48, 49*). This archetype represents activated NK cells with transcriptional regulation that enables controlled inflammatory responses and cytokine production.

The sixth archetype captures proliferation through expression of key cell cycle machinery, including centromere protein F (*CENPF*) essential for kinetochore function, stathmin 1 (*STMN1*) for microtubule regulation, and tubulins (*TUBB, TUBA1B*) for mitotic spindle formation. These cells express cytochrome b-245 α (*CYBA*) for NADPH oxidase activity, high mobility group proteins (*HMGB2, HMGN2*) for chromatin remodeling and immune signaling, and fibroblast growth factor binding protein 2 (*FGFBP2*) for inflammatory microenvironment signaling. The presence of granzyme B (*GZMB*) indicates maintained cytotoxic potential during proliferation. *FGFBP2* and *GZMB* are known marker genes of a previously described NK cell subtype: NK1/hNK_Bl1 cells.(*9, 50*) Sphingosine-1-phosphate receptor 5 (*S1PR5*) suggests lymphoid tissue egress capacity. We also note that in addition to being a known modulator of inflammatory signaling, HMGN2 has a reported role in antimicrobial activity(*51*). Furthermore, microtubule regulation is important not just in the context of the cell cycle, but also for granzyme export.

Therefore, this archetype represents expanding NK cell populations that retain effector functions while undergoing active cell division. These molecular signatures show that particular motifs of antimicrobial resistance and immune defense may be compatible with and efficiently consolidated under a mitotic phenotype.

These six NK cell populations reveal the sophisticated functional architecture underlying immune surveillance, from highly motile infiltrators to specialized killers to immune orchestrators. Recently, a comprehensive characterization of NK cells based on CITEseq and snRNAseq identified six subtypes of NK cells and aimed to standardize NK cell ontology.

However, these subtypes were also acknowledged to be a continuum.(*9*) The alignment of the archetypes with the subtypes identified by Rebuffet et al. supports the archetypes as biologically meaningful functional states rather than technical artifacts, and provides a natural resolution to the type versus state distinction.

### Continuous variation within T cell subtypes

We identified five distinct T cell archetypes based on gene expression signatures, each representing functionally specialized immune states with unique biological roles and tissue adaptations, **Figure 5C**. The first archetype represents metabolically active effector memory T cells with enhanced biosynthetic capacity. *S100A4* serves as a key marker of memory T cell status and is involved in cytoskeletal dynamics and migration.(*52*) The signature includes genes for protein synthesis (*TPT1, ACTB*), cytoskeletal regulation (*TMSB4X*), metal ion homeostasis (*MT2A*), and antigen presentation (*HLA-A, B2M*). *IL32* indicates pro-inflammatory cytokine production capability. This population likely represents tissue-resident effector memory T cells that maintain rapid response capabilities through enhanced metabolic activity and protein synthesis machinery.(*53*)

The second archetype reflects T cells with balanced cytotoxic and regulatory functions. *SRGN* is associated with cytotoxic granule organization, while *CST7* provides protease inhibition to control cytotoxic responses. *CREM* acts as a transcriptional regulator of effector cytokines(*54*), and *PDCD4* functions as a translational repressor that can be modulated during activation. *HLA-C* contributes to antigen presentation capabilities and may influence interactions with NK cells or T cells through its unique binding properties compared to other MHC class I molecules. *RGCC* contributes to cell cycle control during proliferative responses. This archetype appears to represent professional cytotoxic T cells with built-in regulatory mechanisms to prevent excessive tissue damage while maintaining effector capabilities.

The third archetype defines T cells specialized for hostile microenvironments with enhanced stress tolerance. *TSC22D3* (*GILZ*) mediates glucocorticoid-induced anti-inflammatory responses, while *ANXA1* provides phospholipase A2 inhibition for anti-inflammatory activity. *JUN* contributes to AP-1 transcriptional responses, and the heat shock proteins (*DNAJA1, HSPA6*) provide protein quality control under stress conditions. *BRD3* broadly regulates inflammatory gene expression through histone binding and recruitment of transcriptional machinery, although its specific role in T cells remains unclear.(*55*) This population likely represents tissue-resident T cells adapted to maintain function in challenging microenvironments while preventing excessive inflammatory responses.(*56*)

The development and activation signature present in the fourth archetype characterizes transitional T cell populations maintaining developmental plasticity. *IL7R* expression indicates dependence on IL-7 for survival and homeostatic maintenance, typical of naive and memory precursor cells. *PDE4D* provides cAMP regulation during T cell activation and long-term responses.(*57*) *ZFP36L2* offers post-transcriptional control by destabilizing cytokine transcripts.(*58*) This archetype likely includes expression gradients in naive T cells transitioning to memory states, central memory T cell precursors, and potentially precursor exhausted T cells that retain proliferative and differentiation potential.

The fifth archetype represents T cell intratissue organization and management of oxidative stress during activation. *CD69* serves as a tissue residency marker that prevents lymph node homing, while *CXCR4* is broadly expressed across T cells(*59*) and provides responsiveness to CXCL12 spatial gradients in tissues. The antioxidant machinery includes *FTL* for iron sequestration, *ACAT1* for metabolic regulation, and *GCLM* for glutathione synthesis, which buffers ROS produced in activated T cells.(*60*) *RMND5A* supports mitochondrial respiratory function. This archetype captures continuous variation in *CXCR4* expression across the T cell population, and shows the metabolic machinery that enables T cell response.

Typically, T cells are divided in subtypes based on their place in developmental, activity, and functional hierarchies. The T cell archetypes capture core functions of T cells that vary within T cell subtypes, and are shared across them. We would thus interpret these functions as core to the T cell identity, and providing complementary information to the reported T cell subtypes.(*7*) Archetype one provides rapid effector responses, archetype two balances cytotoxicity with regulation, archetype three maintains tissue protection under stress, archetype four preserves developmental plasticity, and archetype five manages oxidative stress during tissue-resident activation.

### Fibroblast archetypes highlight distinct roles in ECM maintenance

Fibroblasts are found in all tissues and have increasingly recognized regulatory roles beyond ECM building. We find three pan-tissue fibroblast archetypes that reflect roles in ECM maintenance, immune modulation, and environmental responsiveness, **Figure 5D**. The first fibroblast archetype represents a specialized collagen production and matrix remodeling phenotype. This population is characterized by high expression of multiple collagen types (*COL1A1, COL1A2, COL3A1, COL4A2, COL6A2, COL6A3*), suggesting it functions as a collagen synthesis specialist. Beyond collagen production, this archetype shows matrix remodeling capabilities through matrix metalloproteinase expression (*MMP2*) and additional ECM components including laminin (*LAMB1*), biglycan (*BGN*), and vimentin (*VIM*). The presence of growth factor binding proteins (*IGFBP7*) and matricellular proteins (*SPARCL1, FSTL1*) indicates this population not only produces structural ECM components but also creates signaling niches that modulate growth factor availability and cell-matrix interactions. Finally, *APOD* expression has been recently implicated in fibroblast dysfunction, although its specific role remains unclear.(*61, 62*) This work implicates APOD in physiological fibroblast function, potentially related to supporting collagen synthesis or matrix remodeling through antioxidant properties.

The second archetype displays a robust cellular defense and metabolic adaptation phenotype. This population is distinguished by high expression of metallothioneins (*MT2A, MT1M, MT1X*), which protect against oxidative damage, heavy metal toxicity, and inflammatory insults. The co-expression of ribosomal proteins (*RPL13, EEF1A1*) and stress-responsive transcription factors (*JUNB*) suggests elevated protein synthesis capacity alongside stress adaptation. This archetype likely represents fibroblasts primed for rapid response to environmental challenges while maintaining active protein production machinery.

The third archetype combines ECM structural functions with innate immune regulation. This archetype highly expresses complement factor D (*CFD*), which plays a dual role as both a complement system activator and an adipokine involved in metabolism and tissue repair. The expression of additional complement components (*C3, C1R*) positions these fibroblasts as regulators of local immune responses. In addition, the expression of ECM organizing proteins including fibulin (*FBLN1*), gelsolin (*GSN*), decorin (*DCN*), and cystatin C (*CST3*) suggests these cells orchestrate ECM assembly while simultaneously modulating complement cascade activation, potentially serving as sentinel cells that integrate tissue homeostasis with immune surveillance. This archetype also expresses insulin-like growth factor binding protein 6 (*IGFBP6*), which is thought to play an important role in modulating matrix deposition following immune infiltration. However, the specific mechanisms remain to be fully elucidated.(*63, 64*) The separation of collagen production from broader ECM organization and immune functions reflects that different aspects of connective tissue biology require distinct cellular specializations and biochemical environments.

### Endothelial cell archetypes capture their roles in tissue homeostasis

In our analysis of endothelial cell function, we identified three archetypes that reflect their roles in tissue homeostasis, repair, and protection, **Figure 5E**. While labeled “angiogenesis” by an LLM, the first archetype represents a more complex collection of functions characteristic of capillary endothelial cells. This archetype is defined by *SPARCL1*, a matricellular protein originally discovered in and historically associated with HEVs that regulates lymphocyte trafficking and vessel homeostasis.(*65*) The archetype also expresses *RGCC*, a capillary endothelial cell marker(*66*) and a regulator of cell cycle that enables the rapid remodeling capacity, and *KDR* (VEGFR2), indicating responsiveness to angiogenic signals. Additional components include *OSBPLA1*, a regulator of lipid metabolism and membrane organization, *NCOA7*, a regulator of metabolism, *S100A4*, a regulator of cytoskeletal dynamics and cell motility, and *HLA-C* for antigen presentation at the blood-tissue interface. This gene signature recapitulates the standard collection of functions performed by capillary endothelial cells(*67, 68*), adapted for monitoring local tissue conditions, facilitating selective immune cell trafficking, and maintaining vessel stability while retaining the capacity for rapid functional transitions. We also note that, because the arteriovenous transition represents a trajectory over space, this archetype captures a spatial coordinate of cell type organization within tissue.

The second endothelial cell archetype is defined by an enriched program focused on cytoskeletal regulation and cellular motility. The key markers of this archetype are β-thymosin family actin-sequestering proteins (*TMSB4X, TMSB10*) that play an important role in cytoskeletal organization by binding to and sequestering actin monomers, thereby inhibiting actin polymerization.(*69*) Both thymosin-β4 and -10 have been implicated in the positive or negative regulation of VEGF expression, angiogenesis, and endothelial cell migration.(*70, 71*) The expression of the cytoskeletal regulator and cell motility promoter *IGFBP3* reinforces this archetype’s specialization in dynamic cytoskeletal remodeling.(*72, 73*) This archetype also expresses *KCTD12*, which encodes an auxiliary subunit of the GABA^B^ receptor.(*74*) Despite its implication in cancer, little is known about the physiological function of KCTD12.(*75, 76*) Its presence in this archetype may support a cytoskeletal organizational role. Overall, this molecular signature indicates endothelial cells poised for rapid morphological changes, migration, and potentially involved in processes requiring extensive cellular reorganization such as sprouting angiogenesis, wound healing, or vascular remodeling.

The third archetype exhibits a comprehensive inflammatory response program centered around neutrophil activation and oxidative stress management. This archetype strongly expresses *CSF3* (*GCSF*), *CXCL8* (IL-8), and *SELE* (E-selectin), which coordinate neutrophil production, recruitment, and recognition of cytokine-activated endothelial cells, respectively. CXCL8 is one of the most potent neutrophilic chemokines and additionally promotes endothelial cell proliferation and migration during tissue repair through CXCR2 binding on endothelial cells.(*77*) This archetype expresses a robust protective program against oxidative and inflammatory stress, including *SOD2* and metallothioneins (*MT1X, MT1E, MT2A*) for antioxidant and metal protection, *FTL* for iron sequestration, and *CD59*, a major inhibitor of complement-mediated cell lysis. *SOD2* encodes a superoxide dismutase that is essential for maintaining vascular function.(*78*) This archetype represents endothelial cells activated for immune surveillance and inflammatory responses, capable of orchestrating neutrophil recruitment, managing oxidative stress, and coordinating tissue defense mechanisms.

These three endothelial archetypes represent distinct functional states that likely correspond to different vascular microenvironments and physiological demands. The angiogenesis-regulatory archetype maintains vessel homeostasis while retaining responsiveness to growth signals, the motility-specialized archetype facilitates dynamic vascular remodeling and repair, and the inflammatory-activated archetype manages immune responses and oxidative stress. This functional specialization allows the endothelium to adapt to diverse, tissue-specific requirements while maintaining its essential barrier and transport functions.

## Discussion

There are numerous methods to draw insight from continuous expression data, and many have been applied to single cell transcriptomic data. PCA, ICA, NMF, WGCNA, and topic modeling reveal co-expression modules (*79*), and trajectory inference methods, including Diffusion Pseudotime, reconstruct lineage trajectories, thus inferring dynamics.(*80, 81*) However, *none* inherently ties their outputs to specific functional tasks. In contrast, Pareto optimality predicts that vertices of the data’s convex hull reflect distinct functions.(*12, 13, 15, 17*) Archetypal analysis is then the natural choice to identify vertices on the hull.(*22*) The advantages of archetypal analysis over SVD, PCA, ICA, NMF, and clustering have been clearly articulated in the literature, and we refer the reader to Mørup and Hanson 2012 for a succinct discussion on the matter.(*82*) Note that archetypal analysis describes the data using a basis consisting of data points—in this case cells, not gene expression programs.

Our results indicate that we can answer the fundamental question posed at the beginning of this paper in the affirmative: The gene expression of most cell types is constrained to low-dimensional polytopes, the expression of key genes is significantly enriched at the vertices of those polytopes, and those genes correspond to functions performed by those cell types. Thus, we conclude that phenotypic variation within most cell types is dominated by Pareto optimality. It directly follows that compositions of optimal cell states account for much of the observed intra-cell type variation in the human body. Finally, we inferred which tasks of each cell type are efficiently performed in tandem, and which tasks must be divided between different states of that cell type. These results are robust as they are reproducible across donors and across different tissues with shared cell types. However, it is important to note that there are some limitations to the analysis and that not every cell type falls neatly into this paradigm.

First, there is clearly some experimental variation as for a given cell type, while a majority of donors may fit a polytope, some donors do not, and the source of this variation is unclear at this time. Also, a few cell types are not well fit by polytopes and it is unclear if this reflects their true biology or whether it is related to experimental artifacts. Nonetheless, it is notable that some of the most widely shared cell types across tissue are well described by polytopes and that the vertices can often be interpreted in terms of their biological function.

One possible source of artefacts in gene expression is tissue processing.(*16, 23*) After attempting to control for this with several conservative filters, and ensuring reproducibility of our findings across donors and tissues, we did not, in general, observe artefactual genes appearing in our gene lists of interest. An exception might be the stress response archetype in fibroblasts, but we cannot say for certain that this phenotype is *only* present due to tissue processing artefacts; fibroblasts are known to be early responders to stress in healthy physiology.

While we do not aim to elucidate the full expressional repertoires of each cell type, it may also be possible to increase the dimension of the polytope fits to uncover richer descriptions of cell repertoires. In the present analysis, we can draw conclusions about what might be the dominating tasks and how several tasks may be consolidated under a shared biochemical signature. Several transcripts discussed here have been previously implicated in niche-specific roles. Here, we implicate these genes as important mediators on a much broader scale. For example, SELENOP has previously been observed to be important to macrophages in muscle tissue, but this pan-tissue analysis supports a broader importance to tissue-resident macrophages. Many of the key macrophage genes that we have previously been implicated in various inflammatory pathologies, especially cancer. In our analysis of non-pathologic tissues, we support these gene’s roles as key to general physiology. Additionally, we found several genes whose mechanisms and functions have not been fully elucidated, including *SH3BGRL3, FAM177A1, BRD3, APOD*, and *KCTD12*. This analysis allows us to speculate about their functional roles.

Finally, Pareto optimality makes no statements about transitions or trajectories between states. In different contexts, optimal states can be terminal or not (see (*23*) for an example of terminal differentiation at an archetype); the extent to which the same actual cells move between archetypes is still an open question.(*20, 83*) In fact, a recent analysis modeled cells jointly by Pareto optimality and cell-circuit signaling loops to understand how regulatory dynamics impact differentiation.(*84*) Ultimately, we conclude that this principled method allows characterization of the collection of functions performed by a cell type.

## Conclusion

We used a principled method to identify archetypes that represent core functions performed by a cell type. For several ubiquitous cell types, we recapitulated known functions without incorporating prior biological knowledge. Placing these cellular tasks on a continuum, with cells performing some combination of specialized functions, advances our understanding of cell typology, beyond discretization of cell populations. Compared to canonical marker gene analysis, this continuous framing allows identification of functional marker genes that is unbiased, and captures continuous cellular heterogeneity. This approach provides a powerful way to combine two disjoint perspectives about cell type and cell state: whether they are discrete states or continuous states. The idea of having multiple discrete functions performed by a single cell type, and different cells within that type having a different balance of performing those functions that varies in a continuous fashion, provides a powerful paradigm to interpret and understand cell type and cell state distinctions.

## Materials and Methods

### Data sources

Single-cell RNA sequencing data was downloaded from public repositories. Sample preparation and processing has been previously described. Briefly, the sc-RNAseq dataset was previously processed including doublet removal, barcode-hopping correction, removal of cells expressing less than 200 genes, and removal of genes expressed in fewer than 3 cells. We also defined artefact genes as those potentially affected by tissue processing and dissociation.(*16*)

### Data Preprocessing

Additional preprocessing and filtering for the present study was performed in python 3.9 using Scanpy v1.10.3 (custom fork modified to add the intercept back to the results of scanpy.pp.regress_out(): https://github.com/ggit12/scanpy). We isolated 10x-processed single cells for analysis. We removed genes expressed in fewer than 5 cells. We then followed a conservative approach to outlier removal and removed cells that were in the upper decile of any of: percent mitochondrial counts, percent artefact counts, number of genes expressed, total counts, and number of UMI counts. At this point, no cells had 100 counts or fewer of protein-coding genes (excluding mitochondrial and artefact genes, as this set of protein coding genes was eventually used to define the expressional phenotype in PCA space), and we mention this a-priori defined cutoff for completeness, **Figure S1** and **Table S1**. This strategy allowed us to focus on major expressional trends and ignore edge cases. Mitochondrial genes and genes affected by tissue processing were then removed before normalizing the expression matrix to 10,000 counts/cell. On the filtered, normalized data, a cell cycling score was calculated as in (*16*). The data was then stratified by cell type, donor, and tissue and each stratum was analyzed separately, treating donors as biological replicates. Finally, as a conservative approach to outlier removal and to stabilize subsequent PCA transformations on linear-scale data, we removed cells in the bottom 10 percent of density in the first 3 components of PCA space for each donor-tissue-cell type.(*22*)

### Data Analysis

#### Pareto Task Inference Analysis

Gene expression data and cell metadata were then read into MATLAB R2022b for further processing using the Pareto Task Inference package (ParTI).

ParTI analysis was performed on all protein-coding genes as defined above, in all donor-tissue-cell types with at least 50 cells available for analysis after all filters in *data preprocessing*, which prior simulations have shown to be sufficient for polytope detection.(*23*) Donor-tissue-cell types with more than 1000 cells were downsampled to 1000 cells. The ParTI algorithm was set to initialize with 5 dimensions and calculate the optimal number of vertices in the polytope based on an elbow-finding algorithm of the variance explained by successive archetypes (the default behavior). The ParTI package calculated p-values as the ratio of times that a polytope fit the real PCA data better than to randomly shuffled PCA data over 1000 trials. When fitting polytopes, we swept a narrow range of dimensions (5 to 2 PCs), yielding between 6-to 2-vertex polytopes, stopping when significance was observed at the optimal number of vertices for the given dimension. Prior works swept similar ranges(*17, 23, 85*), motivated by the following theory: The naïve expectation is that all the tasks of a system vary by orders of magnitude in terms of their impact on overall fitness, and only a few of the highest-magnitude tasks to dominate phenotype space, with other tasks consolidated others under generalist or specialist phenotypes.(*14*) When visualizing fitted polytopes, vertices may appear nearby each other in low-dimensional projections. This can represent a simple projection artifact, or the splitting of a single specialist phenotype into biologically redundant ones.

The genes significantly enriched at each archetype (significant after the Benjamini-Hochberg multiple tests correction with a maximum false discovery rate of 0.10) were then saved for archetype alignment. All genes, including mitochondrial and artefactual genes, were included in the enrichment analysis to allow detection of artifactual archetypes.

#### Statistical Analysis

To define the significance of a cell type, we set α=0.05 and maximum false discovery rate (FDR) =0.10. We then defined a cell type aggregation process, where a cell type was considered significant if 50% or more of available donors had a tissue with p < 0.05. For these cell types that qualified, we directly calculated the probability that the cell type was a false positive based on the number of donors and respective number of p-values calculated for each tissue of that donor (including excess p-values from testing multiple dimensions for significance), by assuming the false positive rate α of each donor-tissue-cell type, and applying a Bayesian correction factor for the empirical distribution of p-values (qvalue v2.30.0). This false positive probability per cell-type can then be used to directly calculate the FDR, see **Supplemental Code**. Finally, all cell types up to a total FDR < 0.10 were considered well-fit by polytopes. This resulted in a total FDR of 0.0849 for the 82 significant cell types in this study (or about 7 false positives). Note that in **Figure 3A**, we plot a single p-value per donor-tissue-cell type (the significant p-value if available, otherwise the first-calculated) to visualize the proportion of significant donor-tissue-cell types, while for statistical purposes and calculation of false discovery statistics, we used the full distribution of all p-values calculated, which is a conservative approach that may artificially overinflate the estimated false discovery rate.

#### Quality Control

To ensure quality control of the fits, we represented quality control metrics in several ways. First, we plotted p-values of fits vs. each of the quality control metrics used in filter (except total counts protein-coding) and the total number of cells included in the fit, for all donor-tissue-cell types, and for donor-tissue-cell types from the five major cell types discussed in main, **Figure S2**. Then, to get a general assessment of the degree to which the quality control metrics may have affected the polytopal fits, we calculated the correlations of several metrics with the percent of cells that were within the bounds of the fitted polytope, **Table S2**. Note that the main fitting algorithm (SDVMM) initialized with data points as vertices, and finds maximal polytopes that lie *within* the data by expansion, so it is expected that some cells are outside the bounds of the fitted polytope. Only very weak (|r| < 0.2) correlations were observed between any of the metrics we considered and the percent of cells that ended up within the final fit. In Figure S2 and Table S2, we used a single p-value per donor-tissue-cell type to attempt to detect quality control factors that may have impacted significant results; the same quality control analysis run on the full distribution of p-values showed smaller effect sizes.

#### Archetype Alignment

Our goal was to identify reproducible archetypes (i.e. expressional programs) of each cell type. Therefore, we grouped all vertices by cell type, yielding several donor-tissues for each cell type, with each of these donor-tissues having multiple vertices. Each vertex represents a specialist phenotype that is defined by a ranked list of enriched genes. We used the number of shared genes between each specialist phenotype to align these phenotypes, **Figure 1B**. This task was accomplished with the following graph-based approach. We defined an adjacency matrix for all specialist phenotypes found in a cell type based on the number of genes shared between each pair of specialist phenotypes. We then clustered (igraph v1.5.1) the specialist phenotypes according to the graph defined by this adjacency matrix, thereby aligning specialist phenotypes based on their shared genes. When at least 50% of biological replicates (i.e. donors) were represented within a cluster of specialist phenotypes, this cluster of specialist phenotypes was considered an archetype that represented a reproducible biological function of that cell type.

This approach also elegantly handles the potential over-splitting of specialist phenotypes discussed above by aggregating biologically redundant specialist phenotypes from single donor-tissues. Then, redundant specialist phenotypes within a donor-tissue do not contribute to a cluster of specialist phenotypes meeting the donor-level reproducibility filter.

#### Archetype Annotation

To annotate each archetype based on the frequency-ranked list of genes shared across comprising specialist phenotypes, we first focused on the most frequent genes by dropping genes with frequency one in the archetype (i.e. present in only one specialist phenotype within that archetype). Then, we used a large language model (Claude 4 Sonnet, as discussed in main) to annotate all archetypes across all cell types by passing the gene list of each archetype to the large language model and asking what biological process the gene list most likely represented, **Supplemental Code**. Notably, the large language model was *not* informed of the associated cell type, but, in several cases, still mentioned the correct cell type in the annotation. We accessed Claude 4 Sonnet through the Anthropic API with the R package claudeR v0.0.0.9000. This automation was needed to handle the large scale and breadth of this interpretative task. We tested several LLMs, including GPT-4, GPT-4o, Claude 3 and 4.1 Opus, and Claude 3.5 and 4 Sonnet. Results were generally similar across all the LLMs; we used Claude 4 Sonnet for final annotations as the Claude Sonnet series has been shown to be the most performant at gene set annotation.(*25*)

#### Computational Resources

Computation for ParTI analysis was done on a high-performance computing cluster with a Slurm scheduler in Matlab R2022b. Downstream analysis was performed in R 4.2.1.

#### Code Availability

All source code required to reproduce this analysis will be posted publicly upon publication.

## Acknowledgements

We thank G. Ed Marti for his helpful discussion of statistical methods. We thank Madhav Mantri and Jaeyoon Lee for their discussion of the investigation. We thank Avi Mayo and Miri Adler for their discussion of the ParTI analysis package.

Some of the computing for this project was performed on the Sherlock and Bruno clusters. We would like to thank Stanford University, the Stanford Research Computing Center, and the Chan-Zuckerberg Biohub for providing computational resources and support that contributed to these research results.

## Author contributions

Conceptualization: GC, UW, SRQ Data curation: GC, UW, SRQ Formal analysis: GC, UW, SRQ Funding acquisition: SRQ Investigation: GC, UW, SRQ Methodology: GC, UW, SRQ

Project administration: GC, UW, SRQ Resources: GC, UW, SRQ

Software: GC, UW, SRQ Supervision: UW, SRQ Validation: GC, UW, SRQ Visualization: GC, UW, SRQ

Writing – original draft: GC, UW, SRQ Writing – review & editing: GC, UW, SRQ

## Data and materials availability

All data used in this study is publicly available from (*16*). All source code used in this study will be posted publicly upon publication.

