## Supplemental Figures and Tables for "Pareto Optimality Reveals an Atlas of Cellular Archetypes"

### Supplementary Materials for

### An Atlas of Cellular Archetypes

George Crowley[1], Uri Alon[2], Stephen R. Quake[1,3 *]

**The PDF file includes:**

Figs. S1 to S5
Tables S1 to S2

**Other Supplementary Materials for this manuscript include the following:**

Supplemental File S1
Table S3
Supplemental Code

### Supplemental Figures

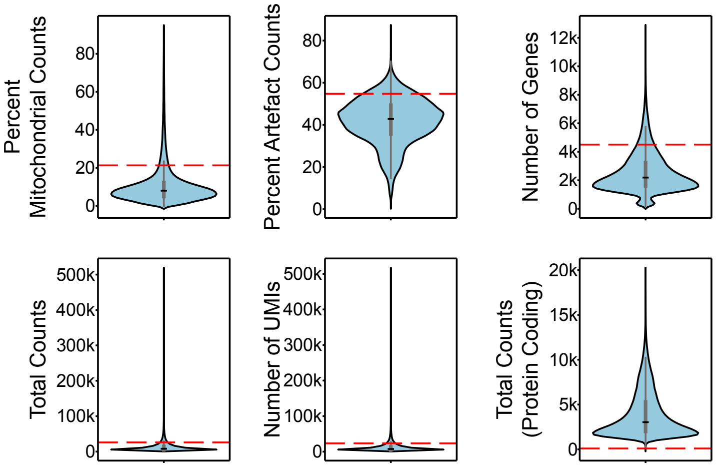


**Figure S1**. Distribution of percent mitochondrial genes, number of genes expressed, total counts, and number of UMI counts across cells. Vertical line indicates 90$^{th}$ percentile.

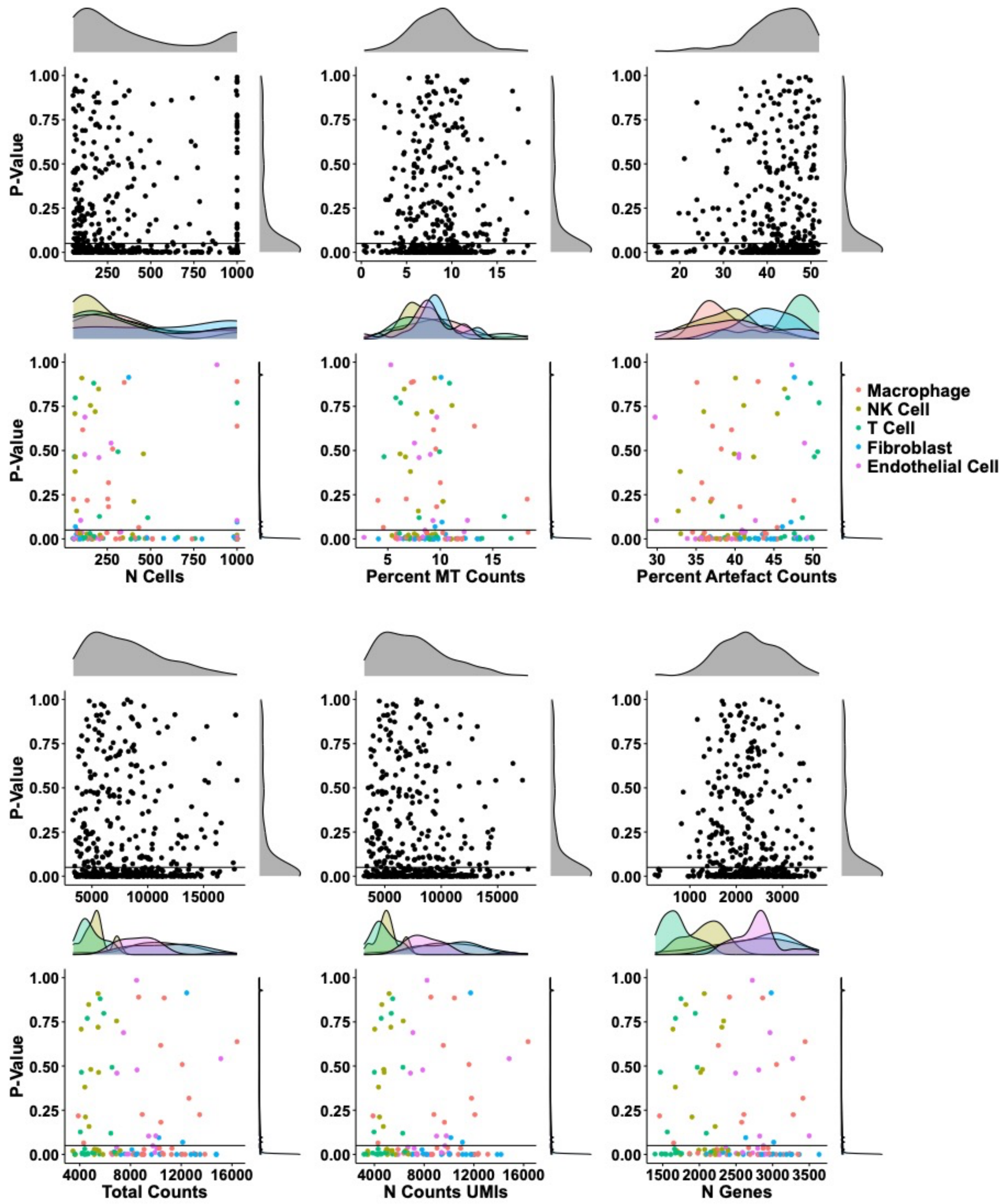


**Figure S2.** Scatterplots of p-value of polytopal fit vs. quality control metrics for all donor-tissue-cell types (greyscale) and for the five major cell types discussed in main (colored by cell type).

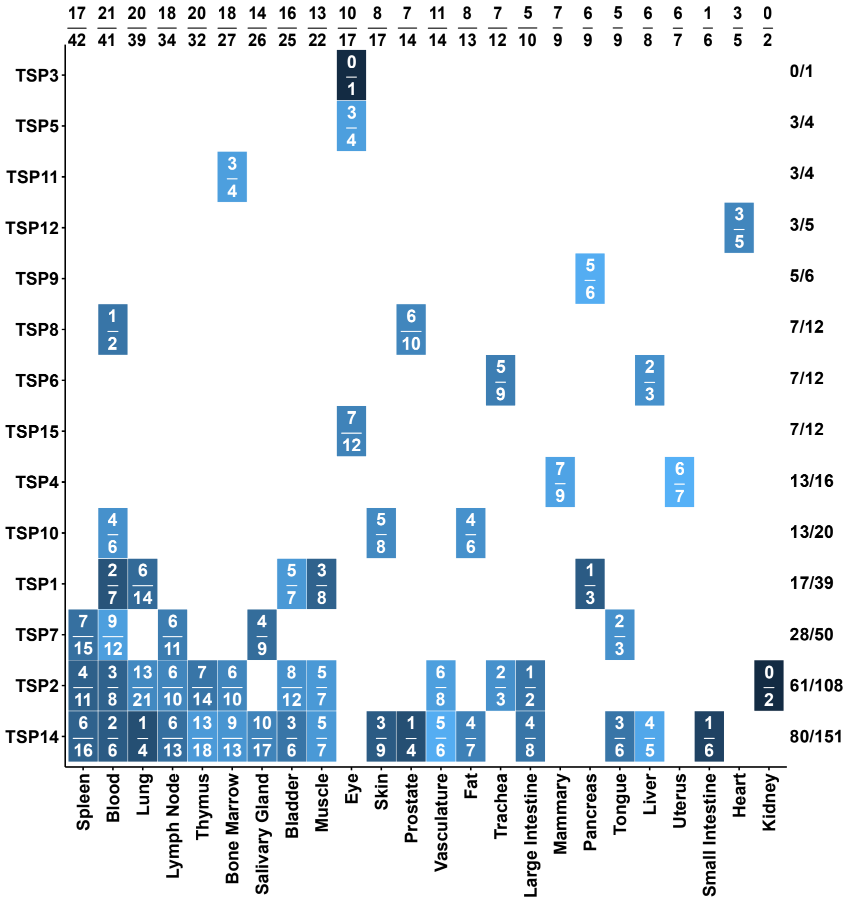


**Figure S3.** Donor-by-tissue heatmap of significant p-values, labeled with number of significant cell types over number of available cell types for each tile in the heatmap. The heatmap has marginal row and column sums of number significant over number available.

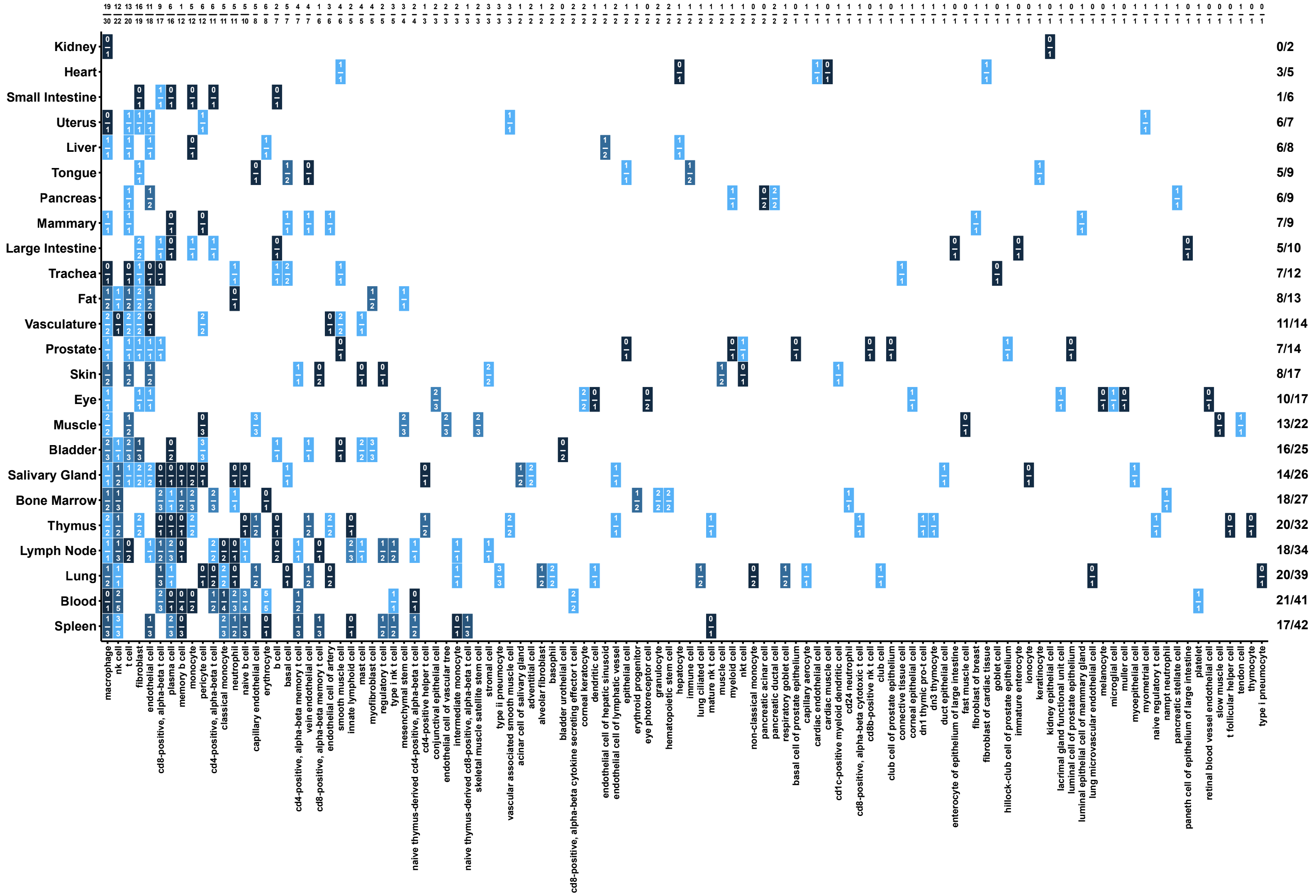


**Figure S4.** Tissue-by-cell type heatmap of significant p-values, labeled with number of significant donors over number of available donors for each tile in the heatmap. The heatmap has marginal row and column sums of number significant over number available.

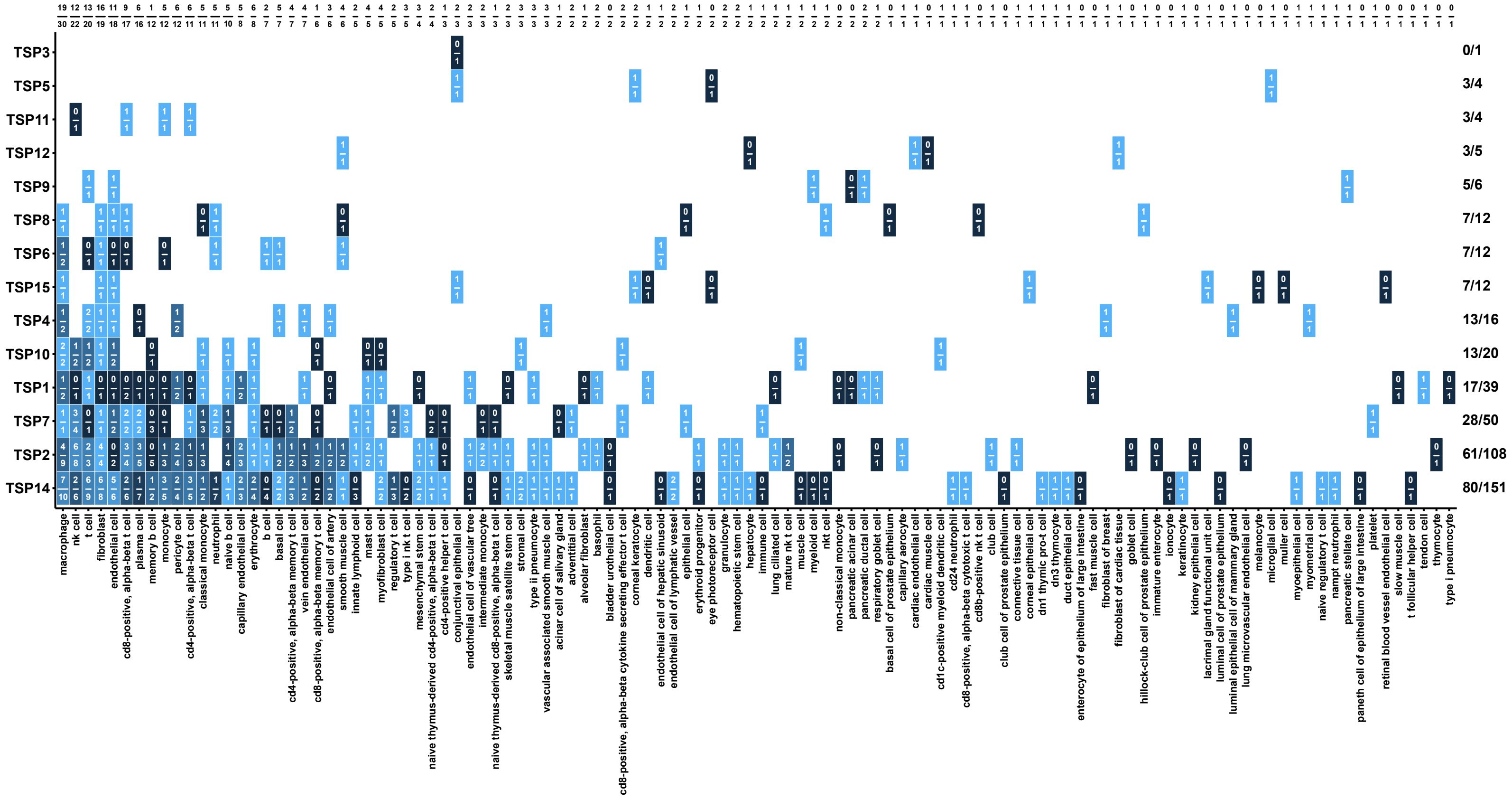

1090

1091 **Figure S5.** Donor-by-cell type heatmap of significant p-values, labeled with number of
1092 significant tissues over number of available tissues for each tile in the heatmap. The heatmap
1093 has marginal row and column sums of number significant over number available.

1094

1095

### Supplemental Tables

| Quality Control Metric | Threshold Value | N Cells Excluded | Bound |
|---|---|---|---|
| Percent Mitochondrial Counts | 21 | 45611 | Upper |
| Percent Artefact Counts | 55 | 45611 | Upper |
| Number of Genes | 4503 | 45636 | Upper |
| Total Counts | 25921 | 45612 | Upper |
| Number of UMIs | 23424 | 45617 | Upper |
| Total Protein-coding Counts | 100 | 0 | Lower |

Threshold values rounded to nearest integer.

**Table S1.** Quality control thresholds used in initial filtering of Tabula Sapiens and number of cells/genes beyond the bound of each threshold. All thresholds except “Total Protein-coding Counts” were based on the 90$^{th}$ percentile of the data.

| Percent of Cells in Polytope vs. | R | P-Value |
|---|---|---|
| P-value < 0.05 | 0.19 | 0 |
| Mean(Percent Counts Artefact) | -0.12 | 0.01 |
| Mean(Total Counts) | 0.1 | 0.04 |
| Mean(N Counts UMIs) | 0.09 | 0.07 |
| Mean FC(N Genes) | -0.06 | 0.21 |
| Mean(Percent Counts MT) | -0.05 | 0.29 |
| Mean FC(N Genes by Counts) | -0.03 | 0.59 |
| Mean FC(Percent Counts MT) | -0.02 | 0.62 |
| Mean FC(N Counts UMIs) | -0.02 | 0.74 |
| Mean(N Genes by Counts) | 0.02 | 0.66 |
| N Cells | -0.02 | 0.63 |
| Mean FC(Percent Counts Artefact) | 0.01 | 0.88 |
| Mean FC(Total Counts) | 0 | 0.93 |
| Mean(N Genes) | 0 | 0.92 |
| Mean Fold Change (Mean FC) | | |

**Table S2.** Correlations of quality control metrics with the percent of cells inside the fitted polytope.

**Table S3. (separate file)** Significantly enriched genes for all cell types and archetypes that passed filters described in main. Two sheets included: 1) Top_5_Cell_Types: information for the 5 cell types discussed in main based on having the most donor-tissues; and 2) All_Cell_Types: information for all cell types.

### Supplemental Files

**Supplemental File 1. An Atlas of Cellular Archetypes.** Archetype figures (same form as **Figure 3**) for all 35 cell types, including cell types not discussed in main manuscript.

**Supplemental Code.** Source code used for the analysis presented in this manuscript will be publicly released upon publication.
