## Supplemental File 1 for "Pareto Optimality Reveals an Atlas of Cellular Archetypes"

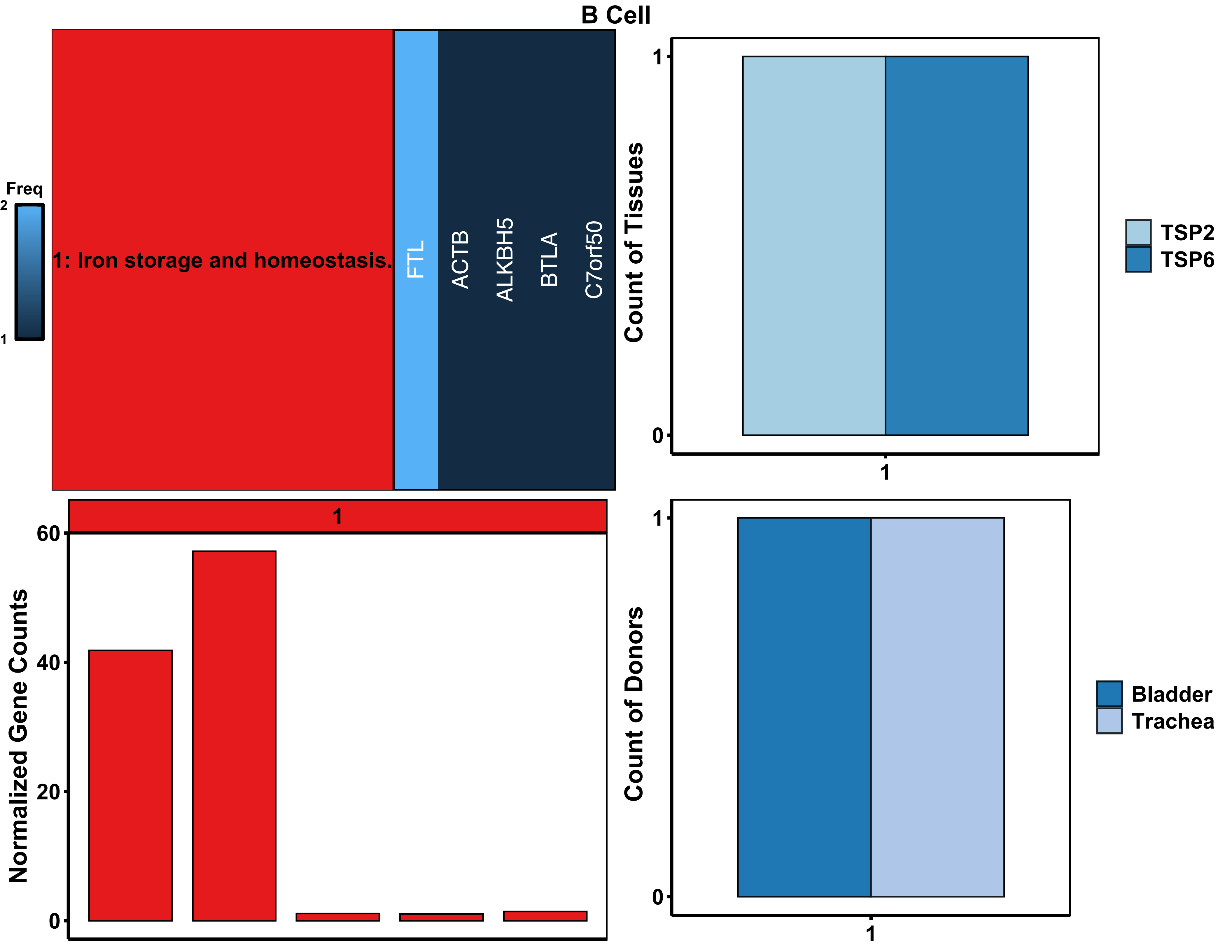

B Cell
Freq
2
1
1: Iron storage and homeostasis.
FTL
ACTB
ALKBH5
BTLA
C7orf50
Count of Tissues
1
0
1
TSP2
TSP6
1
Normalized Gene Counts
60
40
20
0
Count of Donors
1
0
1
Bladder
Trachea

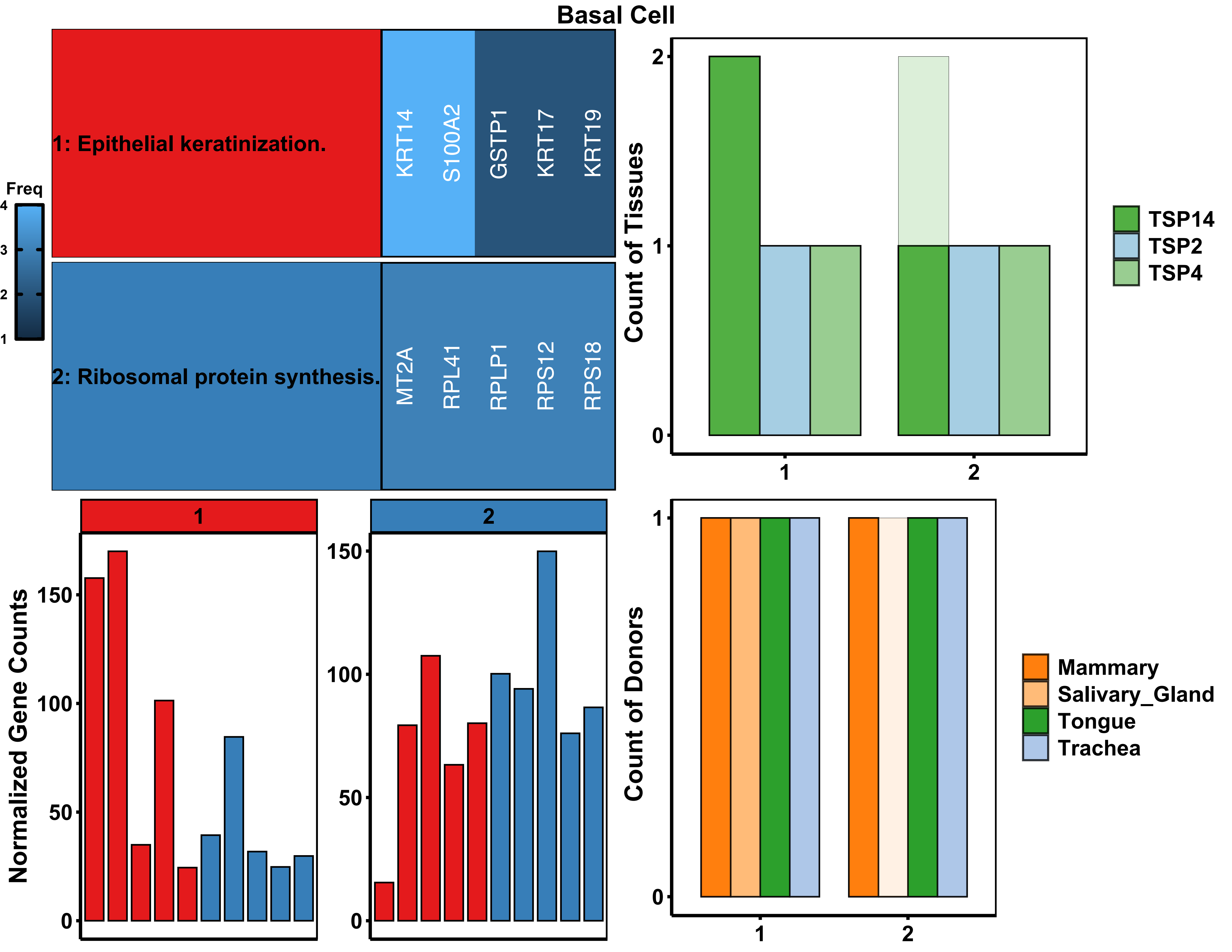

Basal Cell
1: Epithelial keratinization.
2: Ribosomal protein synthesis.
KRT14
S100A2
GSTP1
KRT17
KRT19
MT2A
RPL41
RPLP1
RPS12
RPS18
Freq
4
3
2
1
Count of Tissues
2
1
0
1
2
TSP14
TSP2
TSP4
1
2
Normalized Gene Counts
150
100
50
0
Count of Donors
Mammary
Salivary_Gland
Tongue
Trachea

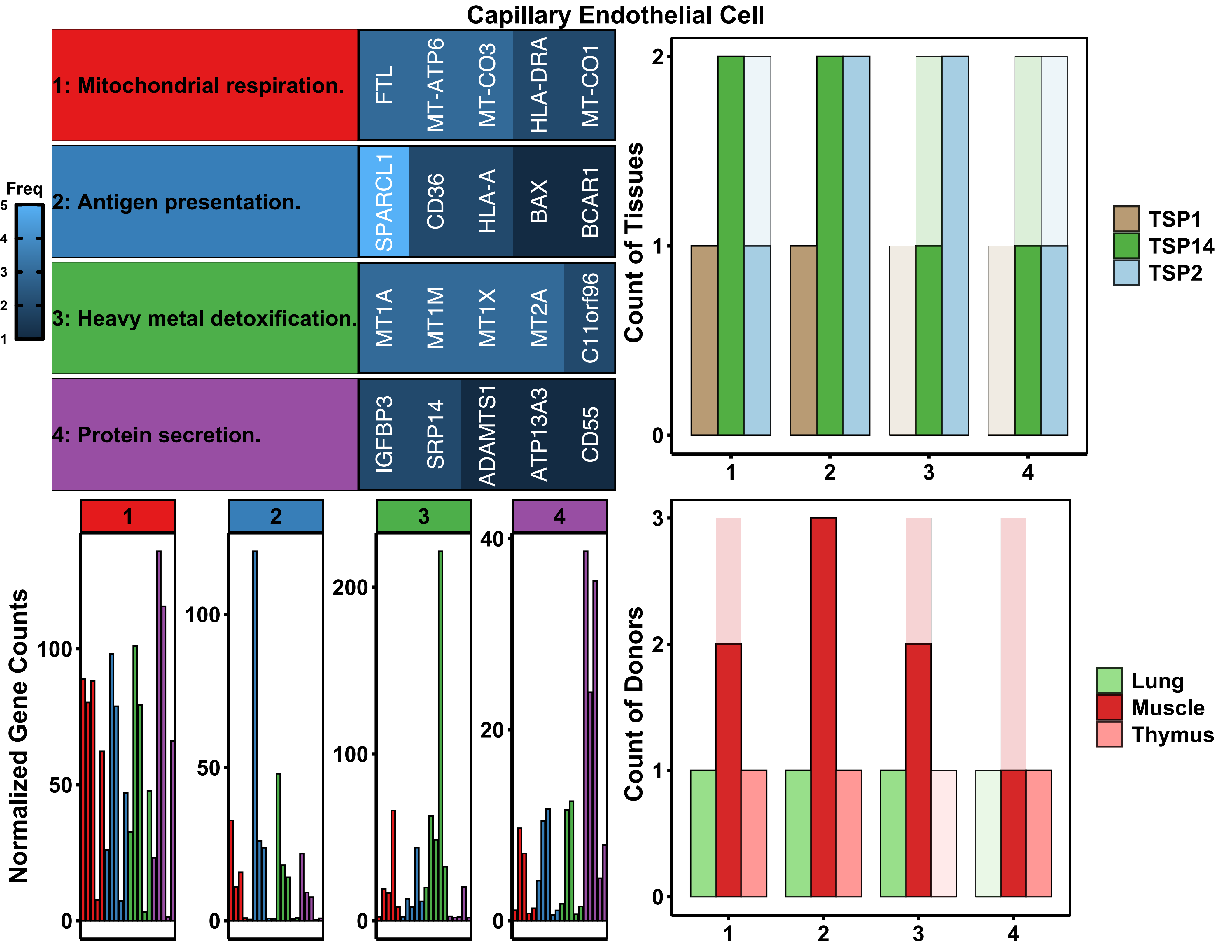

Capillary Endothelial Cell
1: Mitochondrial respiration.
FTL
MT-ATP6
MT-CO3
HLA-DRA
MT-CO1
2: Antigen presentation.
SPARCL1
CD36
HLA-A
BAX
BCAR1
3: Heavy metal detoxification.
MT1A
MT1M
MT1X
MT2A
C11orf96
4: Protein secretion.
IGFBP3
SRP14
ADAMTS1
ATP13A3
CD55
Freq
Count of Tissues
TSP1
TSP14
TSP2
Normalized Gene Counts
Count of Donors
Lung
Muscle
Thymus

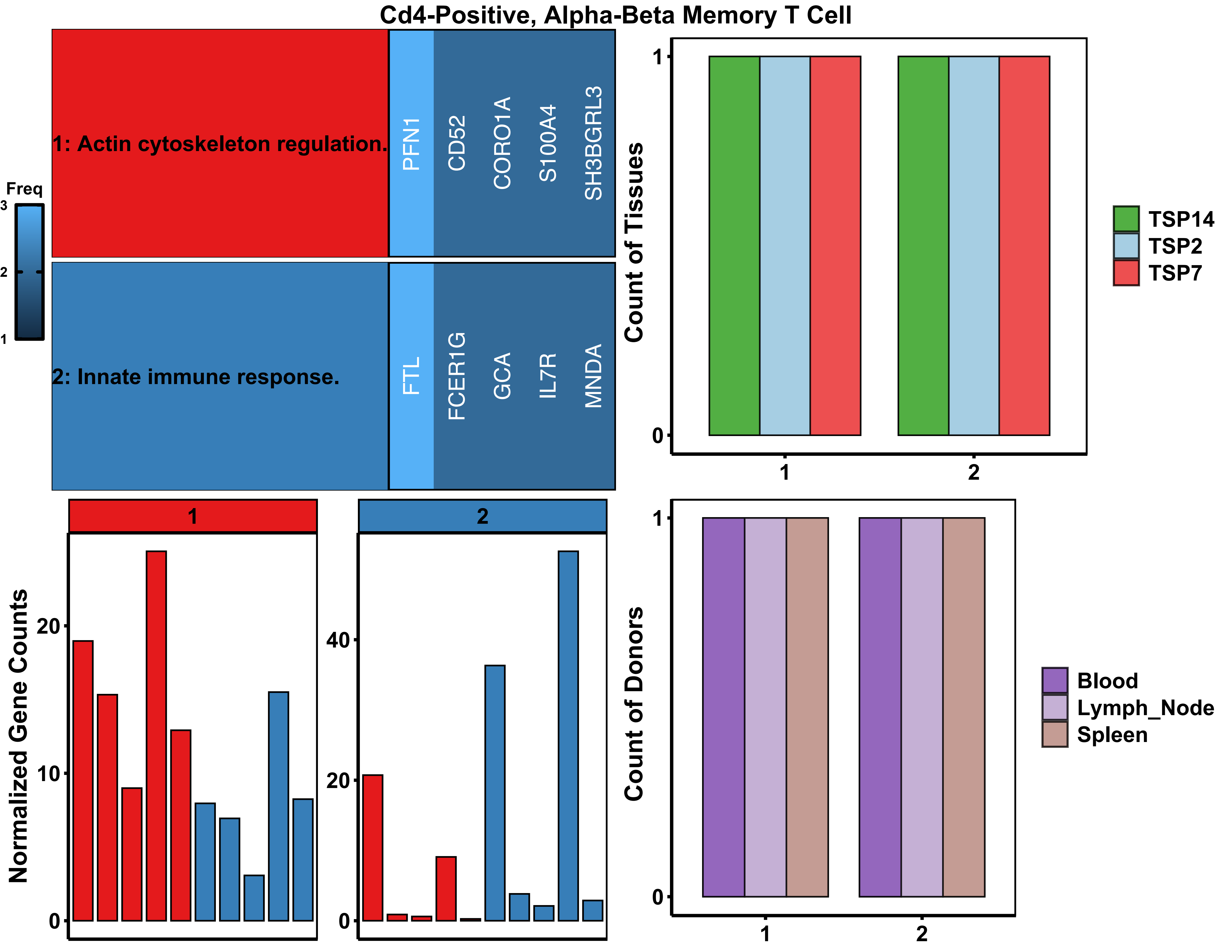

Cd4-Positive, Alpha-Beta Memory T Cell
1: Actin cytoskeleton regulation.
PFN1
CD52
CORO1A
S100A4
SH3BGRL3
2: Innate immune response.
FTL
FCER1G
GCA
IL7R
MNDA
Freq
3
2
1
Count of Tissues
1
0
1
2
TSP14
TSP2
TSP7
1
2
Normalized Gene Counts
20
10
0
40
20
0
Count of Donors
1
0
1
2
Blood
Lymph_Node
Spleen

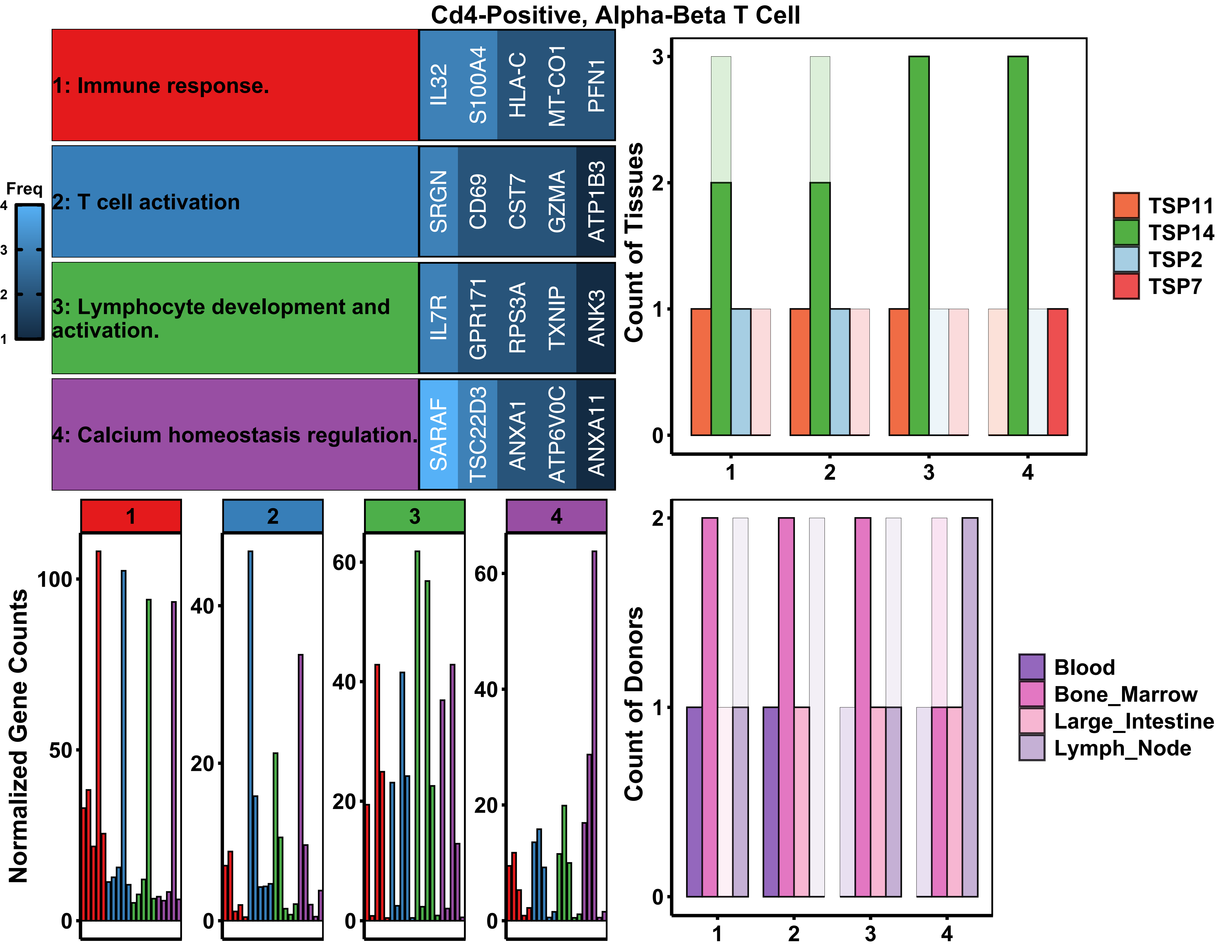

Cd4-Positive, Alpha-Beta T Cell
1: Immune response.
IL32
S100A4
HLA-C
MT-CO1
PFN1
2: T cell activation
SRGN
CD69
CST7
GZMA
ATP1B3
3: Lymphocyte development and activation.
IL7R
GPR171
RPS3A
TXNIP
ANK3
4: Calcium homeostasis regulation.
SARAF
TSC22D3
ANXA1
ATP6V0C
ANXA11
Freq
Count of Tissues
TSP11
TSP14
TSP2
TSP7
Normalized Gene Counts
Count of Donors
Blood
Bone_Marrow
Large_Intestine
Lymph_Node

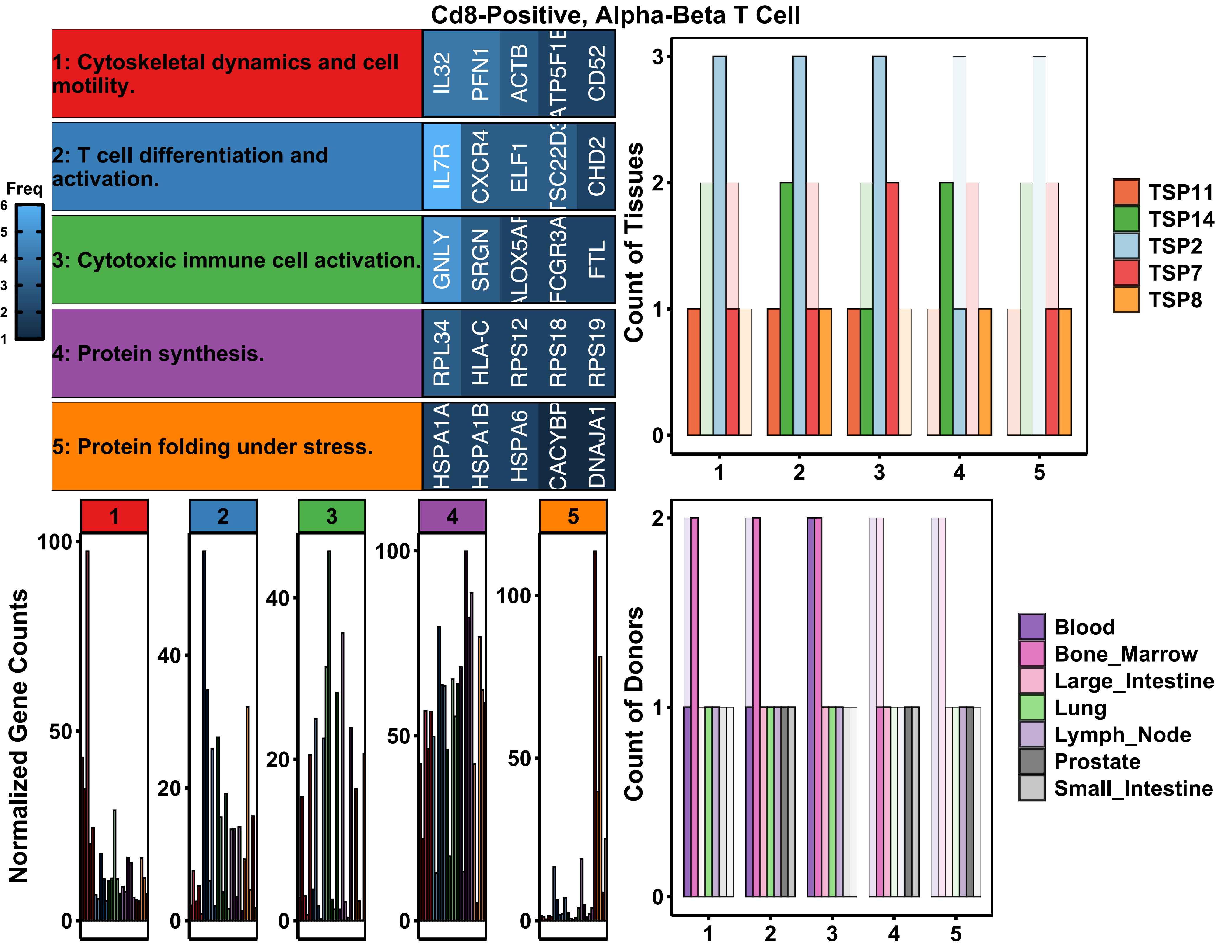
Cd8-Positive, Alpha-Beta T Cell
1: Cytoskeletal dynamics and cell motility.
IL32
PFN1
ACTB
ATP5F1E
CD52
2: T cell differentiation and activation.
IL7R
CXCR4
ELF1
CHD2
3: Cytotoxic immune cell activation.
GNLY
SRGN
FTL
4: Protein synthesis.
RPL34
HLA-C
RPS12
RPS18
RPS19
5: Protein folding under stress.
HSPA1A
HSPA1B
HSPA6
DNAJA1
Freq
Count of Tissues
TSP11
TSP14
TSP2
TSP7
TSP8
Normalized Gene Counts
Count of Donors
Blood
Bone_Marrow
Large_Intestine
Lung
Lymph_Node
Prostate
Small_Intestine

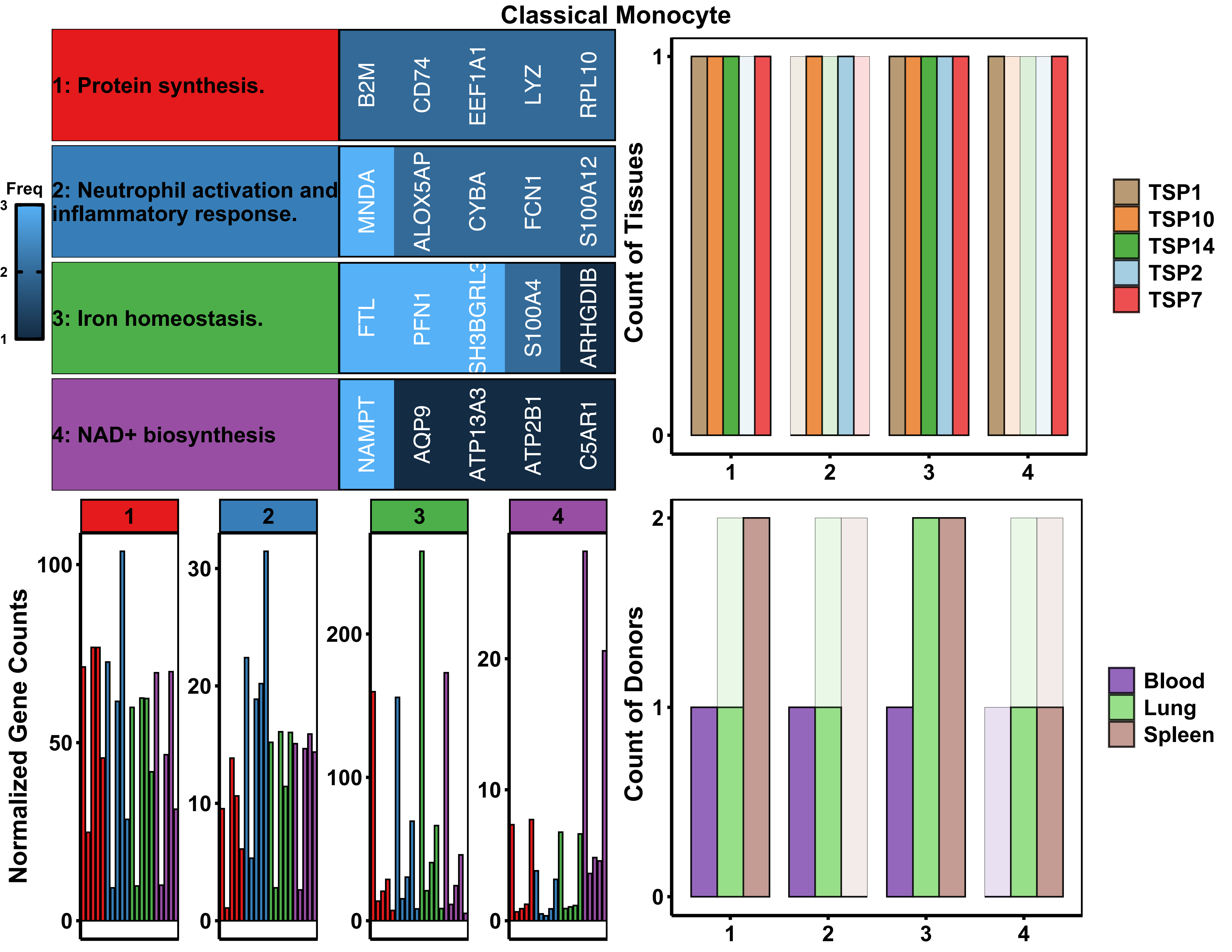
Classical Monocyte
1: Protein synthesis.
B2M
CD74
EEF1A1
LYZ
RPL10
2: Neutrophil activation and inflammatory response.
MNDA
ALOX5AP
CYBA
FCN1
S100A12
3: Iron homeostasis.
FTL
PFN1
SH3BGRL3
S100A4
ARHGDIB
4: NAD+ biosynthesis
NAMPT
AQP9
ATP13A3
ATP2B1
C5AR1
Freq
3
2
1
Count of Tissues
TSP1
TSP10
TSP14
TSP2
TSP7
Normalized Gene Counts
Count of Donors
Blood
Lung
Spleen

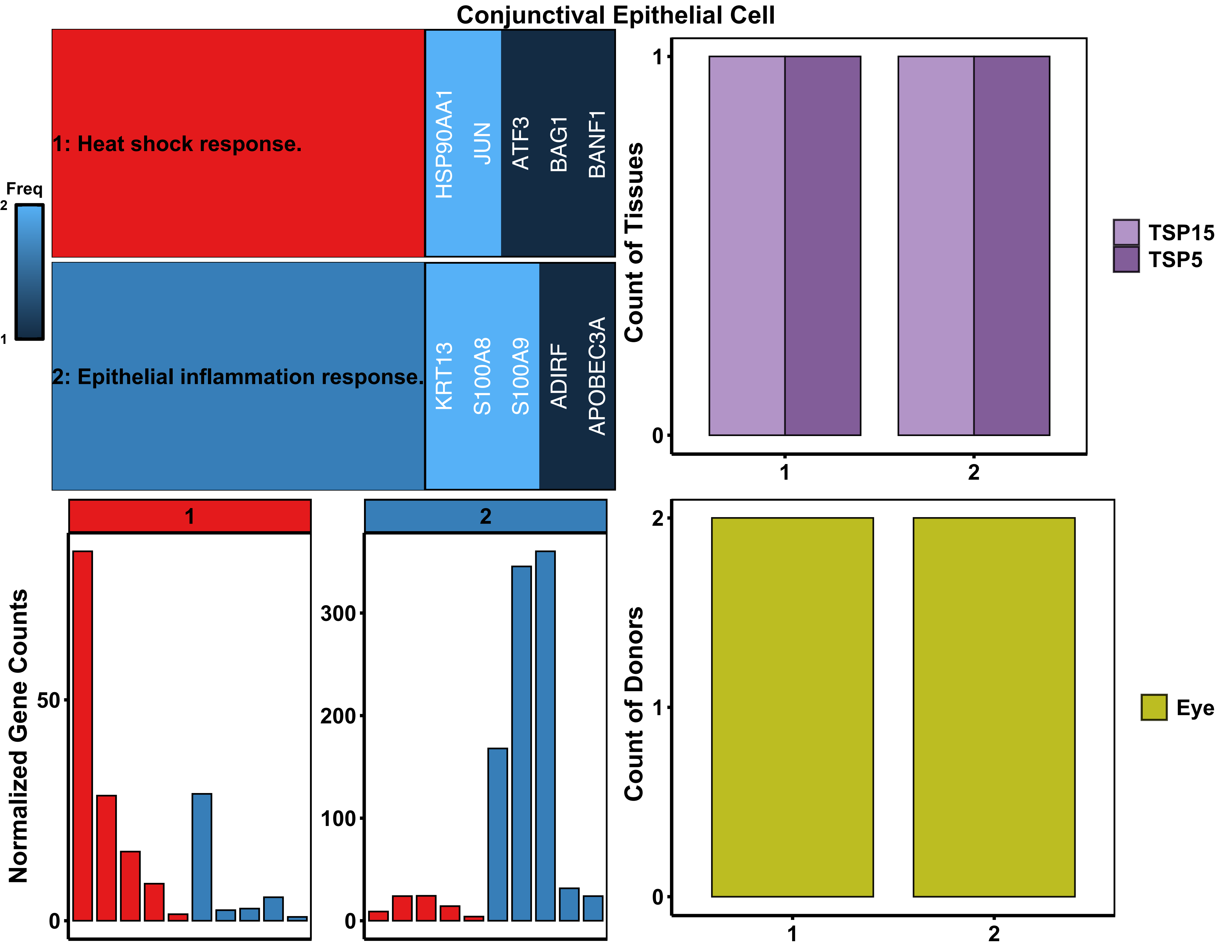

Conjunctival Epithelial Cell
Freq
2
1
1: Heat shock response.
HSP90AA1
JUN
ATF3
BAG1
BANF1
2: Epithelial inflammation response.
KRT13
S100A8
S100A9
ADIRF
APOBEC3A
Count of Tissues
1
0
1
2
TSP15
TSP5
1
2
Normalized Gene Counts
50
0
300
200
100
0
Count of Donors
2
1
0
1
2
Eye

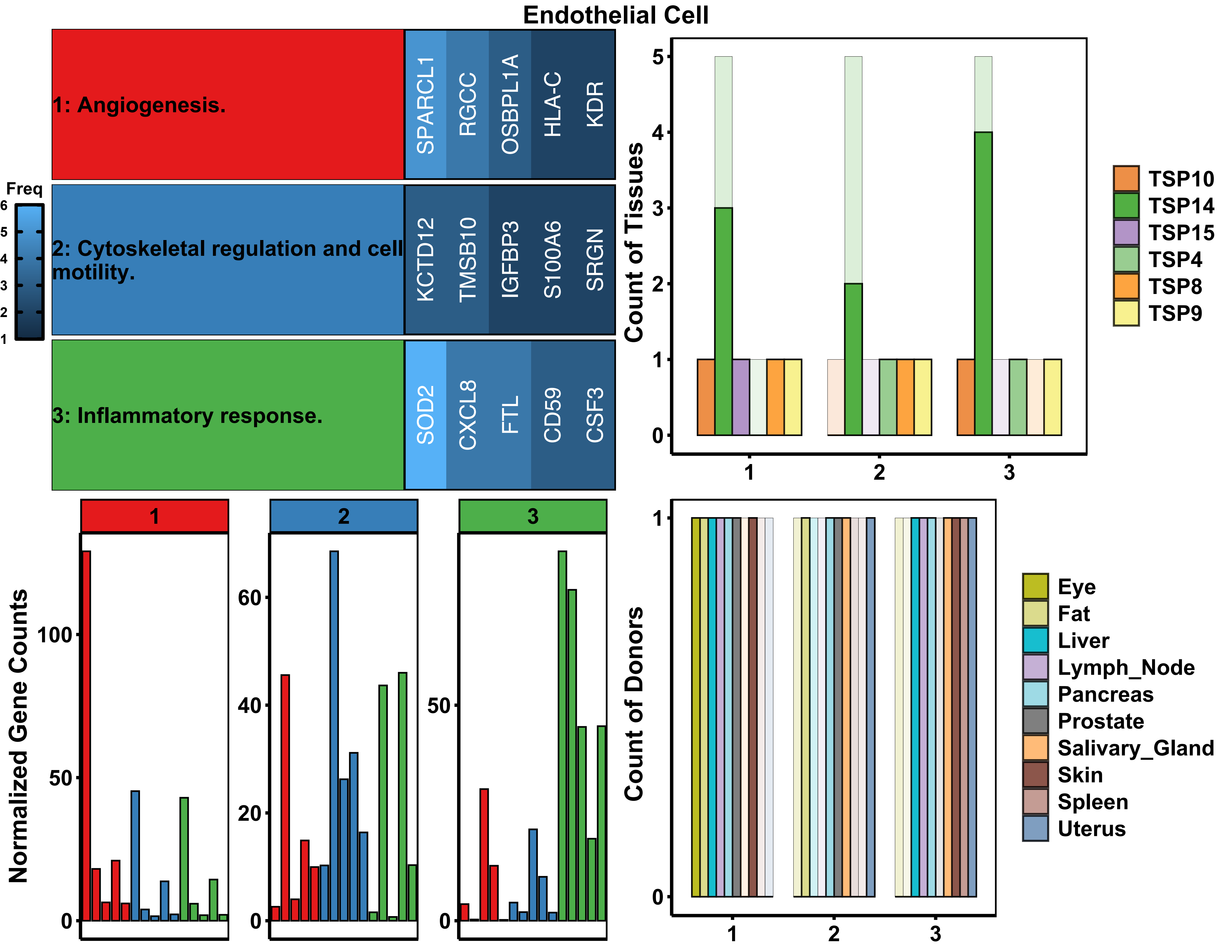
Endothelial Cell
1: Angiogenesis.
SPARCL1
RGCC
OSBPL1A
HLA-C
KDR
2: Cytoskeletal regulation and cell motility.
KCTD12
TMSB10
IGFBP3
S100A6
SRGN
3: Inflammatory response.
SOD2
CXCL8
FTL
CD59
CSF3
Freq
Count of Tissues
TSP10
TSP14
TSP15
TSP4
TSP8
TSP9
Normalized Gene Counts
Count of Donors
Eye
Fat
Liver
Lymph_Node
Pancreas
Prostate
Salivary_Gland
Skin
Spleen
Uterus

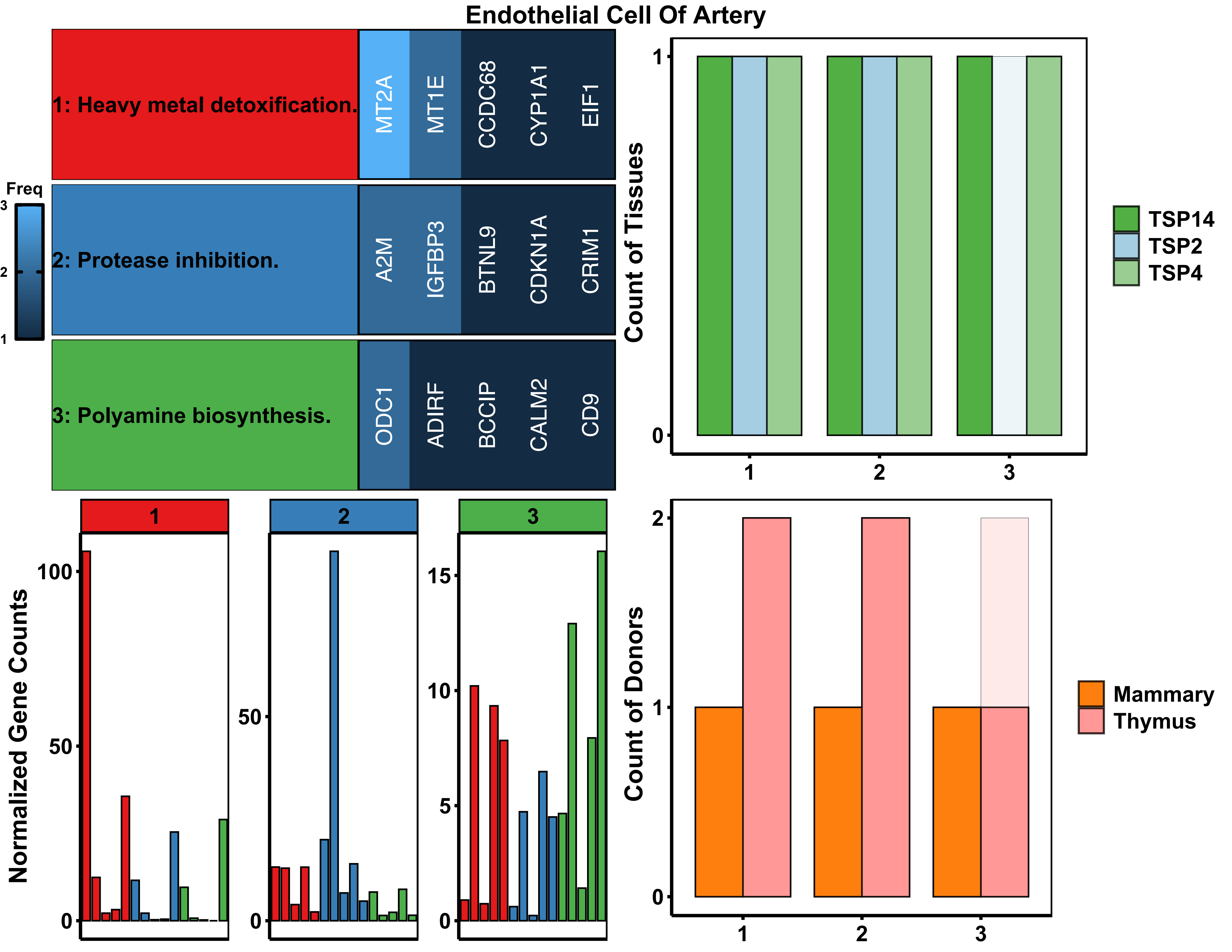

Endothelial Cell Of Artery
1: Heavy metal detoxification.
MT2A
MT1E
CCDC68
CYP1A1
EIF1
2: Protease inhibition.
A2M
IGFBP3
BTNL9
CDKN1A
CRIM1
3: Polyamine biosynthesis.
ODC1
ADIRF
BCCIP
CALM2
CD9
Freq
3
2
1
Count of Tissues
1
0
1
2
3
TSP14
TSP2
TSP4
Normalized Gene Counts
100
50
0
1
2
3
15
10
5
Count of Donors
2
1
0
Mammary
Thymus

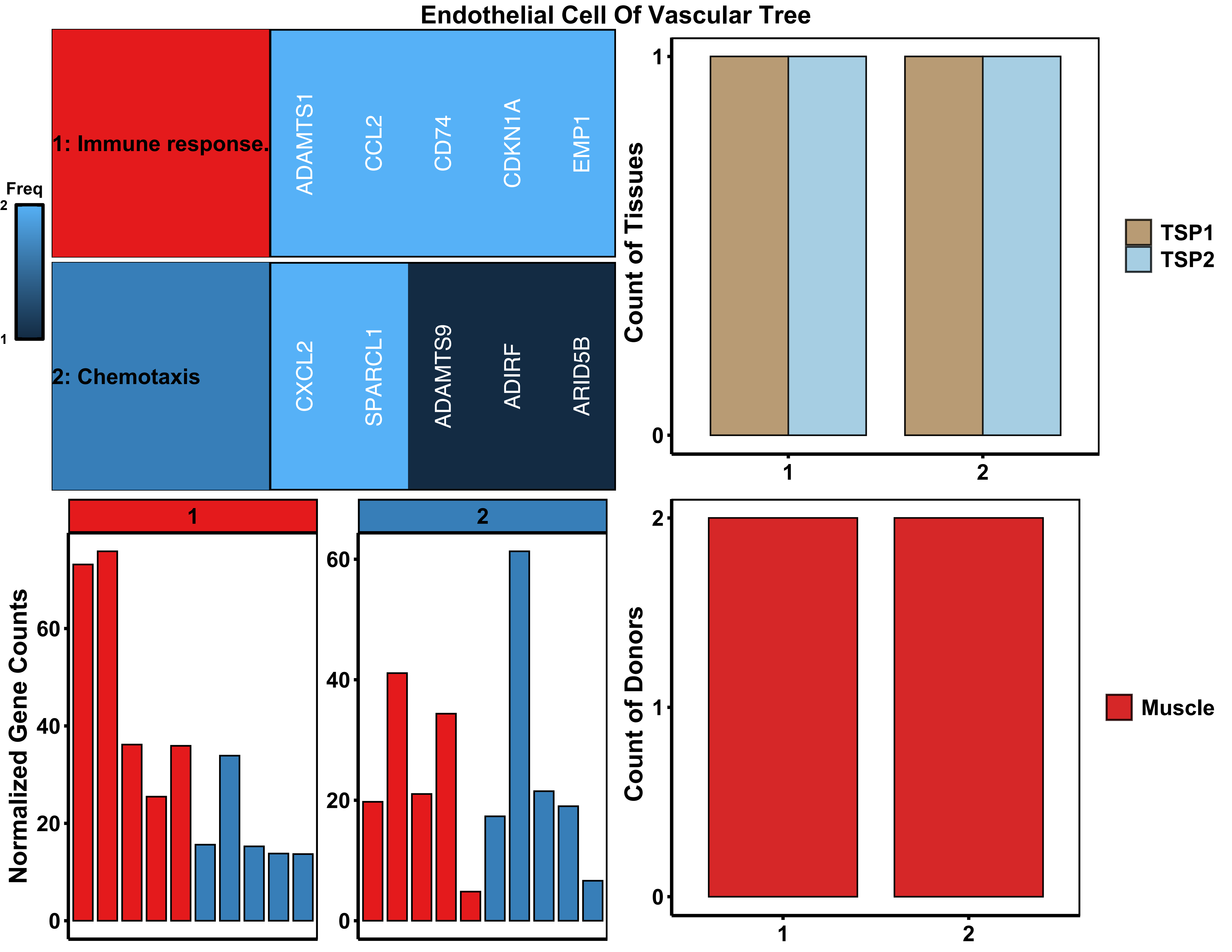
Endothelial Cell Of Vascular Tree
1: Immune response.
ADAMTS1
CCL2
CD74
CDKN1A
EMP1
2: Chemotaxis
CXCL2
SPARCL1
ADAMTS9
ADIRF
ARID5B
Freq
2
1
Count of Tissues
TSP1
TSP2
Normalized Gene Counts
Count of Donors
Muscle
0
20
40
60
1
2

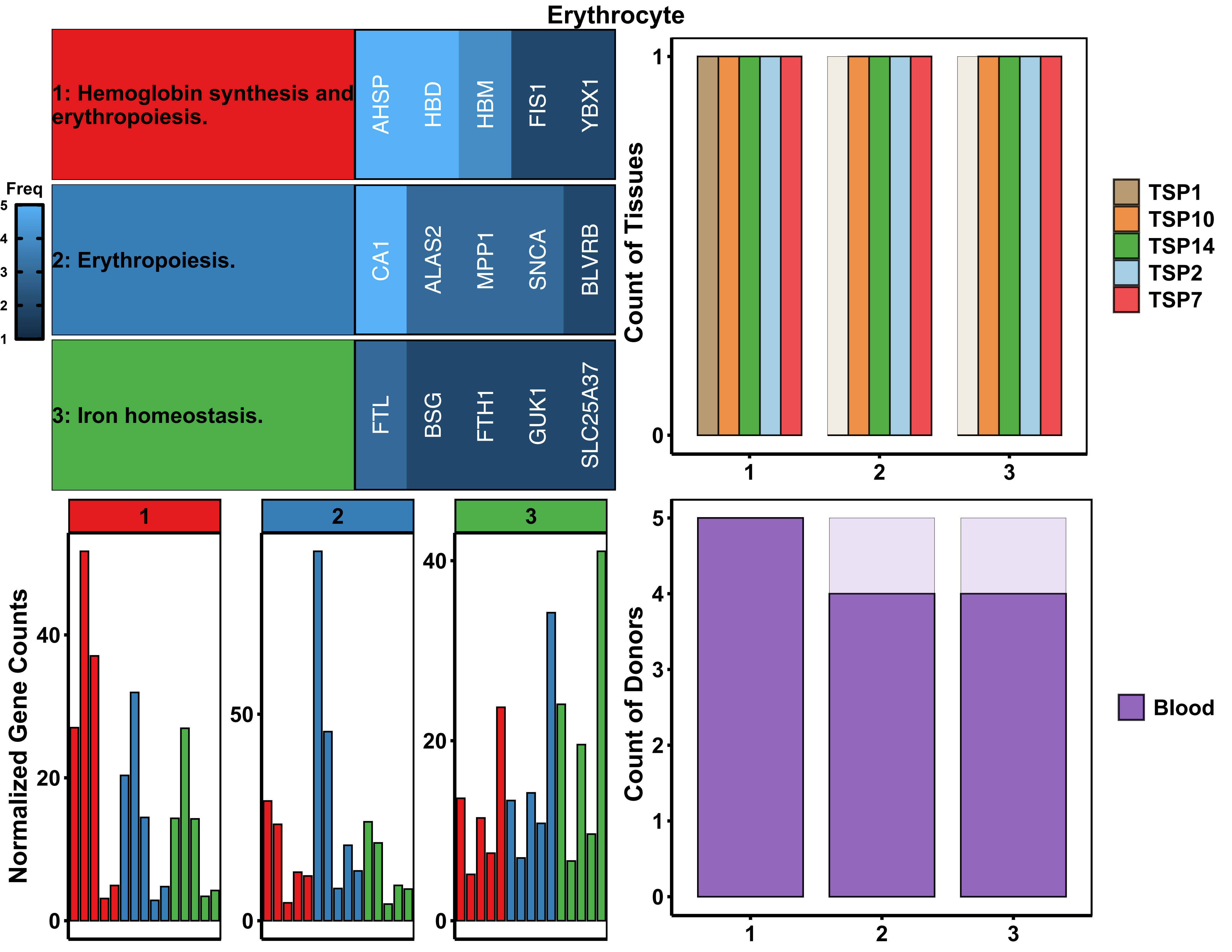

Erythrocyte
1: Hemoglobin synthesis and erythropoiesis.
AHSP
HBD
HBM
FIS1
YBX1
2: Erythropoiesis.
CA1
ALAS2
MPP1
SNCA
BLVRB
3: Iron homeostasis.
FTL
BSG
FTH1
GUK1
SLC25A37
Freq
Count of Tissues
TSP1
TSP10
TSP14
TSP2
TSP7
Normalized Gene Counts
Count of Donors
Blood

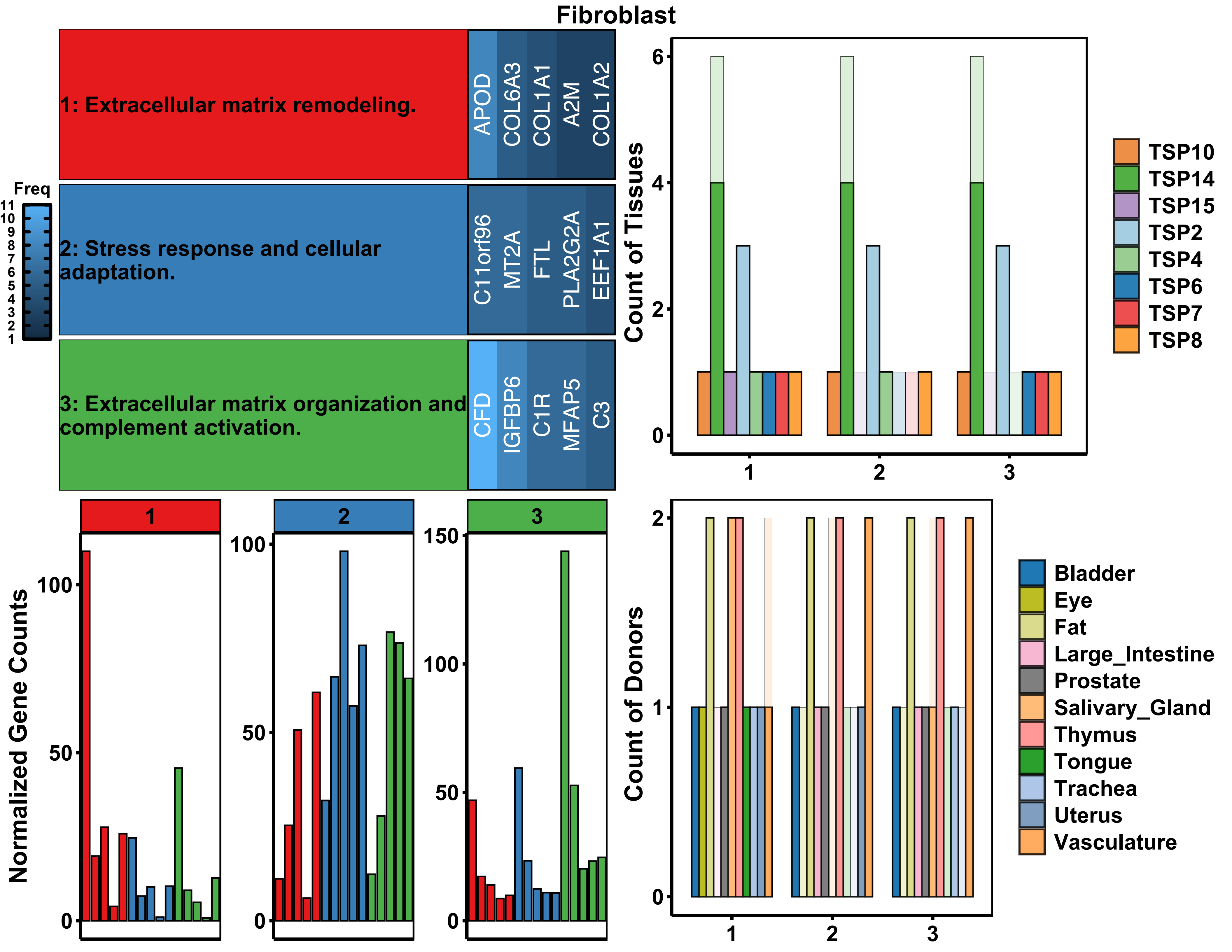
Fibroblast
1: Extracellular matrix remodeling.
APOD
COL6A3
COL1A1
A2M
COL1A2
2: Stress response and cellular adaptation.
C11orf96
MT2A
FTL
PLA2G2A
EEF1A1
3: Extracellular matrix organization and complement activation.
CFD
IGFBP6
C1R
MFAP5
C3
Freq
11
10
9
8
7
6
5
4
3
2
1
Count of Tissues
TSP10
TSP14
TSP15
TSP2
TSP4
TSP6
TSP7
TSP8
Normalized Gene Counts
Count of Donors
Bladder
Eye
Fat
Large_Intestine
Prostate
Salivary_Gland
Thymus
Tongue
Trachea
Uterus
Vasculature

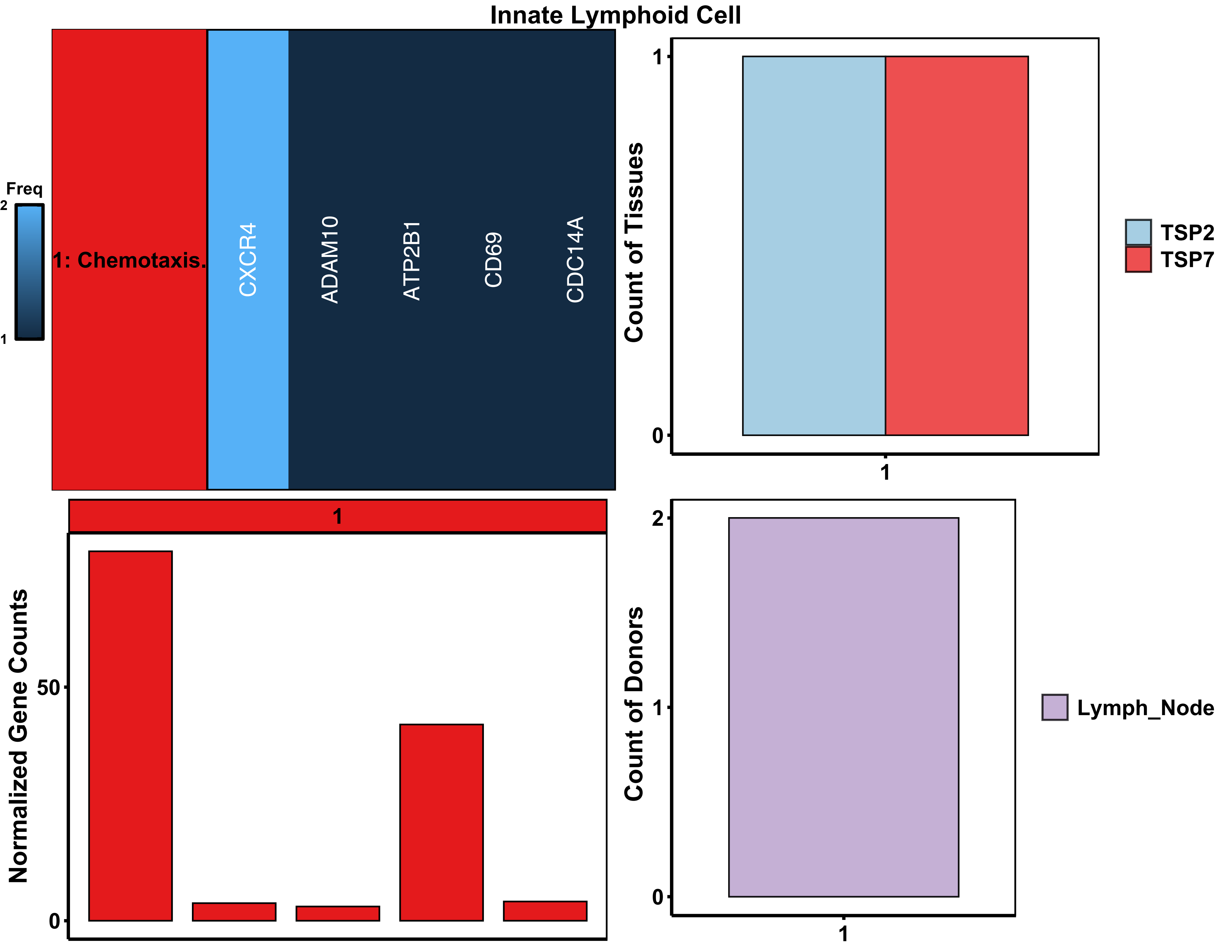

Innate Lymphoid Cell
Freq
2
1
1: Chemotaxis.
CXCR4
ADAM10
ATP2B1
CD69
CDC14A
Count of Tissues
1
0
1
TSP2
TSP7
1
Normalized Gene Counts
50
0
Count of Donors
2
1
0
1
Lymph_Node

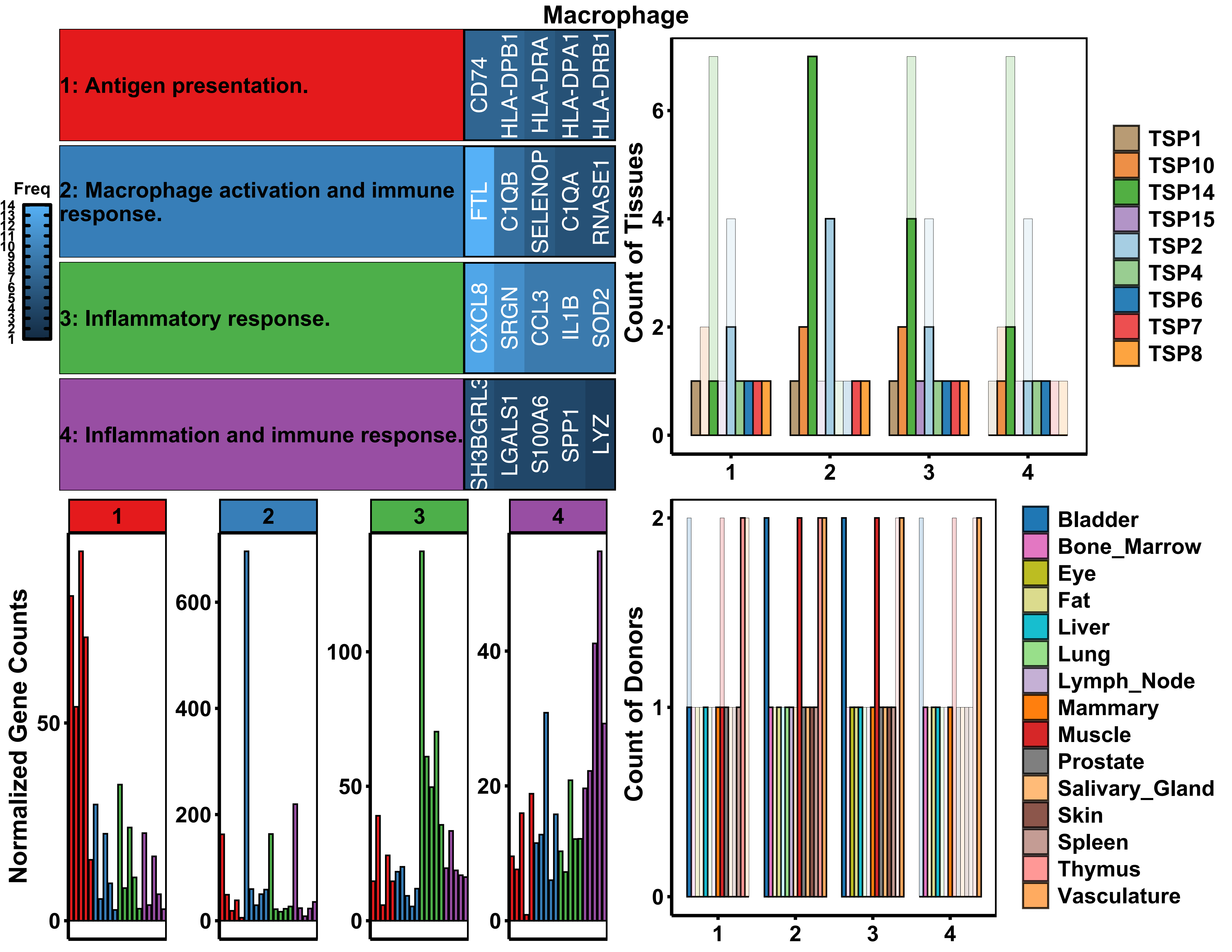

Macrophage
1: Antigen presentation.
CD74
HLA-DPB1
HLA-DRA
HLA-DPA1
HLA-DRB1
2: Macrophage activation and immune response.
FTL
C1QB
SELENOP
C1QA
RNASE1
3: Inflammatory response.
CXCL8
SRGN
CCL3
IL1B
SOD2
4: Inflammation and immune response.
SH3BGRL3
LGALS1
S100A6
SPP1
LYZ
Freq
Count of Tissues
TSP1
TSP10
TSP14
TSP15
TSP2
TSP4
TSP6
TSP7
TSP8
Normalized Gene Counts
Count of Donors
Bladder
Bone_Marrow
Eye
Fat
Liver
Lung
Lymph_Node
Mammary
Muscle
Prostate
Salivary_Gland
Skin
Spleen
Thymus
Vasculature

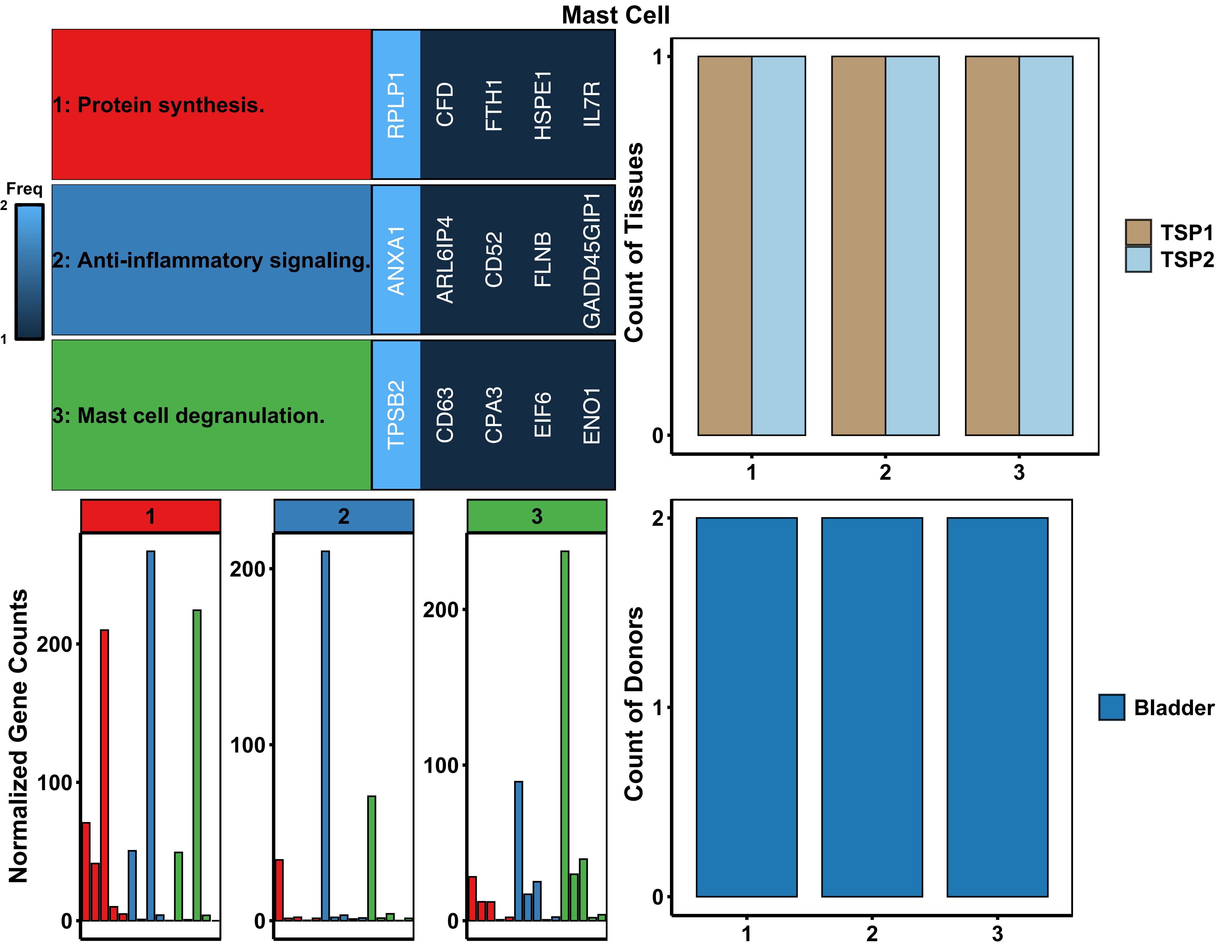
Mast Cell
Freq
2
1
1: Protein synthesis.
RPLP1
CFD
FTH1
HSPE1
IL7R
2: Anti-inflammatory signaling.
ANXA1
ARL6IP4
CD52
FLNB
GADD45GIP1
3: Mast cell degranulation.
TPSB2
CD63
CPA3
EIF6
ENO1
Count of Tissues
TSP1
TSP2
Normalized Gene Counts
Count of Donors
Bladder

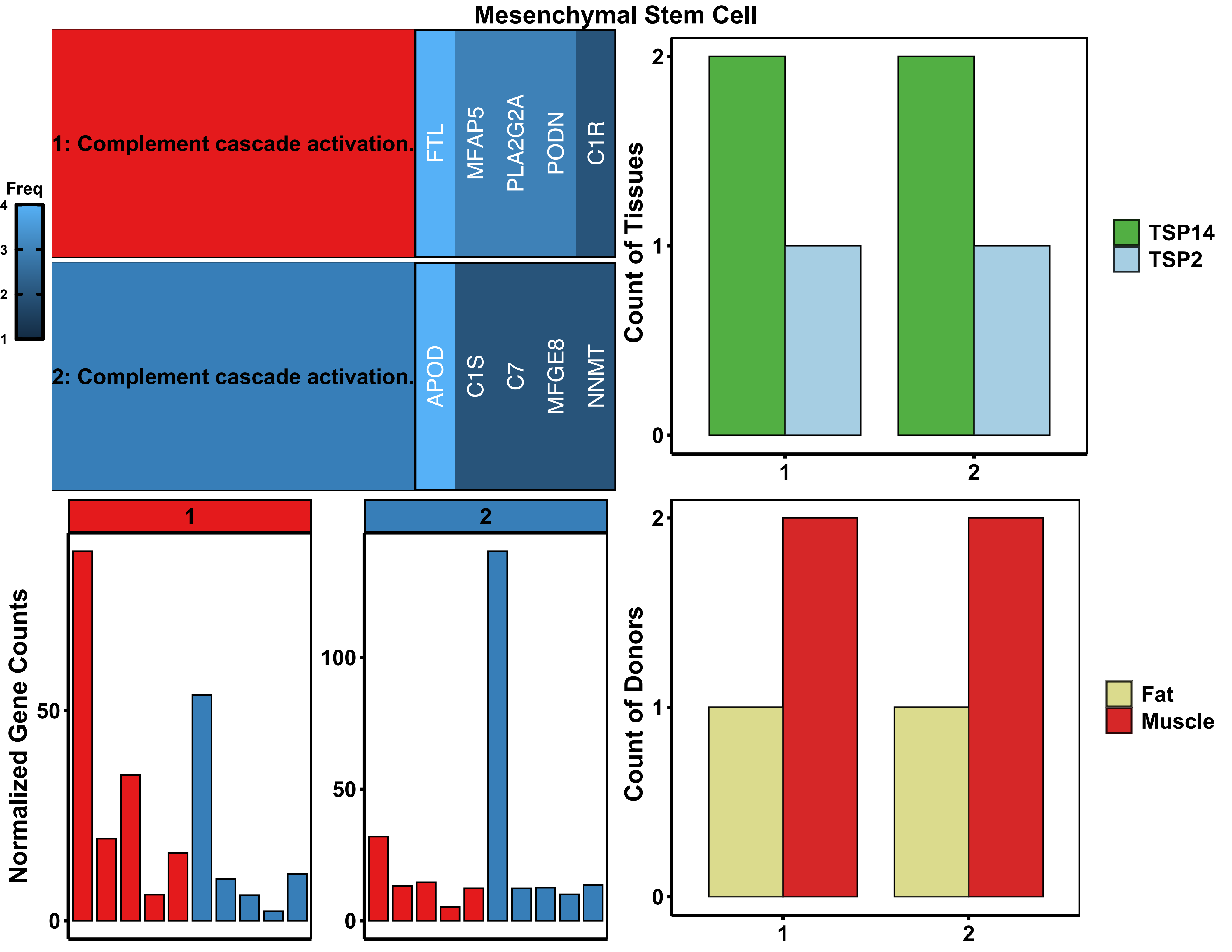

Mesenchymal Stem Cell
Freq
4
3
2
1
1: Complement cascade activation.
FTL
MFAP5
PLA2G2A
PODN
C1R
2: Complement cascade activation.
APOD
C1S
C7
MFGE8
NNMT
Count of Tissues
TSP14
TSP2
Count of Donors
Fat
Muscle
Normalized Gene Counts
1
2
0
50
100

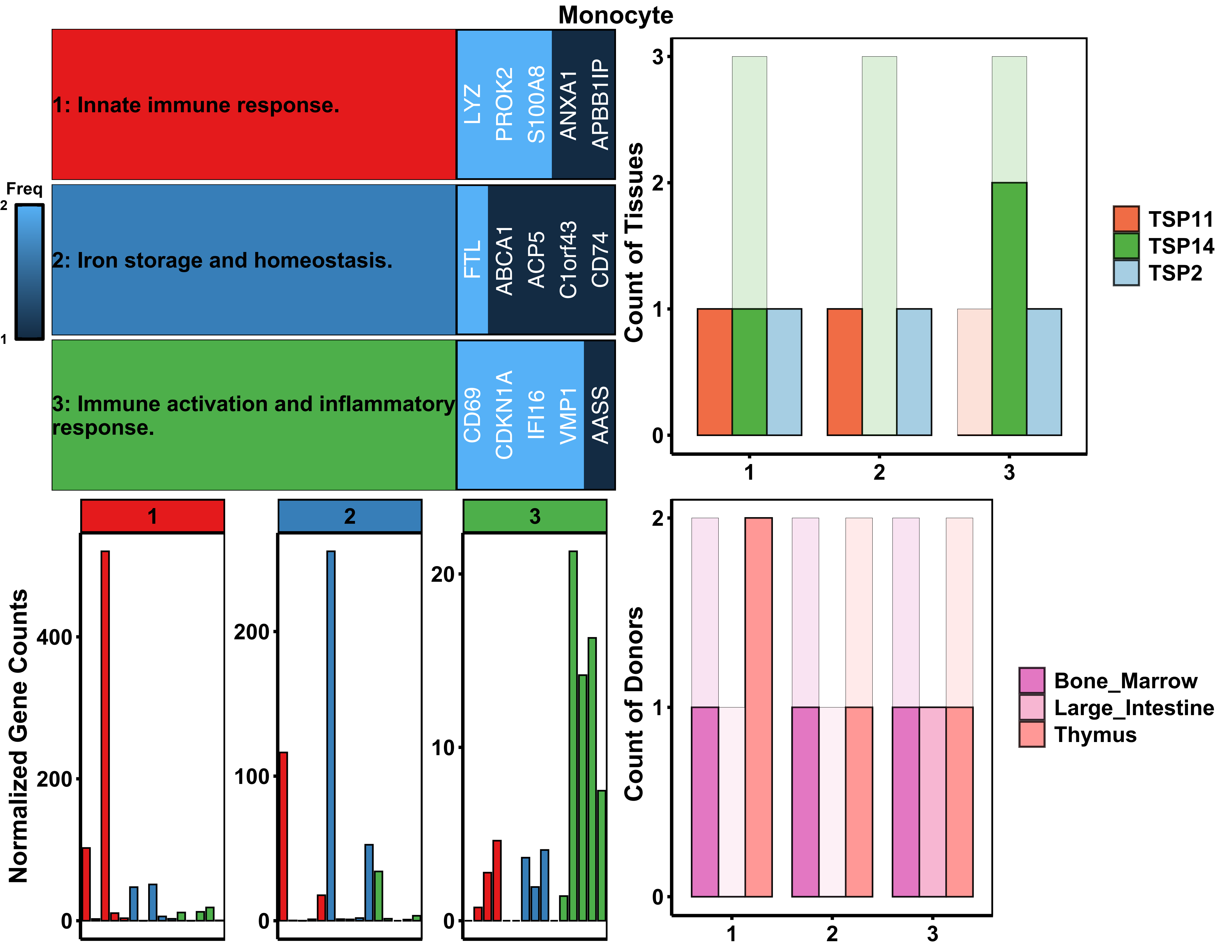

Monocyte
Freq
2
1
1: Innate immune response.
LYZ
PROK2
S100A8
ANXA1
APBB1IP
2: Iron storage and homeostasis.
FTL
ABCA1
ACP5
C1orf43
CD74
3: Immune activation and inflammatory response.
CD69
CDKN1A
IFI16
VMP1
AASS
Count of Tissues
0
1
2
3
TSP11
TSP14
TSP2
Normalized Gene Counts
0
200
400
100
10
20
Count of Donors
Bone_Marrow
Large_Intestine
Thymus

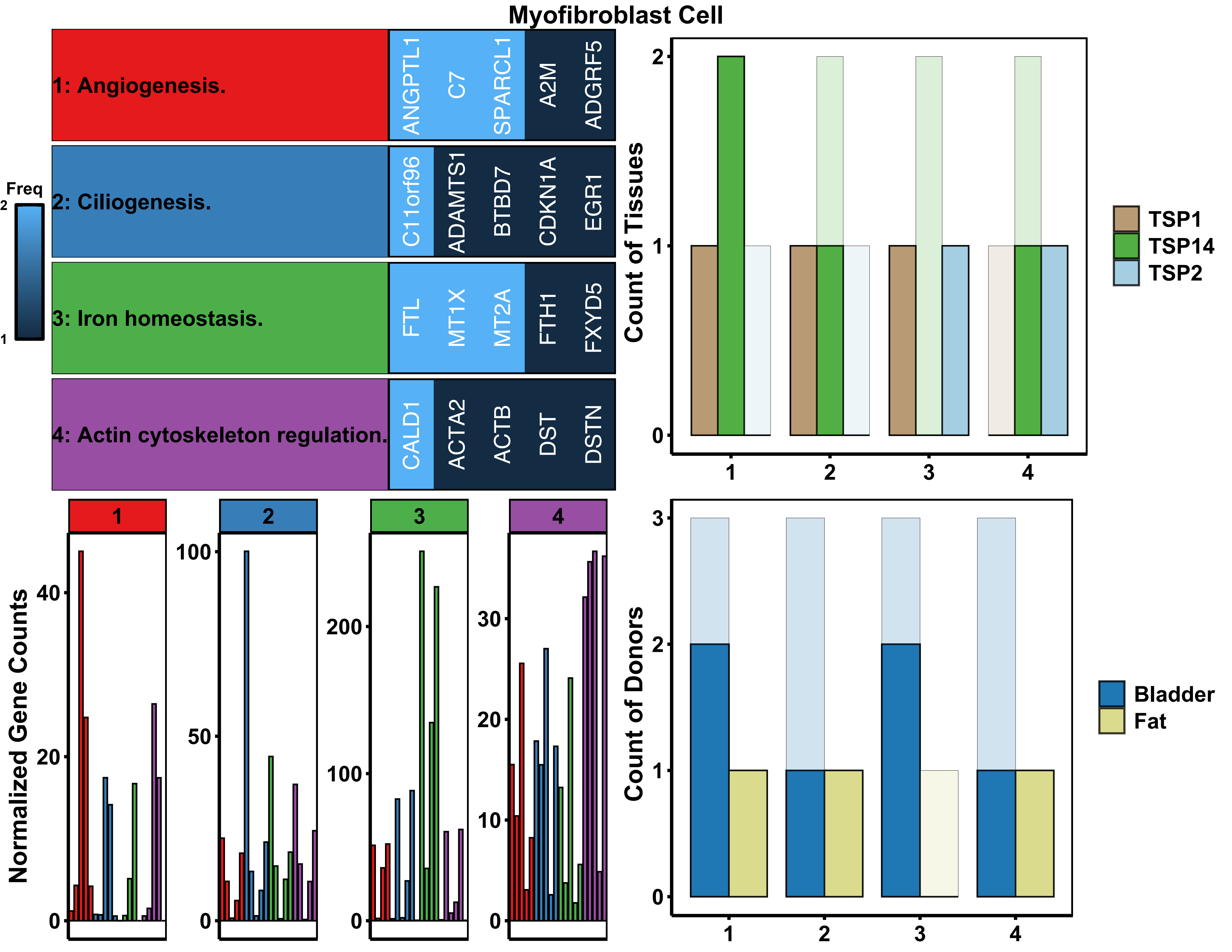
Myofibroblast Cell
1: Angiogenesis.
ANGPTL1
C7
SPARCL1
A2M
ADGRF5
2: Ciliogenesis.
C11orf96
ADAMTS1
BTBD7
CDKN1A
EGR1
3: Iron homeostasis.
FTL
MT1X
MT2A
FTH1
FXYD5
4: Actin cytoskeleton regulation.
CALD1
ACTA2
ACTB
DST
DSTN
Freq
Count of Tissues
TSP1
TSP14
TSP2
Normalized Gene Counts
Count of Donors
Bladder
Fat

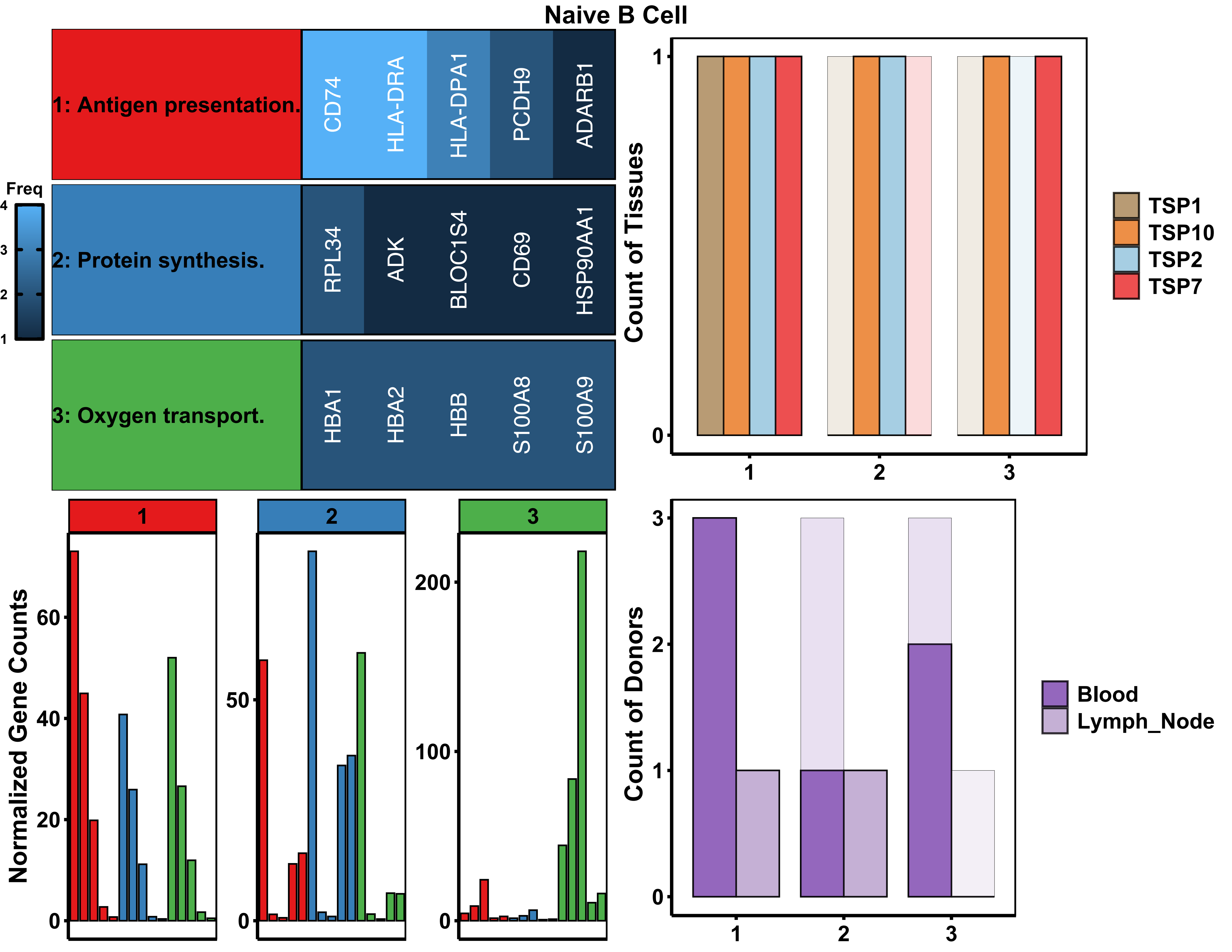

Naive B Cell
1: Antigen presentation.
CD74
HLA-DRA
HLA-DPA1
PCDH9
ADARB1
2: Protein synthesis.
RPL34
ADK
BLOC1S4
CD69
HSP90AA1
3: Oxygen transport.
HBA1
HBA2
HBB
S100A8
S100A9
Freq
4
3
2
1
Count of Tissues
1
0
1
2
3
TSP1
TSP10
TSP2
TSP7
Normalized Gene Counts
1
2
3
60
40
20
0
50
0
200
100
0
Count of Donors
3
2
1
0
1
2
3
Blood
Lymph_Node

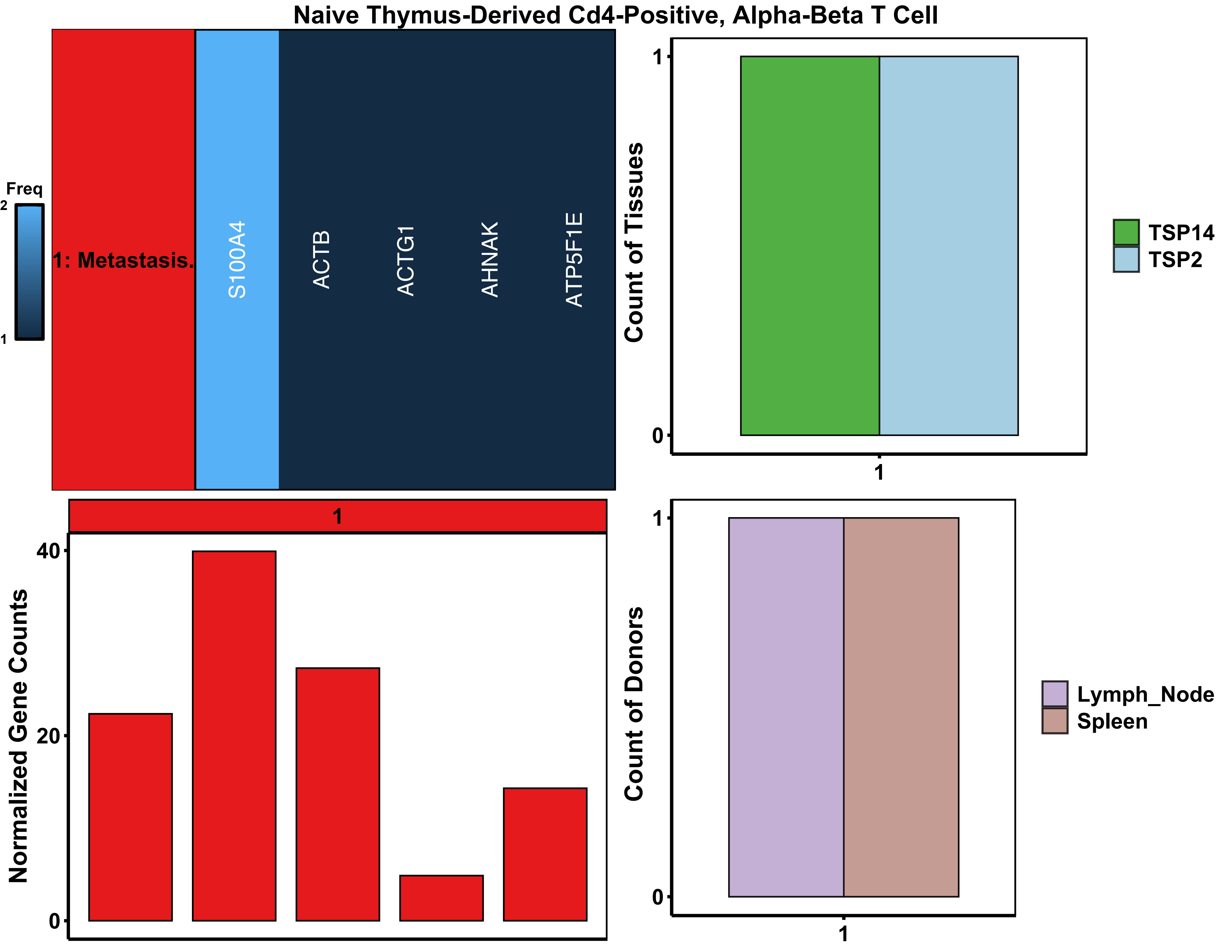
Naive Thymus-Derived Cd4-Positive, Alpha-Beta T Cell
Freq
2
1
1: Metastasis.
S100A4
ACTB
ACTG1
AHNAK
ATP5F1E
Count of Tissues
1
0
1
TSP14
TSP2
1
Normalized Gene Counts
40
20
0
Count of Donors
1
0
1
Lymph_Node
Spleen

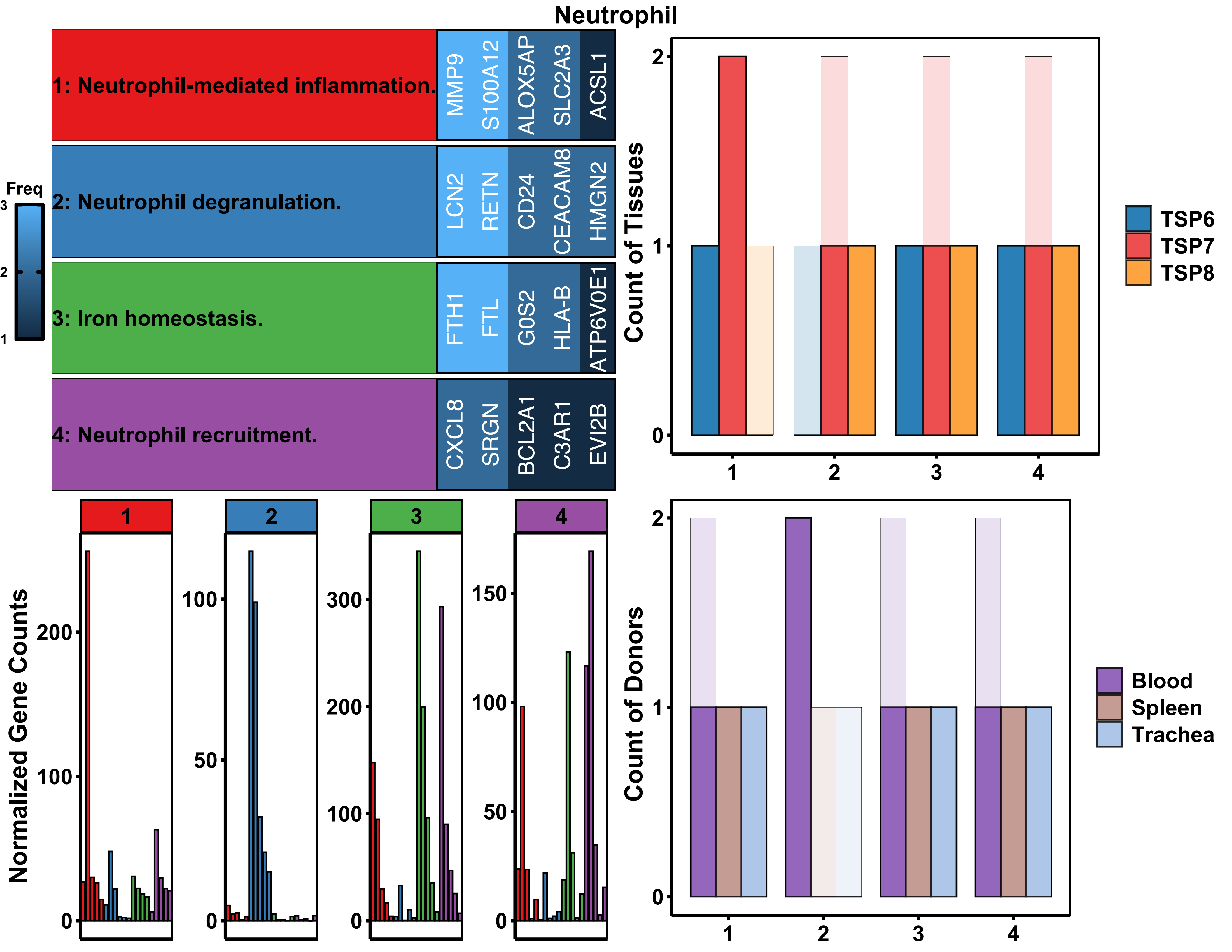

Neutrophil
1: Neutrophil-mediated inflammation.
MMP9
S100A12
ALOX5AP
SLC2A3
ACSL1
2: Neutrophil degranulation.
LCN2
RETN
CD24
CEACAM8
HMGN2
3: Iron homeostasis.
FTH1
FTL
G0S2
HLA-B
ATP6V0E1
4: Neutrophil recruitment.
CXCL8
SRGN
BCL2A1
C3AR1
EVI2B
Freq
3
2
1
Count of Tissues
TSP6
TSP7
TSP8
Normalized Gene Counts
Count of Donors
Blood
Spleen
Trachea

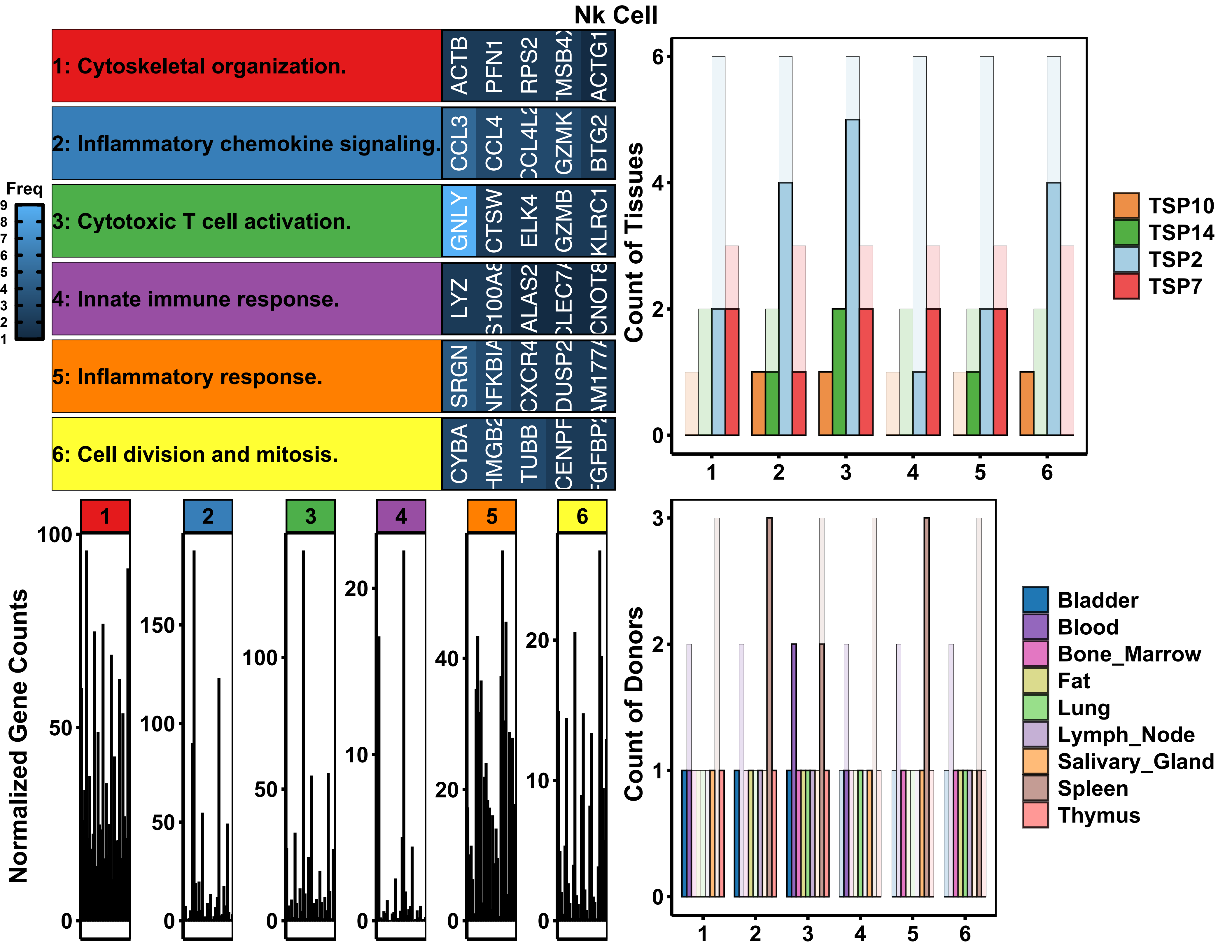

Nk Cell
1: Cytoskeletal organization.
ACTB PFN1 RPS2 TMSB4X ACTG1
2: Inflammatory chemokine signaling.
CCL3 CCL4 CCL4L2 GZMK BTG2
3: Cytotoxic T cell activation.
GNLY CTSW ELK4 GZMB KLRC1
4: Innate immune response.
LYZ ALAS2
5: Inflammatory response.
SRGN CXCR4
6: Cell division and mitosis.
CYBA TUBB
Freq
Count of Tissues
TSP10
TSP14
TSP2
TSP7
Normalized Gene Counts
Count of Donors
Bladder
Blood
Bone_Marrow
Fat
Lung
Lymph_Node
Salivary_Gland
Spleen
Thymus

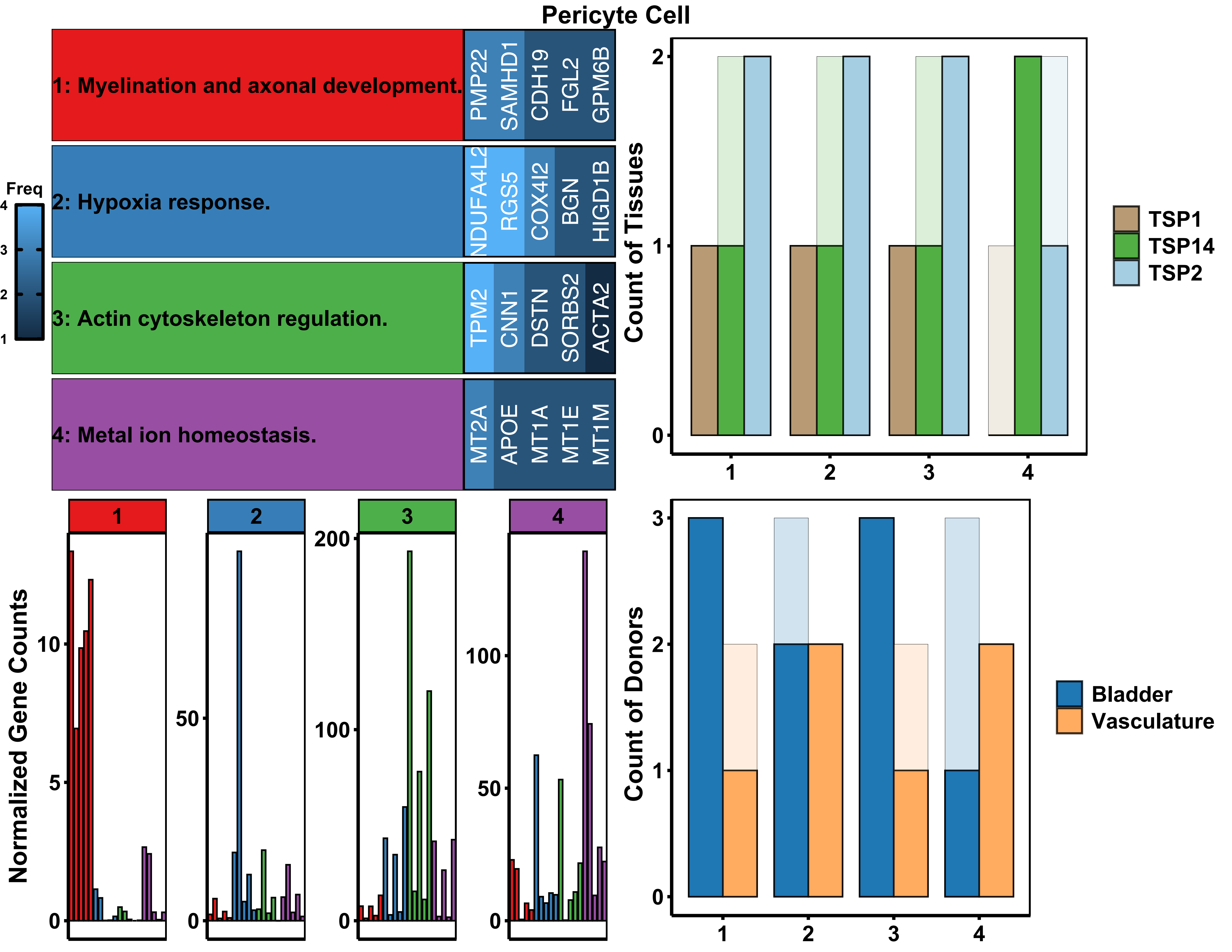

Pericyte Cell
1: Myelination and axonal development.
PMP22
SAMHD1
CDH19
FGL2
GPM6B
2: Hypoxia response.
NDUFA4L2
RGS5
COX4I2
BGN
HIGD1B
3: Actin cytoskeleton regulation.
TPM2
CNN1
DSTN
SORBS2
ACTA2
4: Metal ion homeostasis.
MT2A
APOE
MT1A
MT1E
MT1M
Freq
4
3
2
1
Count of Tissues
2
1
0
1
2
3
4
TSP1
TSP14
TSP2
Normalized Gene Counts
1
2
3
4
10
5
0
50
0
200
100
0
100
50
0
Count of Donors
3
2
1
0
1
2
3
4
Bladder
Vasculature

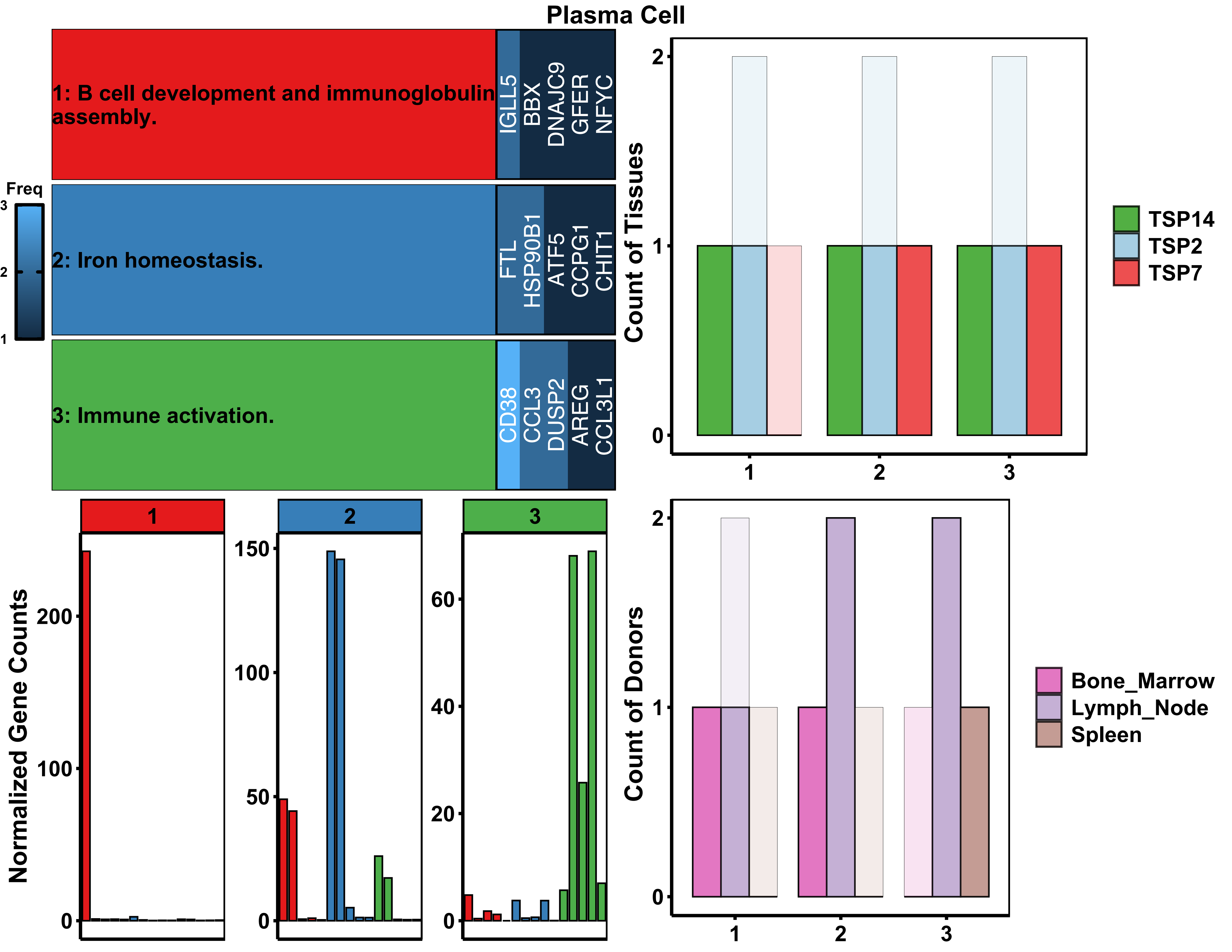
Plasma Cell
Freq
3
2
1
1: B cell development and immunoglobulin assembly.
IGLL5
BBX
DNAJC9
GFER
NFYC
2: Iron homeostasis.
FTL
HSP90B1
ATF5
CCPG1
CHIT1
3: Immune activation.
CD38
CCL3
DUSP2
AREG
CCL3L1
Count of Tissues
TSP14
TSP2
TSP7
Normalized Gene Counts
Count of Donors
Bone_Marrow
Lymph_Node
Spleen

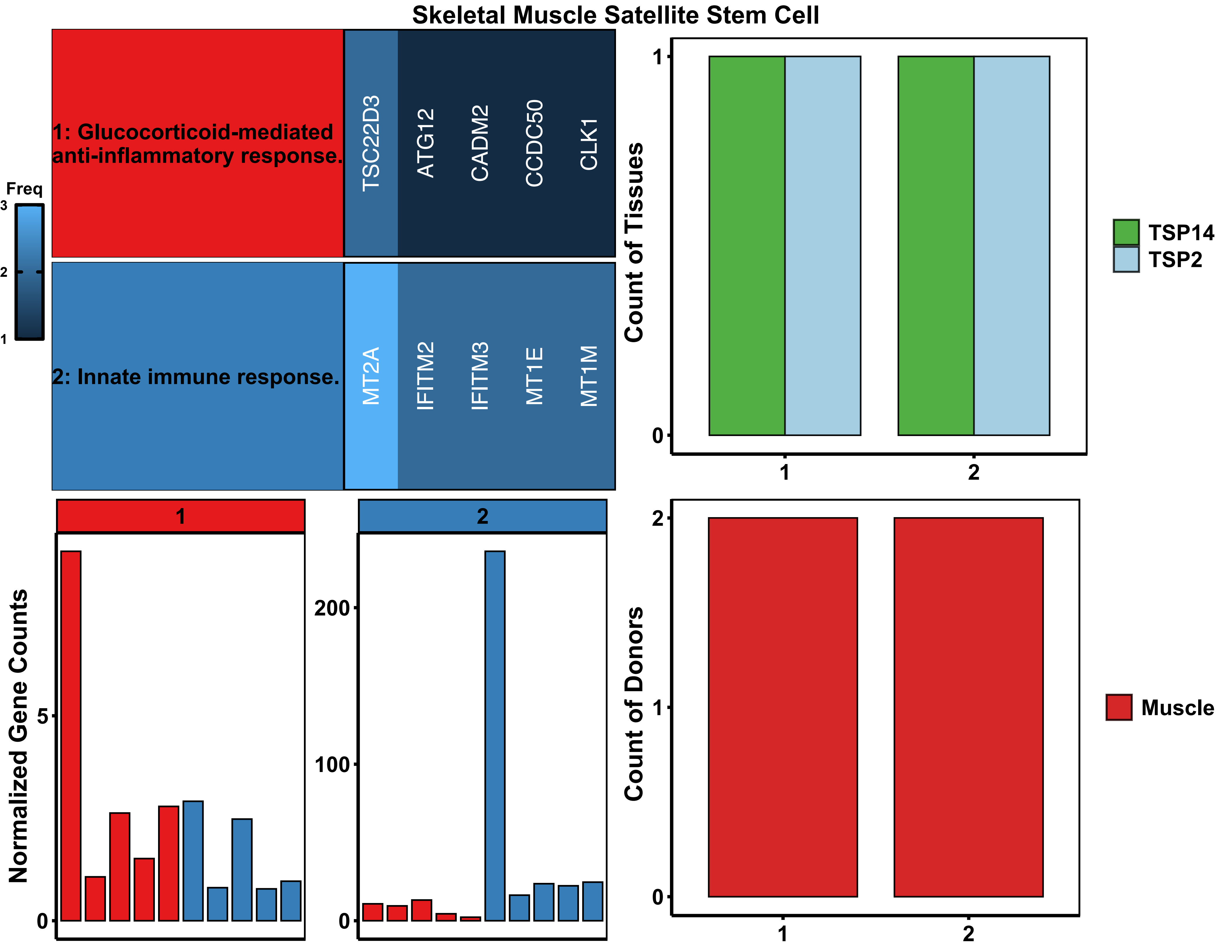

Skeletal Muscle Satellite Stem Cell
Freq
3
2
1
1: Glucocorticoid-mediated anti-inflammatory response.
TSC22D3
ATG12
CADM2
CCDC50
CLK1
2: Innate immune response.
MT2A
IFITM2
IFITM3
MT1E
MT1M
Count of Tissues
TSP14
TSP2
Normalized Gene Counts
200
100
5
0
Count of Donors
Muscle

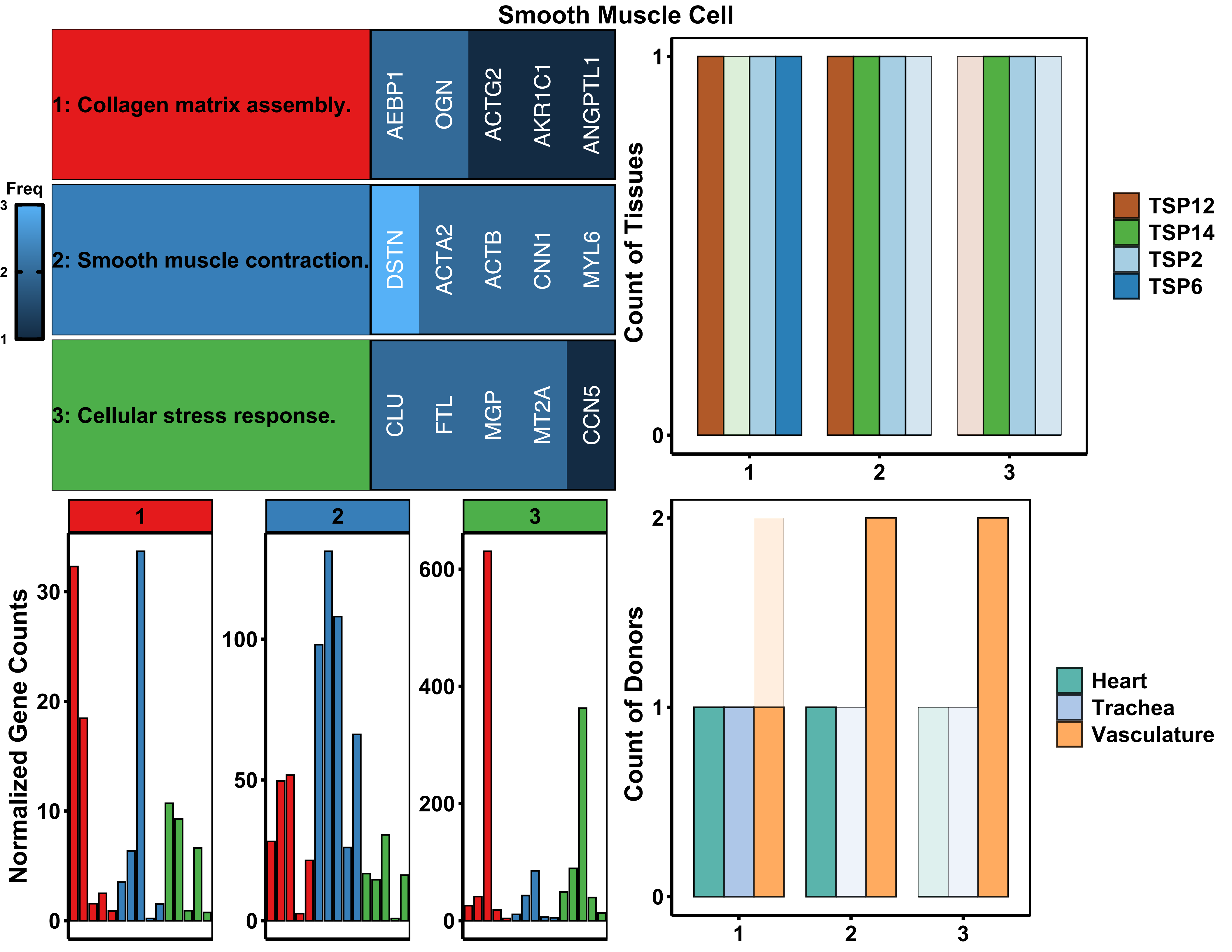

Smooth Muscle Cell
1: Collagen matrix assembly.
AEBP1
OGN
ACTG2
AKR1C1
ANGPTL1
2: Smooth muscle contraction.
DSTN
ACTA2
ACTB
CNN1
MYL6
3: Cellular stress response.
CLU
FTL
MGP
MT2A
CCN5
Freq
3
2
1
Count of Tissues
TSP12
TSP14
TSP2
TSP6
Normalized Gene Counts
Count of Donors
Heart
Trachea
Vasculature

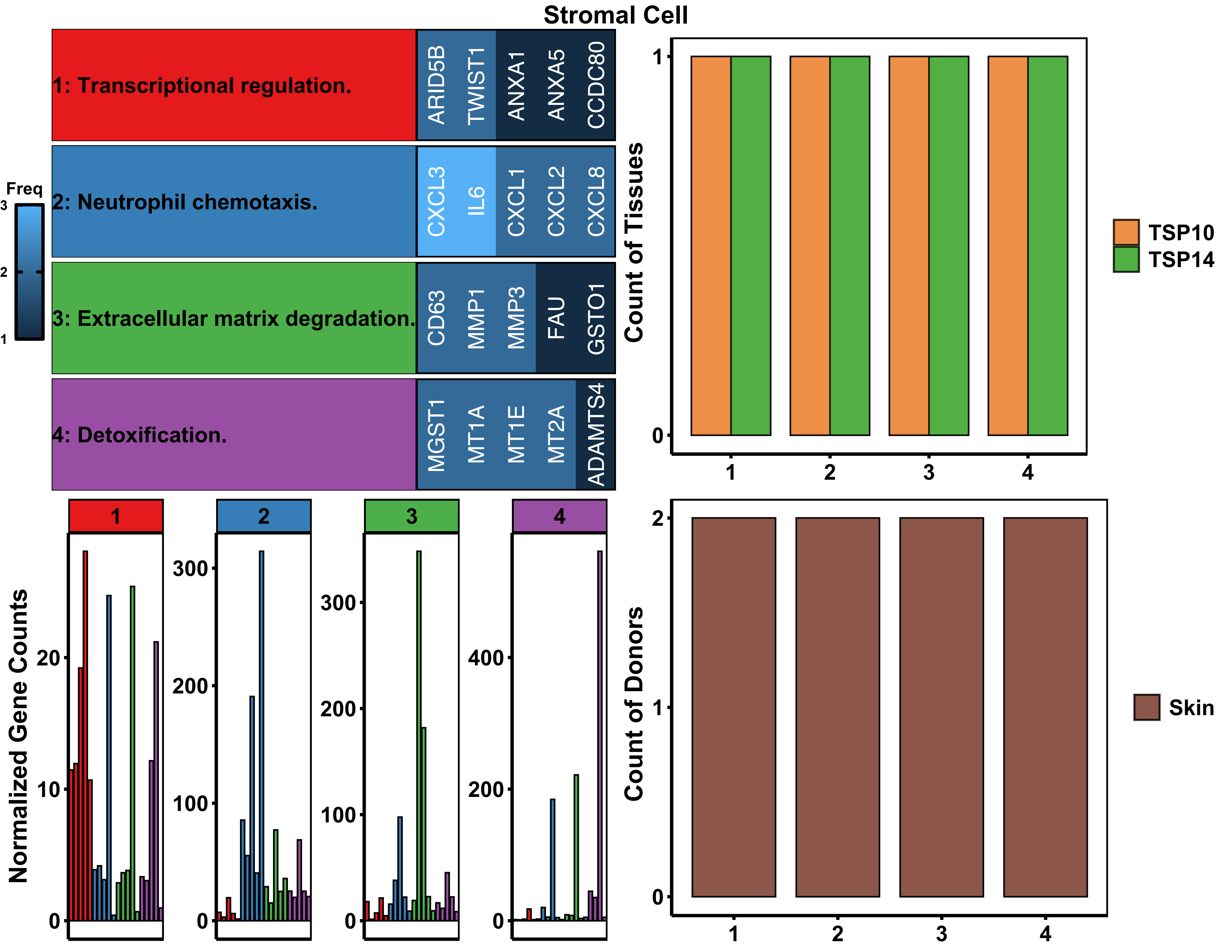

Stromal Cell
1: Transcriptional regulation.
ARID5B
TWIST1
ANXA1
ANXA5
CCDC80
2: Neutrophil chemotaxis.
CXCL3
IL6
CXCL1
CXCL2
CXCL8
3: Extracellular matrix degradation.
CD63
MMP1
MMP3
FAU
GSTO1
4: Detoxification.
MGST1
MT1A
MT1E
MT2A
ADAMTS4
Freq
3
2
1
Count of Tissues
TSP10
TSP14
Normalized Gene Counts
1
2
3
4
Count of Donors
Skin

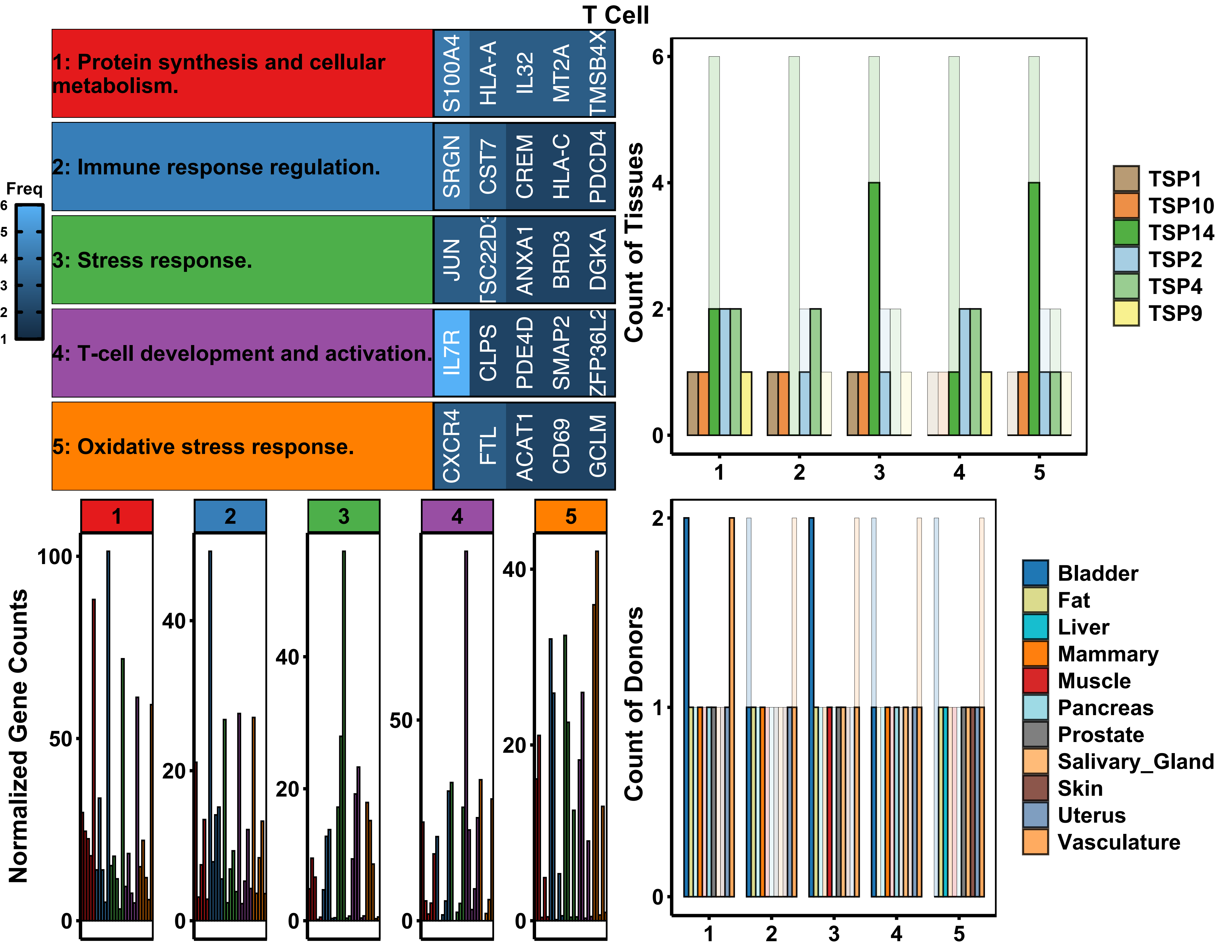

T Cell
1: Protein synthesis and cellular metabolism.
2: Immune response regulation.
3: Stress response.
4: T-cell development and activation.
5: Oxidative stress response.
S100A4
HLA-A
IL32
MT2A
TMSB4X
SRGN
CST7
CREM
HLA-C
PDCD4
JUN
TSC22D3
ANXA1
BRD3
DGKA
IL7R
CLPS
PDE4D
SMAP2
ZFP36L2
CXCR4
FTL
ACAT1
CD69
GCLM
Freq
Count of Tissues
TSP1
TSP10
TSP14
TSP2
TSP4
TSP9
Normalized Gene Counts
Count of Donors
Bladder
Fat
Liver
Mammary
Muscle
Pancreas
Prostate
Salivary_Gland
Skin
Uterus
Vasculature

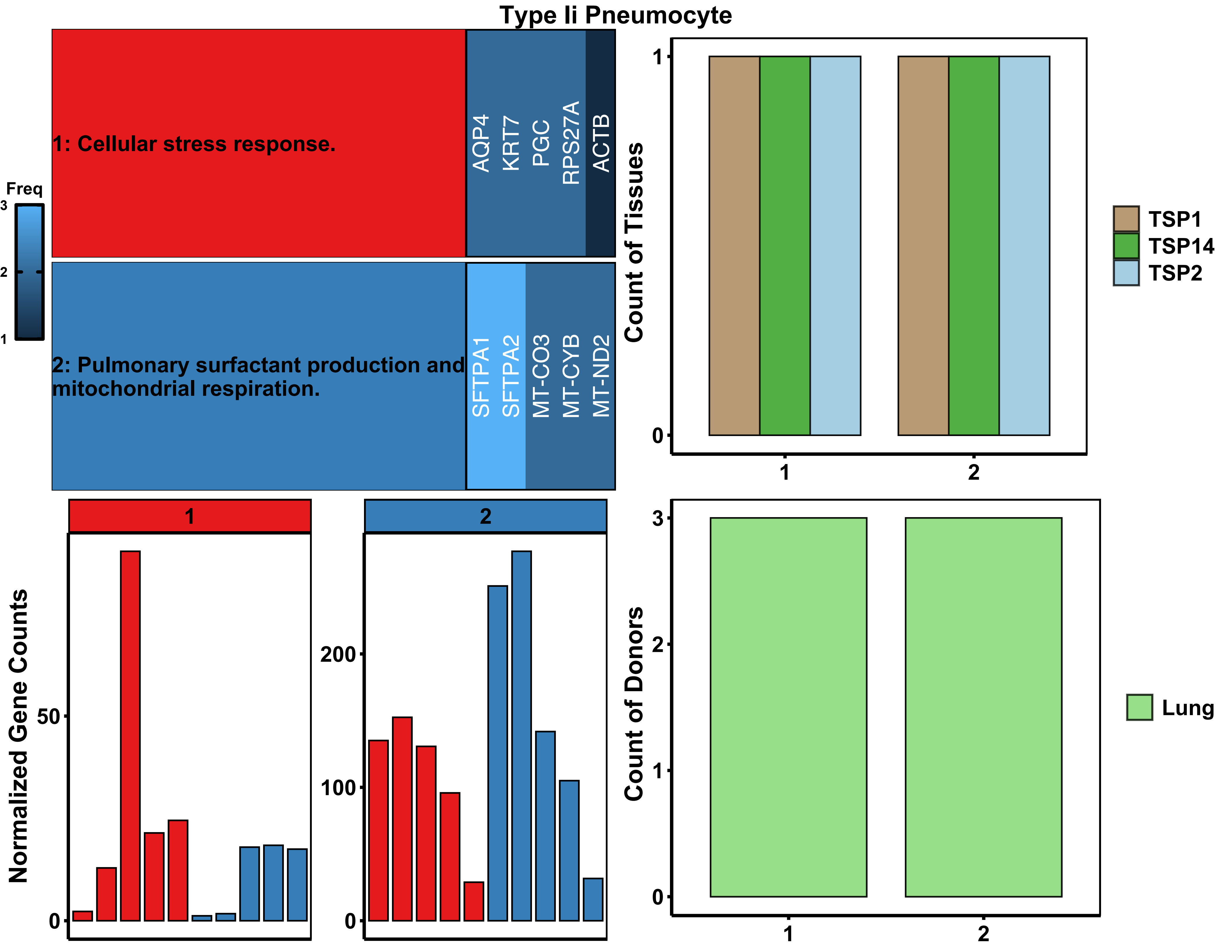
Type Ii Pneumocyte
1: Cellular stress response.
AQP4
KRT7
PGC
RPS27A
ACTB
2: Pulmonary surfactant production and mitochondrial respiration.
SFTPA1
SFTPA2
MT-CO3
MT-CYB
MT-ND2
Freq
3
2
1
Count of Tissues
1
0
1
2
TSP1
TSP14
TSP2
1
2
Normalized Gene Counts
50
0
200
100
0
Count of Donors
3
2
1
0
1
2
Lung

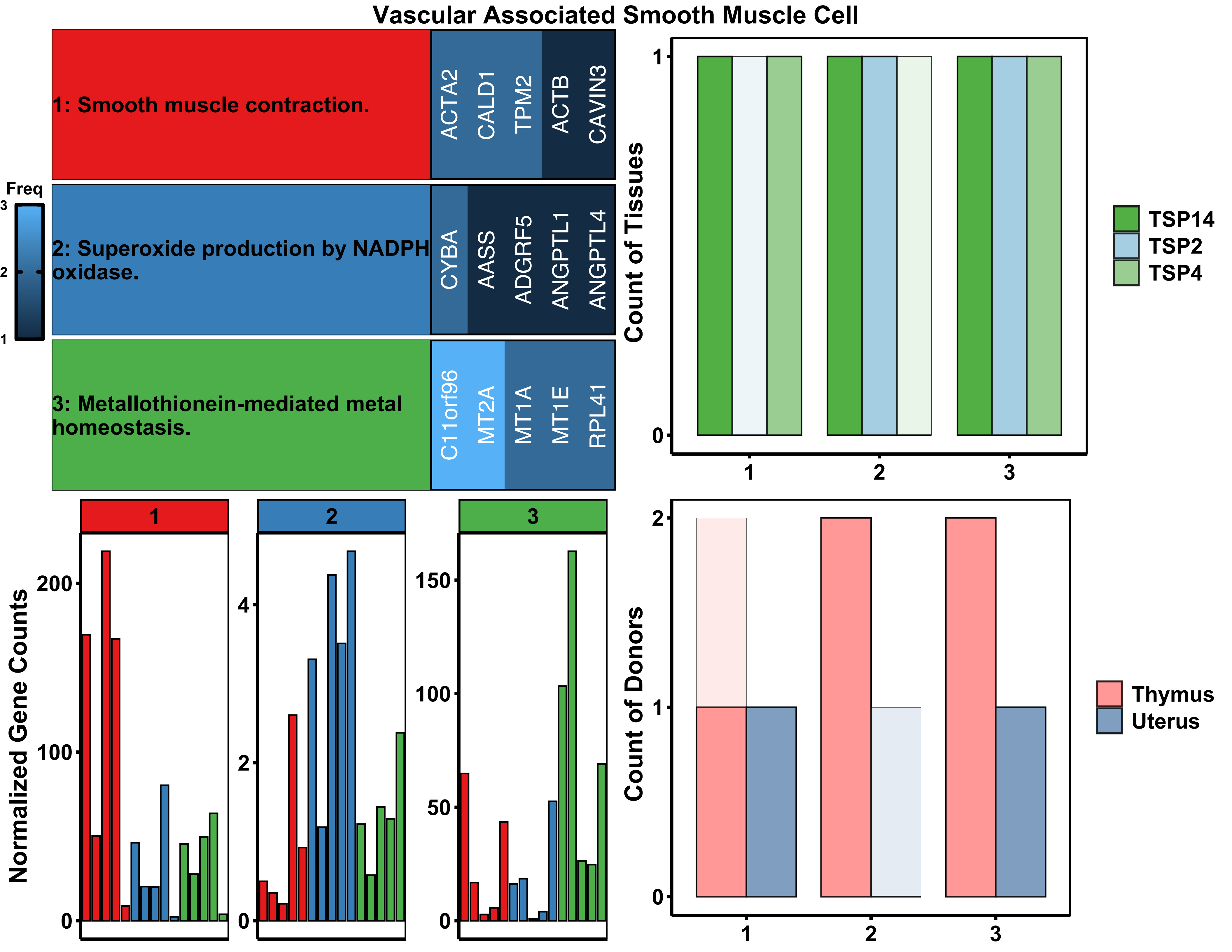

Vascular Associated Smooth Muscle Cell
1: Smooth muscle contraction.
ACTA2
CALD1
TPM2
ACTB
CAVIN3
2: Superoxide production by NADPH oxidase.
CYBA
AASS
ADGRF5
ANGPTL1
ANGPTL4
3: Metallothionein-mediated metal homeostasis.
C11orf96
MT2A
MT1A
MT1E
RPL41
Freq
Count of Tissues
TSP14
TSP2
TSP4
Normalized Gene Counts
Count of Donors
Thymus
Uterus

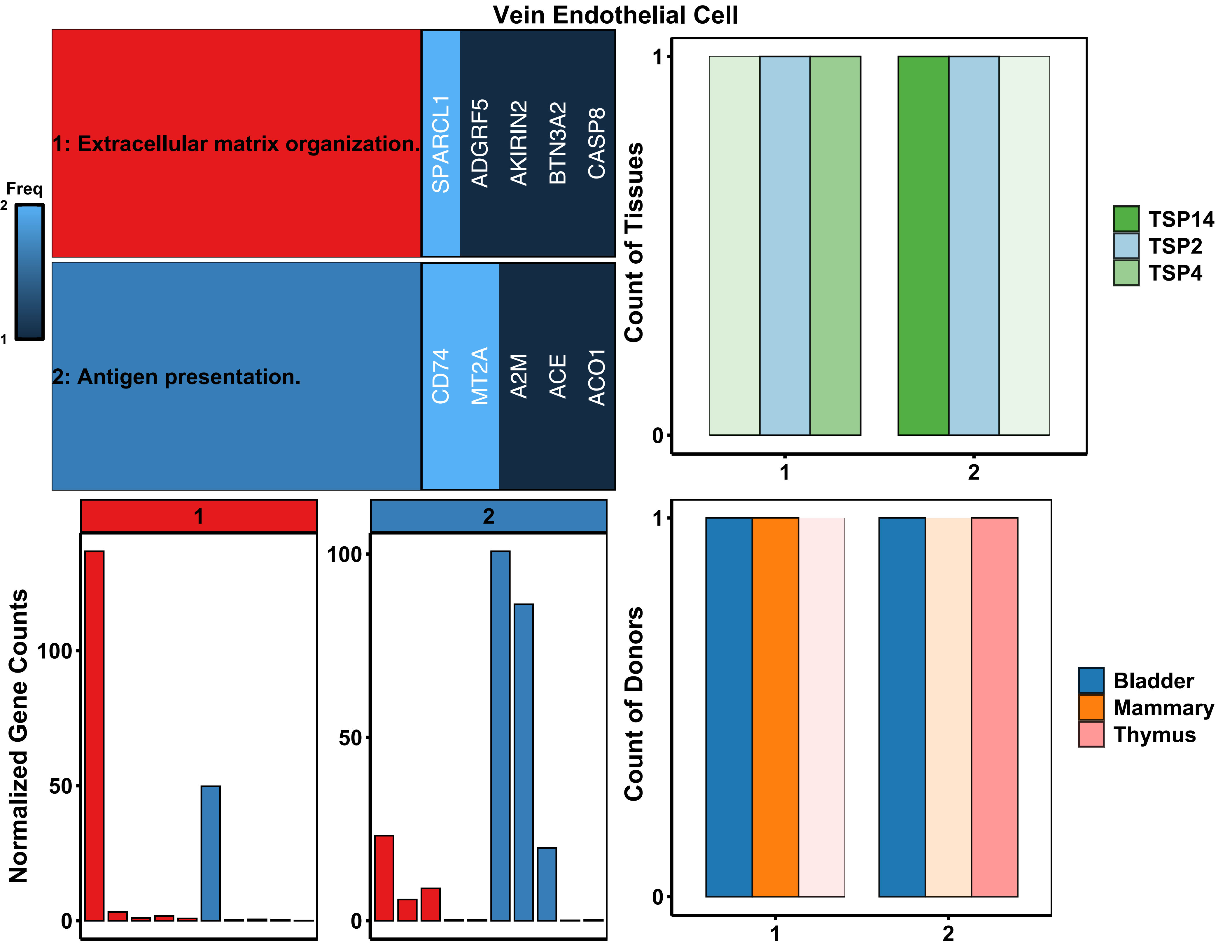

Vein Endothelial Cell
Freq
2
1
1: Extracellular matrix organization.
SPARCL1
ADGRF5
AKIRIN2
BTN3A2
CASP8
2: Antigen presentation.
CD74
MT2A
A2M
ACE
ACO1
Count of Tissues
TSP14
TSP2
TSP4
Normalized Gene Counts
Count of Donors
Bladder
Mammary
Thymus
